# Connectome-based prediction of functional impairment in experimental stroke models

**DOI:** 10.1101/2023.05.05.539601

**Authors:** Oliver Schmitt, Peter Eipert, Yonggang Wang, Atsushi Kanoke, Gratianne Rabiller, Jialing Liu

**Affiliations:** Medical School Hamburg - University of Applied Sciences, Department of Anatomy; University of Rostock, Institute of Anatomy; Department of Neurological Surgery, UCSF; SFVAMC, 1700 Owens Street, San Francisco, CA 94158; Department of Neurological Surgery, Beijing Tiantan Hospital, Capital Medical University, Beijing, PR China, 100050; Department of Neurosurgery, Tohoku University Graduate School of Medicine^3^, 1-1 Seiryo-machi, Aoba-ku, Sendai 980-8574, Japan

**Keywords:** neuroVIISAS, connectomics, network analysis, leaky integrate-and-fire model, FitzHugh-Nagumo model, Wilson-Cowan model, Mimura-Murray reaction-diffusion model, distal middle cerebral artery occlusion model, intracebral hemorrhage model, middle cerebral artery intraluminal suture occlusion model, Barnes maze test, Catwalk test, rotarod, ladder test.

## Abstract

Experimental rat models of stroke and hemorrhage are important tools to investigate cerebrovascular disease pathophysiology mechanisms, yet how significant patterns of functional impairment induced in various models of stroke are related to changes in connectivity at the level of neuronal populations and mesoscopic parcellations of rat brains remain unresolved. To address this gap in knowledge, we employed two middle cerebral artery occlusion models and one intracerebral hemorrhage model with variant extent and location of neuronal dysfunction. Motor and spatial memory function was assessed and the level of hippocampal activation via Fos immunohistochemistry. Contribution of connectivity change to functional impairment was analyzed for connection similarities, graph distances and spatial distances as well as the importance of regions in terms of network architecture based on the *neuroVIISAS* rat connectome. We found that functional impairment correlated with not only the extent but also the locations of the injury among the models. In addition, via coactivation analysis in dynamic rat brain models, we found that lesioned regions led to stronger coactivations with motor function and spatial learning regions than with other unaffected regions of the connectome. Dynamic modeling with the weighted bilateral connectome detected changes in signal propagation in the remote hippocampus in all 3 stroke types, predicting the extent of hippocampal hypoactivation and impairment in spatial learning and memory function. Our study provides a comprehensive analytical framework in predictive identification of remote regions not directly altered by stroke events and their functional implication.

## Introduction

The functional sequelae of stroke vary by nature and degree, depending on the location and extent of brain damage. Although it is relatively intuitive to predict the type and severity of the functional impairment based on brain regions directly impacted by stroke, there is considerable uncertainty regarding how function is affected by the change of brain connectivity due to stroke, particularly in regions remote from the stroke epicenter that still maintain normal structure yet with disrupted functional networks.

Connectomics [156, 158, 172, 178] has been used as a method to investigate and quantify connectivity changes following experimental lesions, injuries and therapies of mammalian brains [7, 107, 166, 172, 180, 213]. Models of neural networks, neural local circuits can be structurally characterized with connectome analyses, while dynamic bioelectrical activity can be modeled in simulated oscillations combining information from neuronal networks and connectomes. By means of a mesoscale connectome analysis using the *neuroVIISAS* platform [157], we previously found that all the damaged cortical regions caused by distal occlusion of the MCA (dMCAO) had a relatively high connectivity with brain regions involved in processing spatial information [156], which may underlie the observed memory impairment [155, 197]. Ample electrophysiological evidence supports the impact of stroke in the remote hippocampus region including changes in hippocampal theta power [68], coherence of theta oscillation between the hippocampus and prefrontal cortex [73, 100], the aberrant increase in sharp-wave-associated ripples and altered theta-gamma modulation [69, 88], reaffirming stroke-induced connectivity changes. By adapting relevant parameter spaces and synaptic connectivity, the neuronal point and population models developed recently exhibited similar dynamic bioelectrical behavior as the recorded data. These simulated oscillations based on mathematical and biophysical principles can be created based on structural connectivity data to provide a reliable prediction model for network activity and behavior. This type of theoretical analysis and computational models was used to explore the dynamics of neuronal networks [25] and [23] and to investigate the propagation of synchronous spiking activity in feedforward neural networks in the absence of electrophysiology data, providing valuable insights into the mechanisms of spiking activity propagation in neuronal networks [2, 120]. Thus, connectomic analysis may help to reveal the functional impairment in greater levels beyond the direct contribution of known lesions caused by experimental brain injury paradigm.

By comparing three experimental models of stroke commonly used in the research community, namely the distal middle cerebral artery occlusion (dMCAO), intraluminal filament model of MCAO and the intracerebral hemorrhage model (ICH) targeting the internal capsule [17, 85, 92, 110, 116], the current study sought to test the hypothesis that lesion location in each stroke model distinctly affects network structure, network dynamics and function [21, 77, 199]. We first compared lesions size and functional impairment with motor and memory tests, and accessed the extent of neural activation via Fos immunohistochemistry in the remote hippocampus region upon spatial exploration [54, 90, 191, 192]. Following the mapping of lesioned regions in each model, by means of connectomics we analyzed the global network structure of control connectome and stroke-induced connectome, local network parameters of control regions following removal of lesioned regions, connectivity relationship between lesioned regions and regions that process motor or learning behavior [197]. We evaluated the differential coactivation of regions in non-lesioned and lesioned connectomes via differential functional connectomics to determine if coactivation changes in functional models may predict motor or learning behavior changes following stroke. We investigated the propagation of spiking activity in remote brain regions relevant to learning and memory function with simulated oscillations using the point neuron FitzHuge-Nagumo (FHN) model, Wilson-Cowan neural mass model and the Mimura-Murray reaction-diffusion model comparing the control and the three lesioned connectomes [2, 120] [25] and [23]. We present our findings in how lesion location affects structural and functional connectivity across the experimental stroke models.

## Materials and Methods

All animal experiments were conducted in accordance with the Guide for Care and Use of Laboratory Animals, and reported in compliance with the Animals in Research: Reporting In Vivo Experiments (ARRIVE) guidelines [44, 99], and were approved by the San Francisco Veterans Affairs Medical Center Institutional Animal Care and Use Committee. The identity of each animal with respect to treatment was concealed to experimenters who conducted the procedures and analysis.

### Models of ischemic and hemorrhagic stroke

#### The distal middle cerebral artery occlusion model (dMCAO)

Stroke was induced unilaterally in male Sprague-Dawley rats (n=21, 2.5 months of age, Charles River, CA) under isoflurane/O_2_/N_2_O (1.5/30/68.5%) according to the well-established distal middle cerebral artery occlusion (dMCAO) method [5, 110, 142, 155, 156, 183]. Briefly, the main trunk of the left MCA was ligated just underneath the rhinal fissure with a 10-0 suture, and the bilateral common carotid arteries (CCA) were occluded for 60 min with 4-0 sutures. The sutures were then removed to restore blood flow, and the cervical incision was closed. Sham-operated rats (n=19) did not receive occlusion of either the MCA or the CCAs. Serial coronal sections were processed by NeuN immunostaining and all affected regions were identified. Six reproducible damaged regions were identified by 2 independent observers include parietal association cortex (PtA), agranular insular cortex dorsal part (AID), agranular insular cortex ventral part (AIV), dysgranular insular cortex (DI), granular insular cortex (GI) and primary somatosensory cortex (S1).

#### The intracerebral hemorrhage model (ICH)

Collagenase was injected in AP: -1.3 mm, L: 3 mm, V: 6 mm in 23 SD rats (right hemisphere). Lesioned regions were identified in the same manner as the dMCAO model. After summarizing subregions, a core subset of 22 significant lesioned regions was used for further analysis: endopiriform system (EnN), basal nucleus Meynert (B), angular thalamic nucleus (Ang), anteroventral thalamic nucleus (AV), centrolateral thalamic nucleus (CL), paracentral thalamic nucleus (PC), mediodorsal thalamic nucleus lateral part (MDL), ventroanterior thalamic nucleus (VA), ventrobasal complex (VB), ventromedial thalamic nucleus (VM), laterodorsal thalamic nucleus (LD), reticular thalamic nucleus (Rt), bed nucleus of the stria terminalis medial division (BSTM), central amygdaloid nucleus (Ce), interstitial nucleus of the posterior limb of the anterior commissure (IPAC), amygdalostriatal transition area (AStr), basolateral amygdaloid nucleus (BL), central amygdaloid nucleus medial division (CeM), lateral globus pallidus (LGP), medial globus pallidus (MGP), caudate putamen (CPu), dorsal endopiriform nucleus (DEn).

#### The intraluminal suture occlusion of the middle cerebral artery (sMCAO)

Stroke was induced by the intraluminal suture occlusion of the MCA (sMCAO or Suture) for 60 mins in 11 SD rats. Summarizing of subregions resulted in 57 significant lesioned regions: ventral endopiriform nucleus (VEn), nucleus of the horizontal limb of the diagonal band (HDB), nucleus of the vertical limb of the diagonal band (VDB), basal nucleus Meynert (B), substantia innominata (SI), lateral preoptic area (LPO), magnocellular preoptic nucleus (MCPO), ventrolateral preoptic nucleus (VLPO), medial preoptic area and medial preoptic nucleus lateral and medial (MPA/MPOL/MPOM), ventromedial hypothalamic nucleus central part (VMHC), anterior hypothalamic area posterior part (AHAP), lateral hypothalamic area (LH), anterior hypothalamic area (AHA), anteromedial thalamic nucleus (AM), ventrolateral thalamic nucleus (VL), ventral posterolateral thalamic nucleus (VPL), ventromedial thalamic nucleus (VM), ventral reuniens thalamic nucleus (VRe), reticular thalamic nucleus (Rt), zona incerta (ZI), tuber cinereum area (TC), anterior basomedial nucleus (BMA), posterior basomedial nucleus (BMP), dorsolateral part of the lateral nucleus (LaDL), ventrolateral part of the lateral nucleus (LaVL), bed nucleus of the stria terminalis (BST), central amygdaloid nucleus (Ce), interstitial nucleus of the posterior limb of the anterior commissure (IPAC), anterior amygdaloid area (AA), cortical amygdaloid nucleus (CAn), nucleus of the lateral olfactory tract (LOT), medial amygdaloid nucleus anterodorsal part (MeAD), amygdalohippocampal area anterolateral part (AHiAL), amygdalostriatal transition area (AStr), intercalated masses (IM), intercalated nuclei of the amygdala (I), basolateral amygdaloid nucleus (BL), central amygdaloid nucleus medial division (CeM), lateral accumbens shell (AcbShl), accumbens nucleus core (AcbC), caudate putamen (CPu), piriform cortex (Pir), cingulate cortex area 1 (Cg1), cingulate cortex area 2 (Cg2), temporal association cortex 1 (TeA), secondary auditory cortex (Au2), agranular insular cortex (AI), dysgranular insular cortex (DI), granular insular cortex (GI), perirhinal cortex (A35), ectorhinal cortex (A36), primary somatosensory cortex (S1), secondary somatosensory cortex (S2), secondary visual cortex lateral area (V2L), lateral orbital cortex (LO), olfactory tubercle (TuO), claustrum (Cl).

### Neurobehavioral assessment

Motor function was evaluated from 5 weeks after stroke or sham surgery in the order listed below: while spatial memory function was determined 8 weeks after stroke or sham surgery.

#### Catwalk-assisted gait test

Rats were subjected to 3 consecutive runs of gait assessment using the CatWalk automated gait analysis system (Noldus Information Technology) as described previously [132, 181, 196, 198]. The images from each trial were converted into digital signals and processed following the identification and labeling of each footprint. Intensity, maximum area and stride length were analyzed.

#### Rotor-Rod test

The task requires the rat to balance on a rotating rod. After a 1-min adaptation period on the rod at rest, rats were acclimated to the rotating rod by 5 rpm every 15 sec, and the total time the rat remained on the rod (also known as fall latency) was recorded. Before and after assessments of roto-rod performance, the equipment was cleaned with 1 mM acetic acid to remove residual odors [112, 144].

#### Horizontal ladder test

Rats were videotaped while traversing a 15o ladder with variable spacing between bars. The percentages of footfalls (slipping through the bars) with the affected or unaffected contralateral limbs were recorded and averaged from 3 trials [182, 198].

#### Barnes maze test

A black acrylic escape tunnel was placed under one of the holes on a circular platform (120 cm in diameter) with 40 holes (6 cm in diameter per hole) along the platform perimeter. Rats from each treatment group were randomly assigned to locate the escape tunnel from one of the four pre-determined locations to rule out spatial preference. In order for the rats to find and get into the box, bright light (300-350 LUC) and blowing fans (strong enough to move the fur) were provided. Once the rats located the box they were allowed to remain on it for 15 seconds. The rats were trained to locate the box from different counterbalanced starting positions in 2 successive daily sessions for for 5 days (3 trials per session, 3 minutes per trial), with a 1-hour intersession interval. The performance was analyzed by Noldus EthoVision video tracking system (Noldus). Following 30 trials of acquisition test, a 1-min probe trial was conducted 24 hours after the last trial on day 6 to determine memory retention with the hidden box removed [81, 144, 181, 198].

### Assessment of neural activation during spatial exploration

Functional recruitment of hippocampal circuitry was determined by mapping/quantifying the Fos expression immediately following spatial exploration in a seperate cohort to avoid the influence of spatial exploration on behavioral tests. Rats were either maintained in standard housing conditions (“home”) (*n* = 22) or underwent a spatial exploration task (“expl”) (*n* = 18), consisting of 15-minute exploration of an open field on a circular table (120 cm in diameter) with a large ball placed on the table each on days 4 and 5 after MCAO or ICH [156, 191]. The spatial configuration of the environment was altered between sessions post MCAO simply by changing the position of the large ball from the center (days 4) to the periphery (days 5) of the table with everything else remaining the same. Ninety minutes after the exploration task (or at the same time of the day for “home cage” controls), rats were perfused with 4% paraformaldehyde and 40 µm coronal sections were stained for the expression of Fos (rabbit-anti-Fos, Oncogene Science, 1:5000) [111, 156]. Fos-positive cells were counted manually under a light microscope at 40x magnification using the NIH *ImageJ* cell counting plugin tool. Three sequential serial sections (480 µm apart) were analyzed per animal and the counts were averaged and expressed as number of cells/mm^2^.

### Lesion size assessment

At the end of Barnes maze test, rat brains were sectioned and processed for NeuN immunostaining. Infarct volume was measured by subtracting the volume of intact tissue in the ipsilateral hemisphere from that in the contralateral hemisphere on NeuN-stained serial coronal sections (480 µm apart) by *ImageJ* (NIH) as described previously [5, 181].

### *neuroVIISAS* -based rat brain connectomics

The neuronal connections of the rat connectome have been collated in a metastudy [162]. For this, peer-reviewed publications which described results of tract-tracing experiments in juvenile to adult healthy rat strains were manually evaluated. All sources and targets of neuronal connections of each experimental observation and their semiquantitative or quantitative connection weights (e.g. light, moderate, strong) were gathered. All connectional data and features of neuronal connections were imported into the rat nervous system project of the generic framework *neuroVIISAS* [107, 157]. The rat connectome project covers all described connections documented in more than 7800 tract-tracing publications since 1972. All network analyses used in this contribution have been conducted consistently in the *neuroVIISAS* framework as shown in further studies [158, 160]. Verifications, reliability and reproducibility studies of the rat connectome managed in *neuroVIISAS* showed that the collation of neuronal connections based on tract tracing studies have a low interrater variability [162] and can be reproduced with other methods [174]. Further details are described in [156].

#### Differential connectomics by comparing the control and lesioned connectome

To investigate how connectivity is affected in each of the stroke models, we determined the control connectome and lesioned connectomes following mapping the damaged regions using stereotaxic rat brain atlas as described above [139]. The control connectome consists of damaged regions and their interconnected regions, while the dMCAO, sMCAO, and ICH lesioned connectomes are built from the control connectome minus the lesioned areas of each model, whereas the control connectome comprises of the damaged regions and their connecting regions. Differential connectomics and lesion connectomics were conducted to quantify the connectome changes specifically induced by each model [117].

#### Assignment of motor and learning behavioral networks

By assigning functional attributes to regions of the connectome it becomes possible to determine functional network changes following stroke experiments. A total of 13 regions are identified for the control of motor functions including primary cortical regions (AGl=M1, AGm=M2), basal ganglia regions (LGP, MGP, CPu, SNR, SNC, STH, VL) and cerebellar regions (CERC, DCeN, Pn) [74, 95, 138]. Motor nuclei of the cranial nerves, visceral motor neurons and spinal cord motor neurons or regions for hypothalamic goal directed behavior are not considered here [200]. Nineteen regions involved in learning behavior are identified including CA1, CA2, CA3, DG, S, LEnt, CMAM, Cg1, Cg2, PrS, PaS, A35, POR, AVv, IAM, Po, RhN, SPFr and ADR [3, 49, 126, 145, 185].

#### Construction of the multidimensional circular relationship diagrams (MCRD)

Based on the following 5 primary parameters, each model’s own MCRD presents the afferent and efferent connections between each lesioned area and regions of the control connectome: (1) the direction of a connection (afferent, efferent or reciprocal), (2) the weight (color code), (3) the number of observations of a connection in tract-tracing studies (line thickness) and (4) the average rank of its local parameters (distance of a region which is connected with the center region of the MCRD), and (5) the number of connections with other lesioned regions is coded as the arc length of the border of the MCRD. If connections between lesioned and non-lesioned regions were reported more than 2 times in tract-tracing publications (coded by line thickness), they were deemed more reliable (RELi) connections. If a non-lesioned region has connections with more than one lesioned region, we perceived it as multiple lesional effect (MULi), which is represented by the distance of regions to the center of the MCRD. The importance of a region in the network (NETi) can be estimated by the average rank of local network parameters. To effectively identify the most important non-lesioned regions sustained the greatest impact of stroke lesion, with smallest average ranks of local network parameters, a triple filtering with the parameters RELi=2, MULi=4, NETi=80% was performed to select out the top 20% most important regions (high ranks or low rank numbers) in the network.

### Structural network analysis

Lesional effects have been quantified via methods of graph theory for the different global and local network parameters as well as particular matrix representations of the connectomes. The following 6 global network parameters were comparatively applied to the 3 lesion models and are discussed in a subsection in the Results section.

Global network parameters include the reciprocal edges, average path length, average cluster coefficient, small-worldness, transitivity and the knotty centrality or centeredness. The following is a very brief consideration of the general definitions and interpretations of these 6 global network parameters.

*Reciprocal edges* indicates the number of neuronal connections between two regions that can send signals from region 1 to region 2 as well as from region 2 to region 1. The reciprocity of a directed graph can be calculated by the ratio between the number of edges oriented in both directions and the total number of edges in the graph.

The *average path length* or *average shortest path length* is defined as the average number of steps along the shortest paths for all possible pairs of network nodes or in other words the average minimal number of edges of all possible node connections [6]. The average path length is calculated from all possible shortest paths such as region 1 to region 2 and region 2 to region 1 by summing all the path values and dividing by the number of paths.

The *clustering coefficient* of a network is a measure of the degree to which nodes in a connectome tend to cluster together. The *local clustering coefficient* is computed for each node in the connectome. The local clustering coefficient of a region describes the likelihood that the neighbors of the region are also connected. To calculate local clustering coefficient we use the number of triangles a node is a part of it, and the degree of the node. At the end the local cluster coefficients are needed to compute the average clustering coefficient for the whole graph by normalizing the sum over all the local clustering coefficients.

The global network measure *small worldness* quantifies the expression of the small world property of a network. The small worldness is determined in the first step by means of the transitivity of the connectome and the average length of the shortest path. Then, the average of the same indices is calculated for a given number of random networks. The small-worldness index is then calculated as the transitivity (normalized by the random transitivity) over the average shortest path length (normalized by the random average shortest path length).

The *transitivity* is the total probability that neighboring nodes are connected in a network, indicating the presence of closely connected clusters. It is calculated by the ratio between the observed number of closed triplets and the maximum possible number of closed triplets in the graph.

*Knotty-centredness* or *knotty-centrality* is a measure of topological centrality and quantifies the extent to which a given subset of regions of a network constitutes a densely intra-connected topologically central connective core [169]. The knotty-centredness can be determined by exhaustively searching all subsets of the connectome whose members fall into the top knots for betweenness centrality and then using the gradient ascent.

To calculate the mean ranks of local network parameters, they were transformed to have comparable numerical ranges. Some exemplary parameters from the total of 49 different local network parameters are: *DG_All_* (Degree all), *Katz* (Katz status index), *CluC_All_* (Cluster coefficient all), *Lev* (Levarage), *Loc* (Locality index), *LE* (Local efficiency), *BC* (Betweeness centrality), *EC* (Eigenvector centrality), *FC* (Flow coefficient), *Shapley* (Shapley rating).

A detailed definition of the parameters have been published previously [148, 158].

#### Connectivity matching index (CMI)

The CMI is a measure for the amount of overlap in connections between a pair of regions and was used with regard to lesioned regions and their connections with functionally defined regions. It can be computed by combining afferents or inputs to regions (*CMI_In_*) with efferents or outputs from regions (*CMI_Out_*) as *CMI_All_*. If two connected regions are very similar with regard to their connections then the values of *CMI_All_* become large. A *CMI_All_* index of 1 refers to the situation that afferents and efferents of region “*i* ” are exactly the same as those of region “*j* ”. The following formula shows the normalized (denominator) calculation of *CMI_All_* of a pair of nodes or regions i and j (A: adjacency matrix, k: number of similar connections):

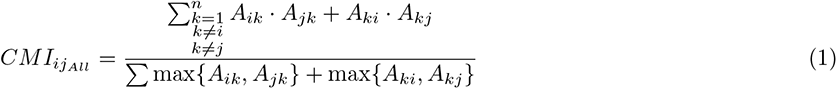

#### Generalized topology matrix (GTOM)

GTOM has been applied to the dMCAO, ICH and sMCAO adjacency matrix [206] for a formal definition see also [158]. It is a measure of pairwise interconnectivity that is proportional to the number of neighbors (two distinct directly connected brain regions) that a pair of nodes respectively a pair of distinct brain regions have in common. The measure is a count of the number of m-step neighbors that a pair of brain regions share and is normalized to a value between 0 and 1. Because we found a stronger connectional relationship of the anterior basal nucleus using GTOM just for the ICH model, only this finding is presented and no further properties of the GTOM matrices for the other two models.

### Functional network analysis

Emerging evidence suggests that changes in connectivity can affect propagation of oscillations in connectomes as reflected by changes in synchronization, coherence, and frequency of oscillations in neural networks. For example, the strength and directionality of connections between neurons can have a significant impact on the emergence and frequency of oscillations [165]. Three established models can be used to simulate the neural oscillatory behavior in a defined network.

#### The FitzHugh-Nagumo (FHN) model

The dynamic effect of stroke lesion in each model on learning/memory function was predicted by modeling the coactivation pattern and interspike variability between neurons in each lesioned region and functional region using the FitzHugh-Nagumo (FHN) model, which pertains an excitable system like a neuron and it acts as a relaxation oscillator if an external stimulus passes a certain threshold Statistical mechanics of complex networks [53, 128]. The FHN model was applied in the connectome by coupling the FHN neurons through the weighted connectome connections [211] in order to investigate the propagation and mutual interference of oscillations in the connectome rather than to reconstruct and simulate microcircuits of neuron populations. Coactivation of two regions arises from common neighbors or similarly connected regions. However, a connection between two regions reduces the likelihood of coactivation because a sequential excitation of the two regions, leading in principle to a negative effect of structural and functional connectivity. Nevertheless, the global topological or connectome-based oscillation transfer can annihilate this effect. The coupled FHN system has been stimulated and after 500 ms or steps the coactivation of pairs of regions or FHN neurons have been computed and visualized in a coactivation matrix, in which a large coactivation value reflects similar FHN activity between two regions. The FHN model has been adapted to be used in weighted adjacency matrices (D) in our study. Sufficient accuracy was obtained by solving the equations with a forward Euler approach. Parameter exploration suggests a slight variation of parameters in comparison with [211] as follows: *α* = 0*, β* = *−*0.064*, γ* = 1.0*, ϕ* = 1.0*, E* = 2.0*, k* = 1.0*, σ* = 0.25*, τ_X_* = 1.0*, τ_Y_* = 1.0*, x*_0_ *<* 2.0*, x*_0_ *>* 2.0*, y*_0_ *<* 2.0*, y*_0_ *>* 2.0. *x* is the membrane potential and y the recovery variable. *_X_* and *_Y_* are time scale factors for *x* and *y*. *k* is a global scaling parameter for the coupling strength and σ is a scaling factor for uncorrelated Gaussian noise ν with 0 mean and unit variance. ε is a scaling factor for the recovery function. *D* is the transposed and weighted adjacency matrix of the connectome and *f* (*x*) is the logistic coupling function.

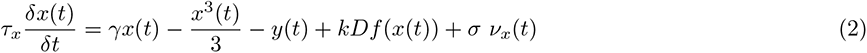

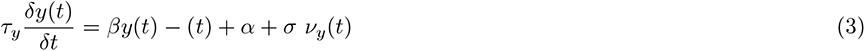

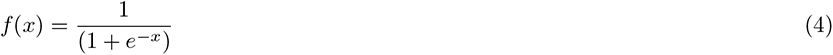

One FHN-neuron (Figure 1) was modeled for each node of the connectome and connectome connections were used as the coupling matrix. After a simulation time of 500 ms with the step size of 1 ms the coactivation matrix has been calculated. It was used in the same way as the *CMI_All_* matrix for pairwise ranking. For the coactivation matrix *Co* a spike detection *Ci*(*t*) on the basis of average excitation and standard deviation of excitations of a region need to be performed:

**Figure 1.**
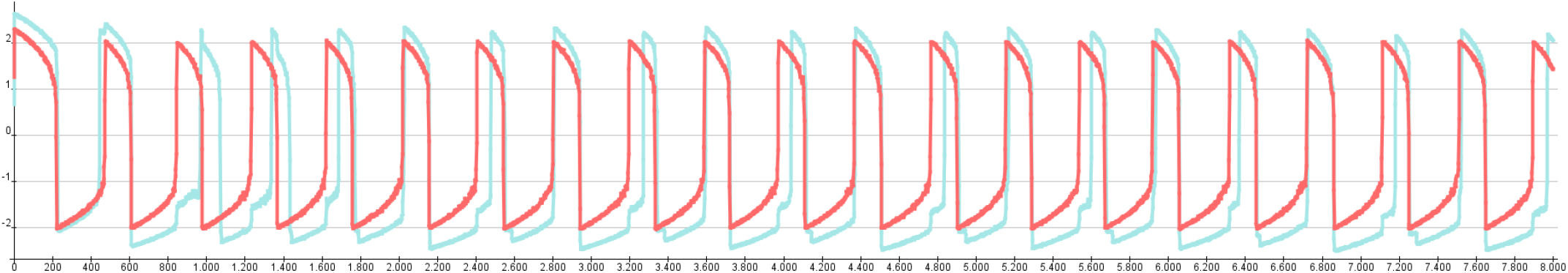
The membrane potentials of a FHN model. The FHN dynamics of two coupled FHN neurons over 8000 ms. The same parameters were used by [211], however, ε was set to 1.

In most cases a relative coactivation to excitatory states *Co_ij_*has been calculated instead of a coactivation relative to time steps (1*/t_max_*Σ*C_i_*_(*t*)_*C_j_*_(*t*)_):

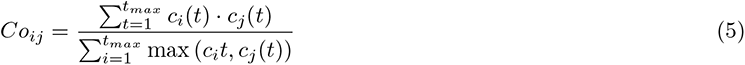

### Functional Network Analysis by modeling hippocampal network dynamics

To reveal dynamic changes resulting from connectivity loss that led to hippocampal functional impairment [19,33,40,60,168], we tested if differences of signal patterns in the hippocampal regions CA1 and DG [8, 177] appear in dynamic modeling after removing stroked lesioned regions. Since changes in information transmission from inter-cortical networks have been documented after stroke [71, 91], we investigated how signal propagate in connectome after stroke taking into considerations of connectome properties such as directionality [108], weighting, axon collaterals, bilaterality, completeness and transsynaptic pathways [193, 195].

#### Wilson and Cowan model

The Wilson and Cowan model is a neural mass model that provides a framework for studying large-scale neural networks in the brain. It is commonly used to model the dynamics of populations of excitatory and inhibitory neurons in the cortex, and is especially useful for studying the generation of oscillations and other complex patterns of neural activity. The model is based on the idea that the activity of a population of neurons can be described by a single variable, which represents the average firing rate of the population. It has been successfully applied to study signal propagation in neural networks. The application in mouse cortex models allowed for the first time a mechanistic interpretation of information transmission in a realistic neuronal network [175]. Therefore, we also adopted this approach to analyze signal-stimulus response changes in a connectome perturbed by stroke using another model category.

The Wilson and Cowan model describes the dynamics of two interacting populations of neurons, an excitatory population (E) and an inhibitory population (I). The specific equations depend on the assumptions made in the model, such as the properties of the neurons, the sigmoidal activation functions, the strength and directionality of the connections, and the external inputs. The model assumes that the activity of each population can be described by a single variable, which represents the average firing rate of the population. Let x(t) and y(t) be the average activity of the excitatory and inhibitory populations, respectively, at time t. The dynamics of the model are governed by the following set of coupled differential equations:

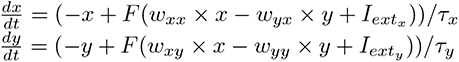

where:

*• F* is a sigmoidal activation function that captures the non-linear relationship between neural activity and synaptic input, typically given by *F* (*z*) = 1*/*(1 + *e^−z^*)
*• w_xx_*, *w_yx_*, *w_xy_*, and *w_yy_* are the connection strengths between the populations, where *w_xx_* and *w_yy_* are the self-connections of the populations
*• I_extx_* and *I_exty_* are external inputs to the populations
*• tau_x_* and *tau_y_* are time constants that govern the decay of the activity in the populations.

These equations describe the evolution of the activity of the excitatory and inhibitory populations over time, taking into account the interactions between the populations through their connections and the external inputs to each population. The Wilson and Cowan model is useful for studying the dynamics of large-scale neural networks in the brain, and has been used to model a wide range of phenomena, including the generation of oscillations, the emergence of spatial patterns of activity, and the effects of neuromodulation on neural activity. The model can also be extended to include additional populations of neurons, as well as more complex network architectures, making it a versatile tool for studying the neural mechanisms underlying cognition and behavior.

#### Mimura-Murray reaction-diffusion model

Flow-based network analysis using diffusive processes (network diffusion) [1] has been successfully used to study the relationship between structure and function in the nervous system. Based on the Lotka and Volterra model, the diffusion efficient and flexible reaction-model was developed by Mimura and Murray. In the context of neural networks, the model has been used to study the formation of spatial patterns of activity in directed networks, where the activity of one population of neurons affects the activity of another population through excitatory or inhibitory connections [163]. The model can also be extended to include additional populations of neurons, as well as more complex network architectures, making it a powerful tool for studying the dynamics of large-scale neural networks.

In a more formal definition the reaction-diffusion model of Mimura-Murray is a system of partial differential equations that describe the spatiotemporal dynamics of a system of interacting objects. Here, the two populations are considered on a domain with two spatial dimensions in such a way that the PDEs have the form [123]:

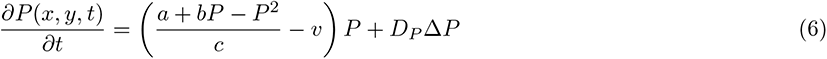

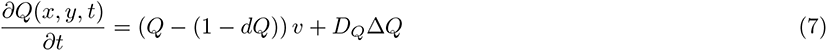

Parameters of the local reaction of the Mimura-Murray model were used as suggested by [9] *a*: 13, *b*: 16, *c*: 9, *d*: 0.4, *D_u_*: 0.1, *D_v_*: 0.01.

#### Neural population simulation engine Neural Simulation Tool (NEST)

To investigate the particularly large lesion effects of the sMCAO experiment in a dynamic model, we used the interface of *neuroVIISAS* [157] and NEST [79, 129]. In *neuroVIISAS*, each parameter of the neuron models available in NEST can be specified precisely, the populations and couplings (weighted and directed neuronal connectivity) can be defined exactly, and the simulation in NEST can be executed efficiently on parallel computers. The results of the simulations are read into *neuroVIISAS* and can be dynamically visualized in the available digital atlases in *neuroVIISAS* as well as coupled with structural network data and statistically analyzed. In the following, we will briefly describe the well-known leaky integrate-and-fire neuron model, which is particularly suitable for use in neuron populations.

The leaky integrate-and-fire (LIF) model is a simplified mathematical model commonly used in neuroscience to study the behavior of neurons for building artificial neural networks. The LIF model describes a neuron as a simple electrical circuit, where the neuron’s membrane potential is represented by a voltage variable, *V* . The model assumes that the neuron receives input currents, *I*, from other neurons or from the environment. These input currents cause the neuron’s membrane potential to change over time, according to the equation:

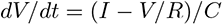

where *R* is the neuron’s membrane resistance and *C* is its capacitance. This equation describes the rate of change of the membrane potential, *dV/dt*, which depends on the difference between the input current *I* and the membrane potential *V*, divided by the product of the resistance *R* and the capacitance *C*. The leaky part of the LIF model refers to the fact that the neuron’s membrane potential leaks over time due to the presence of ion channels that allow ions to pass through the membrane. This leakage is modeled by adding a term proportional to the difference between the resting potential, *V_rest_*, and the current membrane potential, *V*, to the equation:

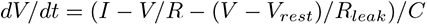

where *R_leak_* is the membrane leakage resistance. This equation describes how the membrane potential changes over time, including the effect of the input currents and the leakage. The integrate-and-fire part of the LIF model refers to the fact that the neuron ”fires” when its membrane potential reaches a certain threshold value, *V_thresh_*. When the membrane potential exceeds this threshold, the neuron generates an action potential or spike, which is transmitted to other neurons. After firing, the membrane potential is reset to a lower value, *V_reset_*, and the neuron enters a refractory period during which it cannot fire again. This behavior is modeled by adding a threshold condition to the equation:

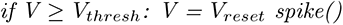

where *spike*() is the function that generates an action potential or spike. In summary, the leaky integrate-and-fire model describes the behavior of a neuron as a simple electrical circuit, taking into account the effect of input currents, leakage, and firing threshold. For the simulation of the sMCAO connectome we parameterized the LIF model of NEST by 80% excitatory neurons (total of 229116 LIF neurons). The synaptic delay was set to 1 ms, the locus coeruleus was stimulated by 10kHz and simulation time was 300 ms. Spike distributions were computed and interspike intervals were calculated. An interspike interval (ISI) refers to the time interval between two consecutive action potentials or spikes generated by the same neuron. When a LIF neuron receives input currents that cause its membrane potential to reach the firing threshold, it generates an action potential or spike, which is transmitted to other neurons in the population. After firing, the neuron’s membrane potential is reset to a lower value, and the neuron enters a refractory period during which it cannot fire again. The duration of this refractory period, combined with the input currents received by the neuron, determines the interspike interval between successive spikes. In the LIF model, the refractory period is typically modeled as a fixed duration during which the neuron cannot fire again, regardless of the strength of the input currents. The interspike interval is an important measure of the activity of neurons in a population, as it reflects the frequency and regularity of their firing. By analyzing the distribution of interspike intervals across the population, we can gain insights into the dynamics of the neural network and the functional properties of the neurons. For this purpose, we calculated and compared the variability of the spike intervals using the mean error coefficient.

### Statistical analyses

To determine differences among stroke models in global and local network parameters, ANOVA, student t-test or U-test will be used. To determine whether there was a significant difference between control and lesioned connectomes of the 3 stroke models, we applied the concept of permutation of random networks (null models) [20, 55, 118, 179, 194]. Multivariate analyses were used to determine groups of classes of regions with modularity testing, hierarchical cluster analysis (spectral clustering [18, 109, 154, 201, 212], Markov chain clustering [43], Girvan-Newman [106, 114, 133, 134], Louvain modularity [22], principal component analysis [PCA] [48, 140], multidimensional scaling [MDS] [205], self-organizing maps [SOM] [153]). We further defined and compared structural groups of regions among experimental models by analyzing connections weights, connectivity matching and local network parameters.

## Results

### The level of functional impairment depends on both: lesion size and location

The sMCAO model produced damage to the sensorimotor cortex, the lateral portion of the striatum, and the fiber tracks between the cortex and the basal ganglia. The dMCAO model on the other hand, caused damage only in the sensorimotor cortex. Finally, despite yielding the smallest infarct volume, the ICH model induced damage to the internal capsule, the striatum, and the anterior portion of the thalamus and its neighboring fiber tracts (Figure 2A). Regarding lesion size, sMCAO led to the most extensive damage, approximately twice the size as that in the dMCAO model, while the ICH model produced the smallest injury, 15% of the sMCAO infarct volume (Figure 2B).

**Figure 2.**
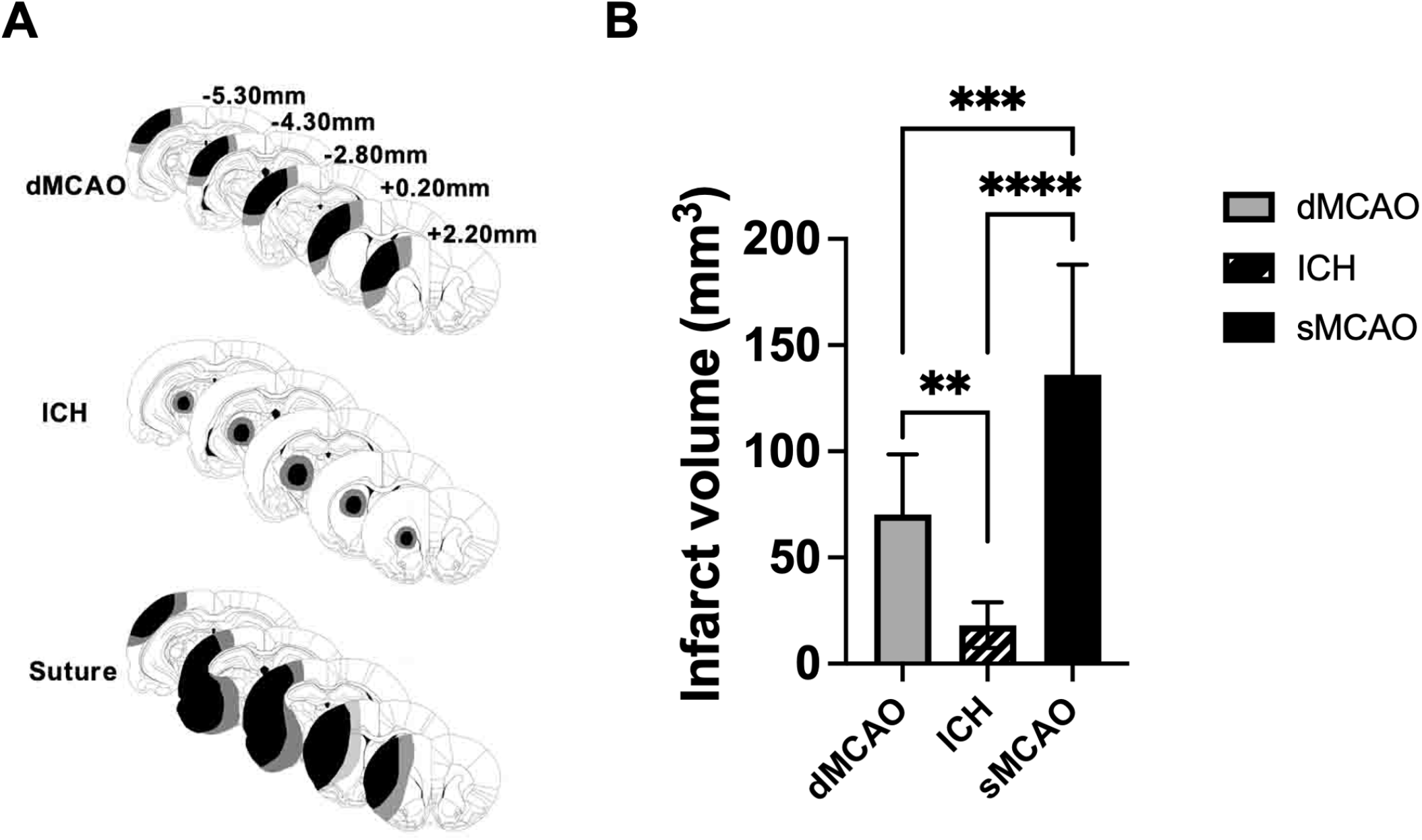
Comparison of the damage produced by the three types of stroke. (A) Reconstructions of coronal sections showing the extent of infarct in the rats of three stroke models. Smallest and largest damaged areas appear in black and gray, respectively. Numbers indicate the section distance in millimeters from Bregma. (B) The mean infarct volume of three stroke models. **P*<*0.01, ***P*<*0.005, ****P*<*0.001. N=9-14/group.

Motor function was examined using the accelerating rotor-rod, the ladder test, and the catwalk test. Although all three models showed impaired motor function in comparison to the sham group, rats subjected to the sMCAO model consistently exhibited the worst motor function outcomes in all the tests (Figure 3). Surprisingly, despite sustaining small infarct volumes, rats subjected to the ICH model also exhibited very poor motor function outcomes indistinguishable from the sMCAO rats such as the rotor rod test (Figure 3A), and some of the intensity parameter of the catwalk gait test (Figure 3C). Rats that have undergone dMCAO appeared to suffer the least extent of motor impairment among 3 models, or remaining a comparable level of performance in some tests relative to the ICH rats. (Figure 3).

**Figure 3.**
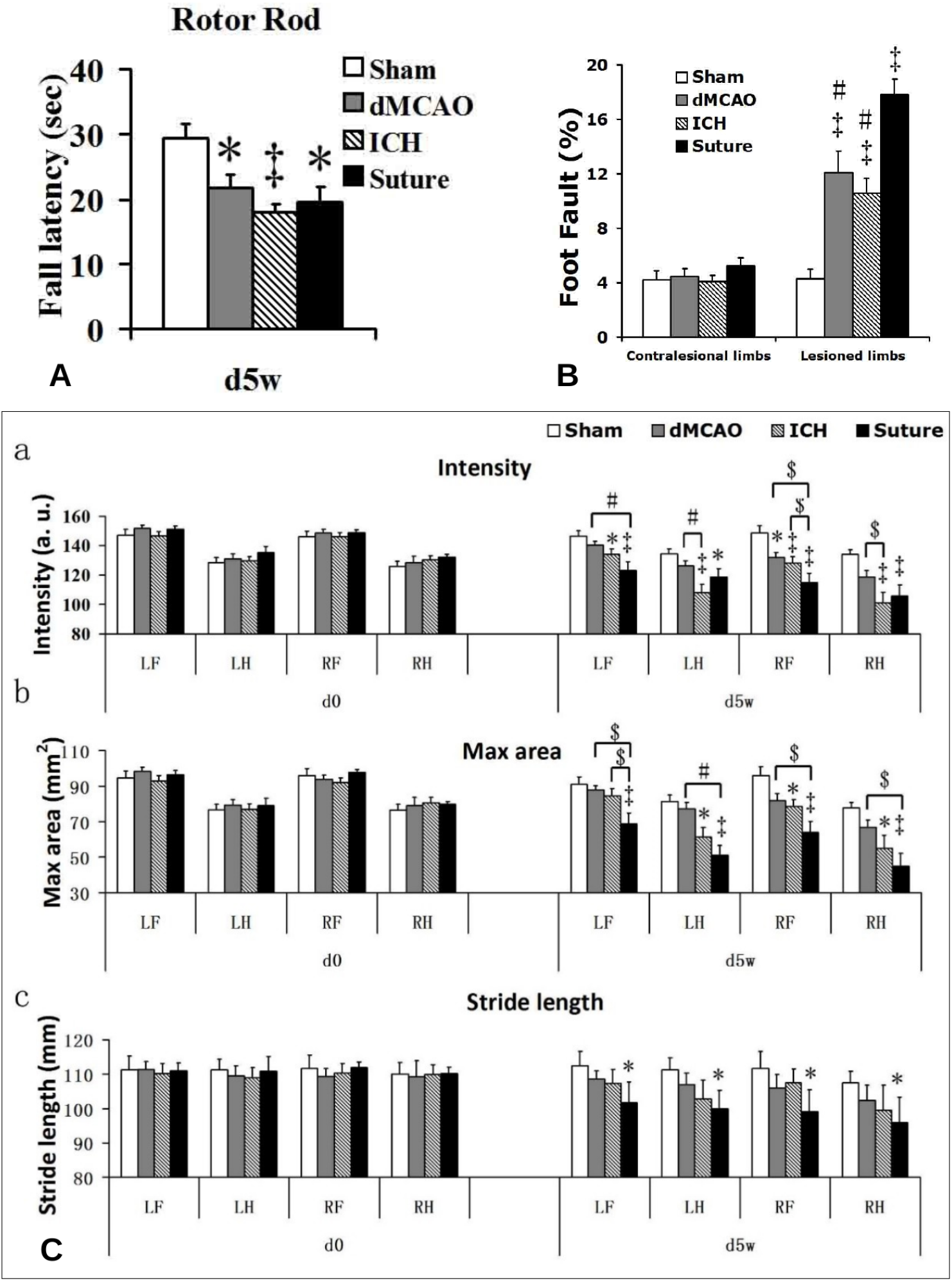
Motor function impairment in the three stroke models. (A) Performance on the accelerating Rotor Rod in all three stroke groups of rats had the significant motor function impairment compared to the sham rats (*p*<*0.05; *‡*p*<*0.01), but no difference among the three stroke groups was seen. (B) Ladder test. Performance on the ladder test in all the three stroke groups of rats had significant motor function impairment compared to the sham rats (*‡*p¡0.01), and the impairment of the sMCAO MCAO rats was significantly worse than those of the other two stroke groups (#p*<*0.01). dMCAO and ICH group vs sMCAO MCAO group). No difference between the dMCAO and ICH stroke groups. (C) Catwalk test. Effects of stroke on paw pressure (a), area of paw contacts (b) and stride length (c). Gait parameters were assessed at preoperative (d0) and 5 weeks (d5w) after left stroke surgeries. All three parameters had no difference among the four groups before stroke surgery, but five weeks after stroke surgery, the sMCAO MCAO rats presented very severe gait impairments compared to the sham rats, especially the impairments of paw intensity and maximal area. The paw intensity, max areas and stride length were reduced at all paws after sMCAO stroke. Different from the sMCAO MCAO, the dMCAO and ICH had no significant change in the temporal parameter of stride length, The paw intensity and max area of affected limbs of ICH rats had a similar significant decrease with sMCAO MCAO. The paw intensity and max area were reduced at the paws after dMCAO, but only had significant difference in the paw intensity of the affected forepaw

All three stroke models showed impairment in spatial memory, as determined by path length to escape box in the Barnes maze, compared to the sham control group. sMCAO rats showed the greatest deficits in spatial learning in the Barnes maze, compared to comparable performance between the dMCAO and ICH rats (Figure 4).

**Figure 4.**
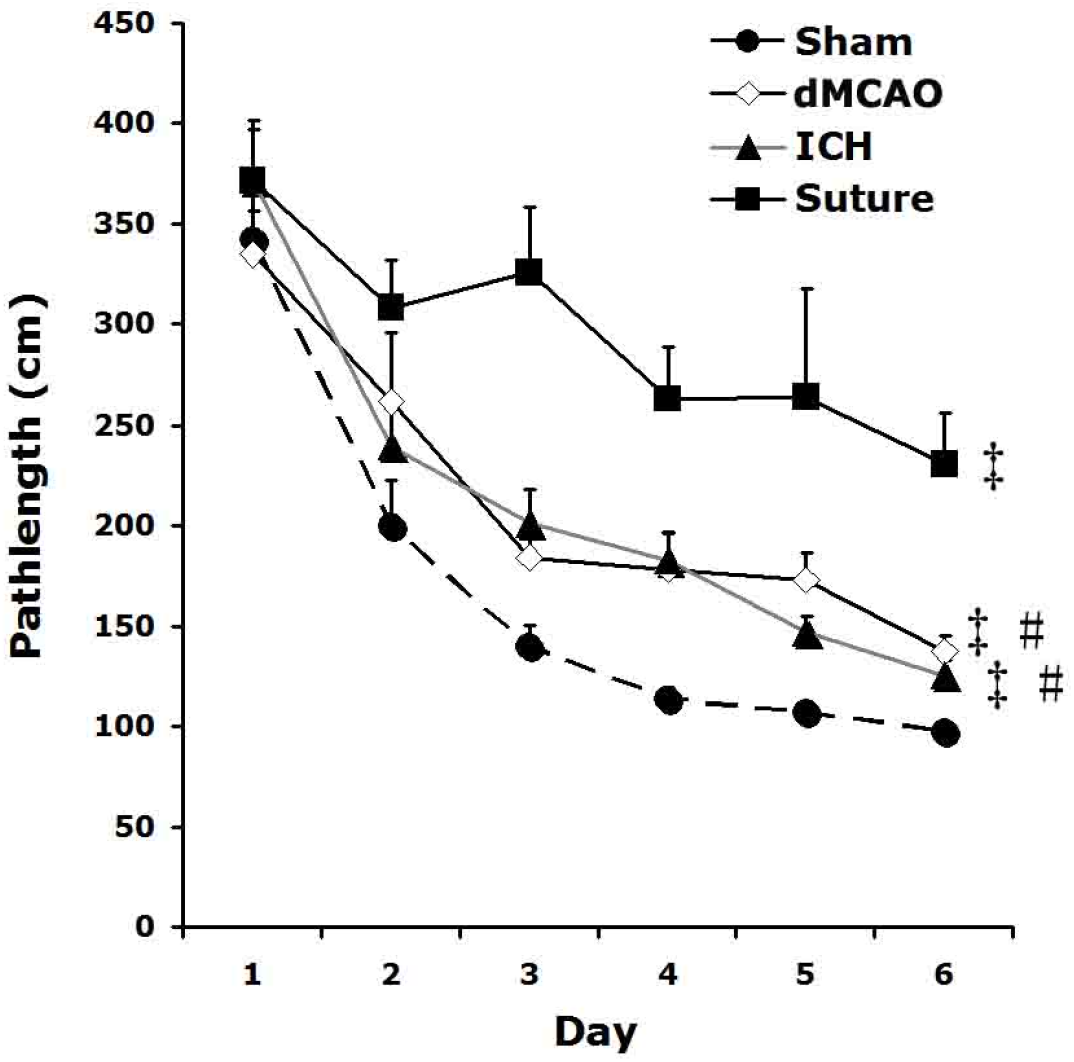
Spatial memory impairment of the three stroke models. All three stroked rats performed more poorly than sham controls during spatial learning in the Barnes maze, especially the sMCAO MCAO rats (*‡* p *<*0.01 vs sham). Their path length to locate the escape box in the Barnes maze was also significantly longer than those of dMCAO and ICH rats (# p *<*0.01 vs suture MCAO).

Hippocampal activity following spatial exploration was determined by Fos-immuno-reactivty in the CA1, CA3 and the DG regions. All rats that were allowed to spatially explore showed increased Fos expression; however, the amount of Fos-positive nuclei was markedly reduced in ischemic groups compared to the sham group. In all three hippocampal regions, the sMCAO group exhibited the least amount of Fos-positive nuclei, or the greatest extent of hippocampal hypoactivation, whereas the dMCAO and ICH groups exhibited comparable level of hypoactivation (Figure 5). Taken together, the results suggest that both large infarct size and subcortical injury location contribute to a greater degree of functional deficits.

**Figure 5.**
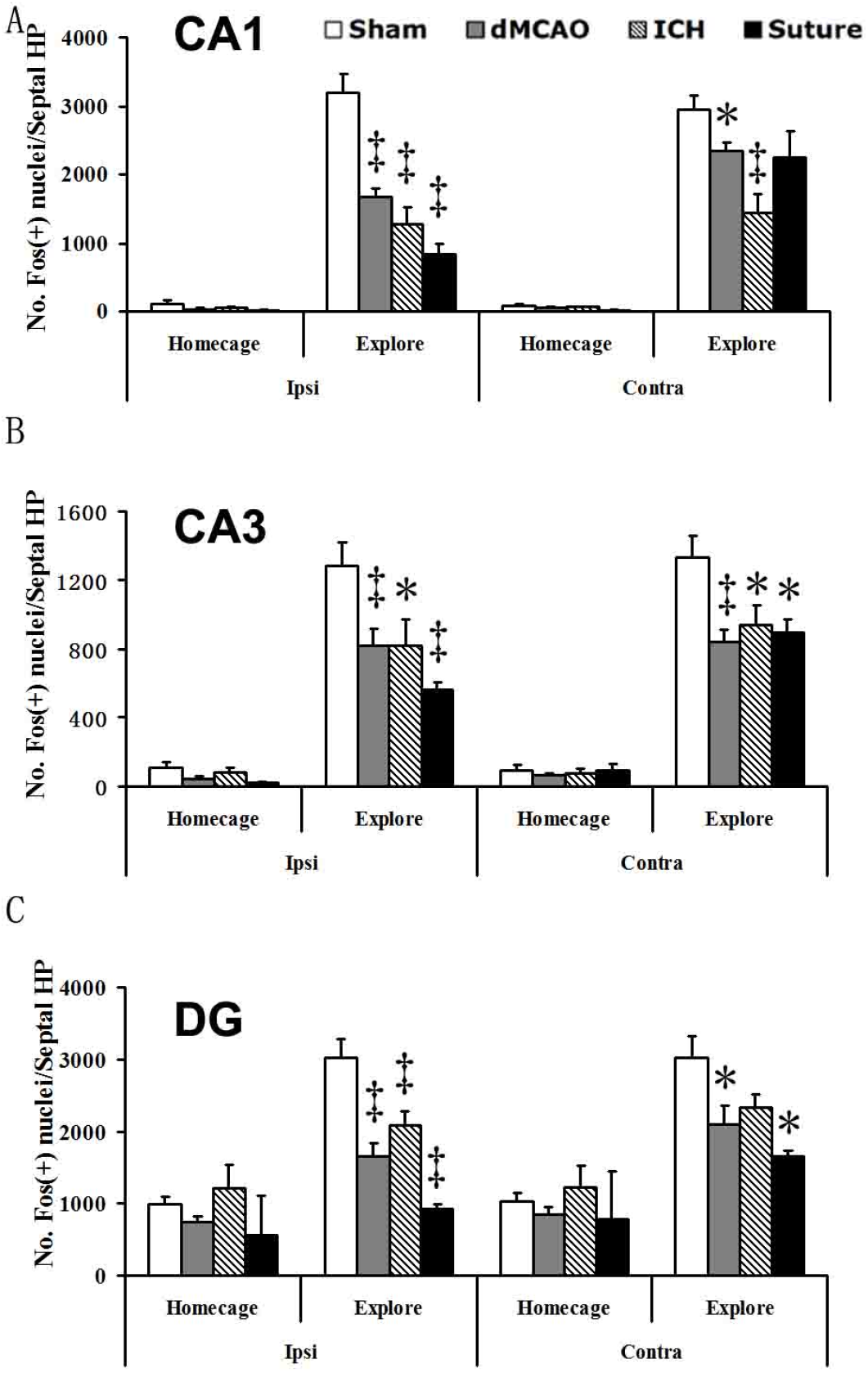
Stroke or hemorrhage reduces hippocampal activation following the exploration of a novel environment. (A) Counts of Fos-positive nuclei in the hippocampal CA1 region of shams and dMCAO rats remaining in their home cage or exploring a circular arena. Spatial exploration increased Fos expression (# p *<*0.01) but this effect was reduced in ischemic rats. Although affecting both hemispheres, it was predominant in the ipsilateral infarcted hemisphere (*P¡0.05; *‡*P¡0.01), (B) A similar pattern of effect was observed in the CA3 region of the hippocampus. Spatial exploration increased Fos expression compared to home cage controls (p *<*0.01). This effect was diminished in all stroked rats (*P*<*0.05; *‡*P*<*0.01). (C) In the DG of the hippocampus, spatial exploration enhanced Fos expression (p *<*0.01) and revealed hypoactivation in stroke.

#### Stroke induced a greater connectivity loss within the lesioned regions than the interhemispheric connections

Connectivity changes in each stroke model are reflected by adjacency matrices among the damaged regions (intrinsic) and with contralateral homotopic regions (extrinsic) 6. The ipsilateral lesioned right hemispheric regions in the dMCAO, ICH and sMCAO models are highly connected intrahemispherically compared to modest connections with the intact homotopic regions of the contralateral hemisphere. Hence, the connectivity of lesioned regions is greatly disrupted intrinsically. In the ICH model for example, BL has most output connections (histogram of rows of the adjacency matrix) among lesioned regions and the CPu most input connections (histogram of columns of the adjacency matrix).

By incorporating the lesioned regions into the existing *neuroVIISAS* connectome, the control connectome of each model provides a thorough connectivity mapping. Here we show an example of the control connectome of the dMCAO model in the form of a weighted and directed adjacency matrix with hierarchical organization of regions as well as column and row frequencies of connections (Figure 6). A concentration of connections is visible around the main diagonal in the lower right of the adjacency matrix, indicating a relatively strong interconnectedness of cortical regions.

**Figure 6.**
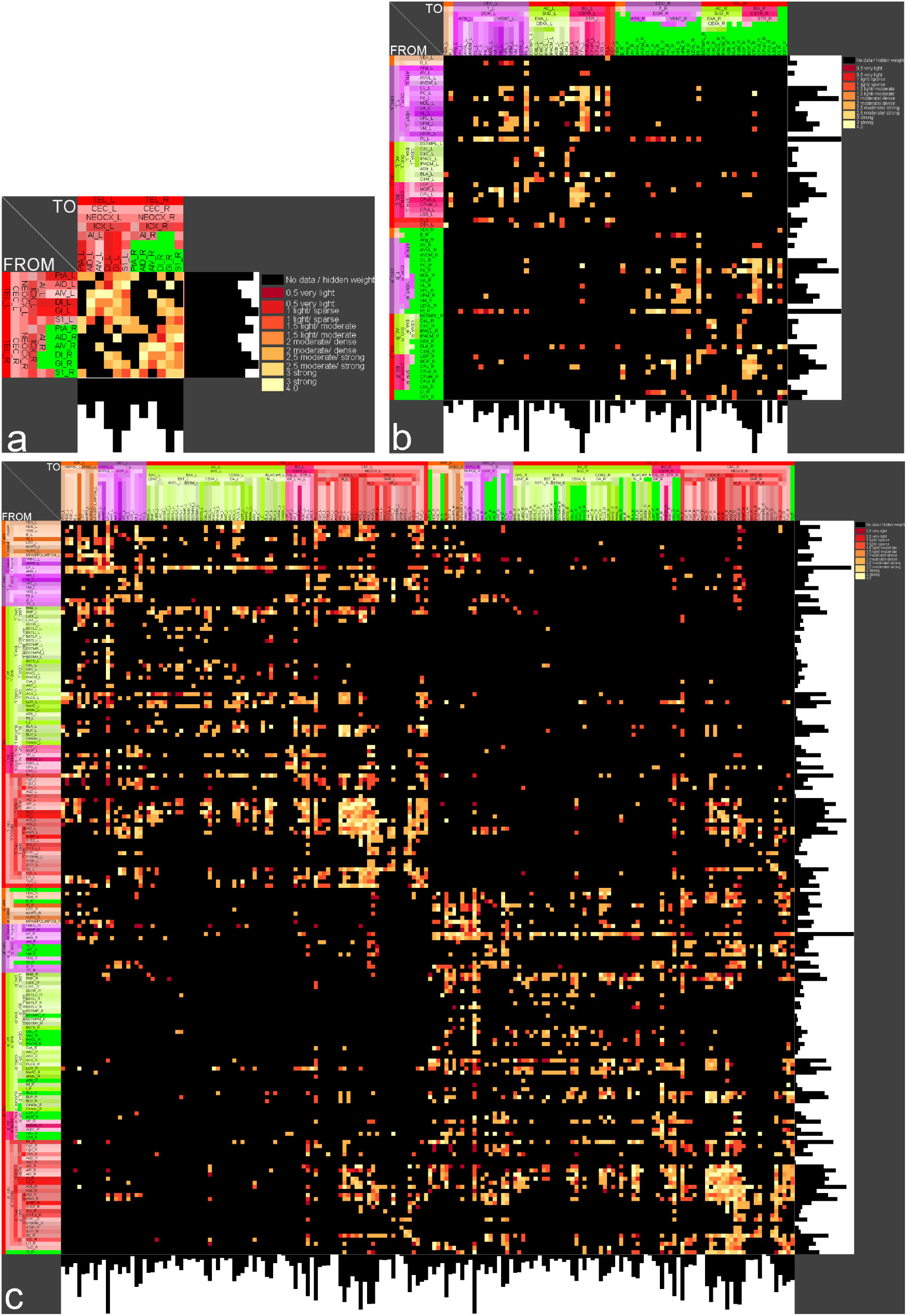
Weighted and directed bilateral adjacency matrices with right hemispheric lesioned regions. a) dMCAO model (6 lesioned regions: green), b) ICH model (22 right hemispheric lesioned regions: green), c) sMCAO model (57) right hemispheric lesioned regions (blue). Histograms of rows and columns represent the number of output and input connections, respectively.

#### Stroke altered the global network structure

Three lesioned connectomes were compared to the control connectome in the first step of our differential connectome study. All three lesioned connectomes displayed reduced total numbers of regions, links and reciprocal connections compared to the control connectome as summarized in Table 1, together with other changes of global network parameters. Not surprisingly, the sMCAO model with the largest lesion size resulted in the most numbers of lesioned regions (57), which gave rise to the greatest change in global network structure including the biggest loss of reciprocal connections by 59.81% and largest increase of small-worldness by 34.7%, significantly altering the network architecture. Surprisingly, albeit with a smaller lesion size, the ICH model had a greater change in global network structure compared to that of the dMCAO model.

**Table 1.** Comparison of global network structure change after stroke.

| Parameter | dMCAO | ICH | sMCAO |
| --- | --- | --- | --- |
| Regions in control connectome | 540 | 540 | 540 |
| Links in control connectome | 13183 | 13183 | 13183 |
| Lesioned regions | 6 | 22 | 57 |
| Regions in lesioned connectome | 534 | 518 | 483 |
| Links in lesioned connectome | 11693 | 10462 | 6883 |
| Reduction in reciprocal connections [%] | 8.52 | 20.4 | 59.81 |
| Increase in small worldness [%] | 6.41 | 12.24 | 34.7 |
| Reduction in cluster coefficient [%] | 0.748 | 0.705 | 19.36 |
| Increase of average pathlength [%] | 0.42 | 2.27 | 12.38 |

To better visualize the global connectome differences, some well-chosen global network parameters were comparatively visualized for the real tract-tracing connectome and randomized connectomes. The detailed results of the empirical control (red dots) and lesioned connectomes (blue dots) combined with 100 randomized Erdős-Rényi (ER) type and 100 null model of the rewiring or degree preserving network (RW) simulation of each of six global network parameters, such as reciprocal edges, average path length, average cluster coefficient, small-worldness, transitivity, and knotty-centredness, are shown in Figure 7.

**Figure 7.**
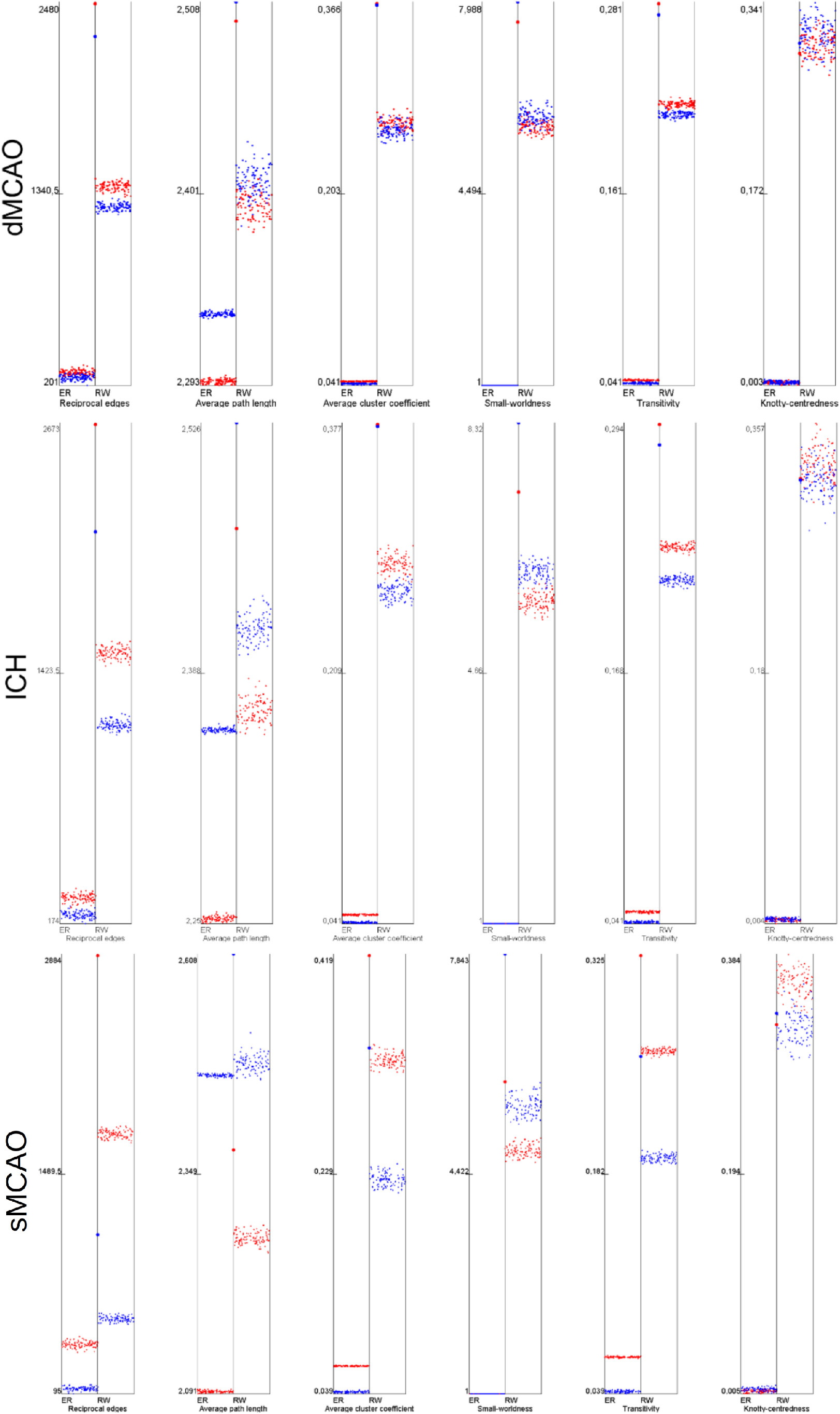
Comparison of 6 global network parameters of the dMCAO, ICH, sMCAO and control connectome. The average path length do not show a larger difference of the control (red large point) and dMCAO lesioned connectome (blue large point). Empirical parameters were tested in the Erdős-Rényi (ER) and the degree preserving rewiring model (RW) for 100 iterations (small dots). The parameters of the empirical ICH-lesioned connectome and the ICH control connectome can be clearly distinguished from randomized connectomes. This distance between the large red and blue points indicates the difference of the sMCAO lesioned connectome and the non-lesioned control connectome. The strongest difference was found for the number of reciprocal edges.

The relationship of the 6 global parameters for the non-simulated control and lesioned connectome appears to be mirrored by the simulated counterparts. For example, the numbers of reciprocal edges, transitivity, and mean cluster coefficients appear larger both in the simulated and non-simulated control connectomes (red), in contrast to the smaller values seen in mean path lengths, small worldness and knotty-centredness in both types of control connectomes. It suggests that the simulated connectomes are equally useful in comparing global network parameters compared to the real connectomes.

The impact of lesion severity seems to have a compounding effect on reciprocal connections and average path length, mean cluster coefficients and transitivity, compared to a proportional effect in other global parameters among the 3 models. It suggests that greater number of lesioned regions disproportionally reduced the reciprocal connections and increased mean pathlength, decreasing the accessibility between any given two regions and thus reducing the efficiency of signal processing and speed of synchronization.

The *mean cluster coefficient* shows similar values for the dMCAO and ICH lesioned connectome in contrast to the great reduction for the sMCAO model, suggesting that the basis of the small world architecture in the sMCAO network has changed significantly, causing the rapid exchange and interaction of bioelectrical signal patterns to decrease dramatically. This is supported by the differences among models in global parameter small-worldness values, suggesting that this network property cannot be protected by the network architectures after lesioning and is better preserved as a value. The transitivity parameter behaves inversely to the small world property, a disproportionally decrease in transitivity of the lesioned connectome was seen in sMCAO compared to ICH and dMCAO models. Since the differences of Knotty-Centredness between control and lesioned connectomes is only minor, this parameter seems to indicate only a small change of densely interconnected regions as network core subsets.

#### Identification of important regions in each model based on centrality parameters

We then performed a local network analysis to determine the importance of a given region with respect to connectivity in each model by ranking the averaged scores over 50 local network parameters (see Methods and Materials). The angular thalamic nucleus, the secondary visual cortex, the submedius thalamic nucleus and the ventral posterior thalamic nucleus were found to have much greater connectional importance in the lesioned dMCAO connectome than in the control connectome. On the other hand, the posterior hypothalamus, median raphe nucleus, and medial preoptic area are most important in the control connectome as well as in the lesioned connectome. The medial agranular prefrontal cortex, posterior hypothalamic nucleus, infralimbic cortex, M1 and ventral pallidum are most important in the ICH control connectome, while the posterior hypothalamic nucleus, median raphe nucleus, ventral pallidum and the medial preoptic area are most consequential with the lowest rank number in the ICH-lesioned connectome. Furthermore, the lateral hypothalamic area (16), medial agranular prefrontal cortex (15) prelimbic cortex (15) and infralimbic cortex (14) were identified to have dense connections between ICH lesioned regions and non-lesioned regions (Supporting information Table 1). The medial agranular prefrontal cortex (M2), the lateral agranular prefrontal cortex (M1), infralimbic cortex, and posterior hypothalamic nucleus have the lowest average ranks in the sMCAO control connectome, compared to the median raphe nucleus, infralimbic cortex, posterior hypothalamic nucleus, prelimbic cortex and the medial septal nucleus in the sMCAO lesioned connectome.

#### The effect of lesion on connectivity and functional network of the models dMCAO model

The afferent and efferent connectivity relationship between individual lesioned regions and regions of the control connectome for dMCAO are shown as 6 *multidimensional circular relationship diagrams* (MCRD) in Figure 8a in sorted order. Following the selection for the top 20% regions with the most importance and connected to at least 4 lesioned regions, the filtered MCRDs are shown in Figure 8b. The analysis indicated that S1 has the most numbers of connections with regions of the control connectome while DI has the least. The regions that were filtered and visualized in the MCRD are arranged selectively in an adjacency matrix to demonstrate there dense interconnectivity (Figure 9b). Each of 23 non-lesioned regions (those regions in Figure 8b that are not labeled with green) is connected with all 6 lesioned regions (Figure 9a), e.g., the perirhinal cortex, rhomboid thalamic nucleus, reuniens thalamic nucleus, S2 and medial orbital cortex, building a core network that are intensively interconnected. In particular, five thalamic, the lateral hypothalamic, mesocortical and orbitofrontal regions show intense interconnections with lesioned regions of the dMCAO model (Figure 9b). Conversely, all 6 lesioned regions are connected to the following learning and memory functional regions, namely the perirhinal cortex (6 reciprocal), lateral entorhinal cortex (6 reciprocal) and postrhinal cortex (5 reciprocal) (Supporting information Table 2), while CA1 has 5 non-reciprocal connections to lesioned regions. On the other hand, all lesioned regions are also connected to motor function regions M1 (6 reciprocal connections), M2 (5 reciprocal connections), SNR, SNC, CPu, STh and pontine nuclei. Details about reciprocal connections of dMCAO lesioned regions and functional defined regions are shown in Supporting information Table 2 and Supporting information Table 3.

**Figure 8.**
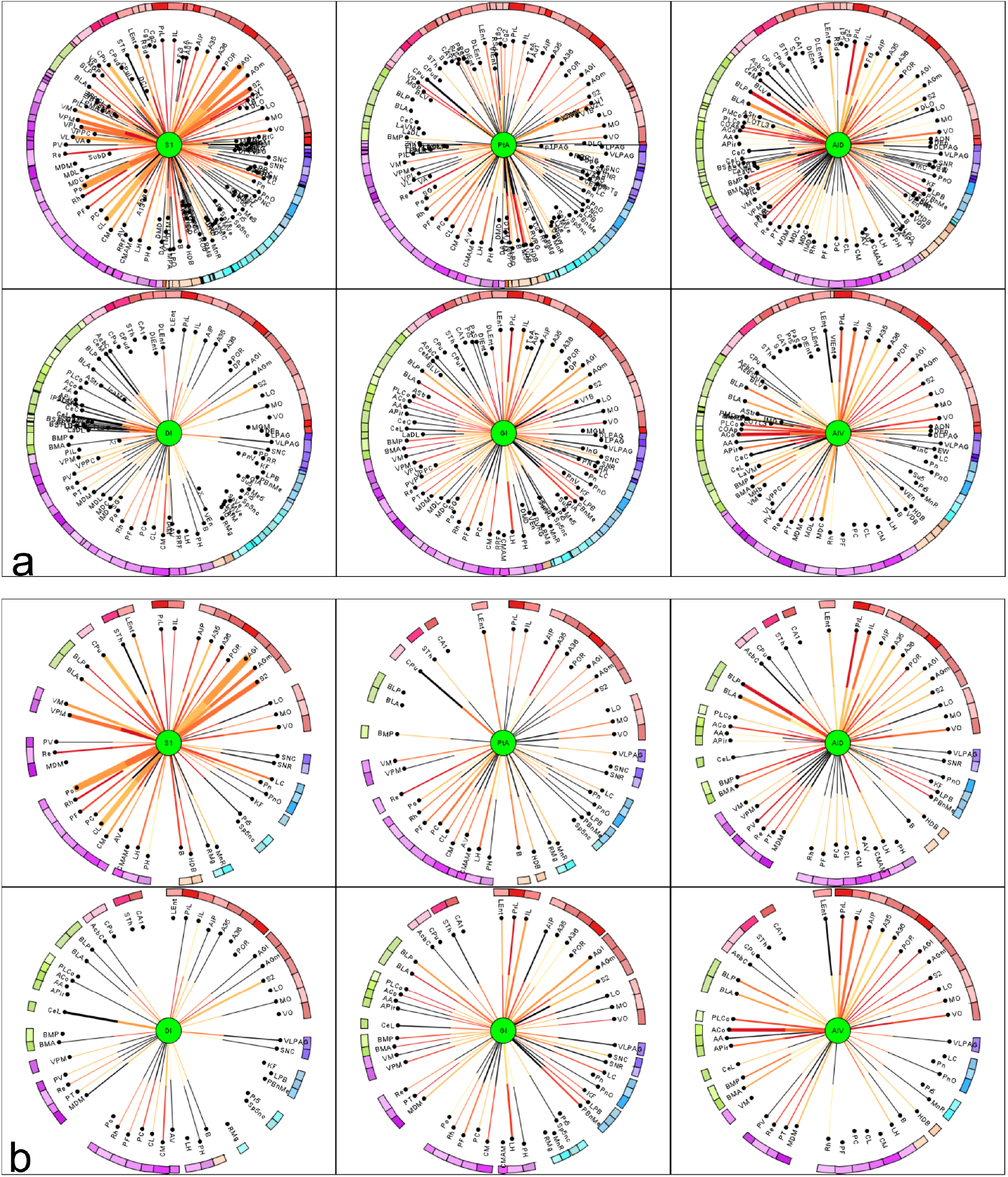
Multidimensional circular relation diagrams (MCRD) of the 6 lesioned regions (dMCAO) and control connectome regions. The lesioned regions were sorted with regard to the number of filtered and connected regions. Center circle: lesioned region. Distance from center: Average rank of 50 local network parameters, Line thickness: number of observations of a connection. Line color: color codes of connection weights. Black line: connection is not reciprocal. Arc length: Number of connections to other lesioned regions. Outer half of a line: connection from the non-lesioned region to the lesioned region, Inner half of a line: connection from the lesioned region to the non-lesioned region. a) The sorted non-filtered MCRD. b) The 20% of regions with highest importance or lowest mean ranks and regions which are connected to at least 4 lesioned regions.

**Figure 9.**
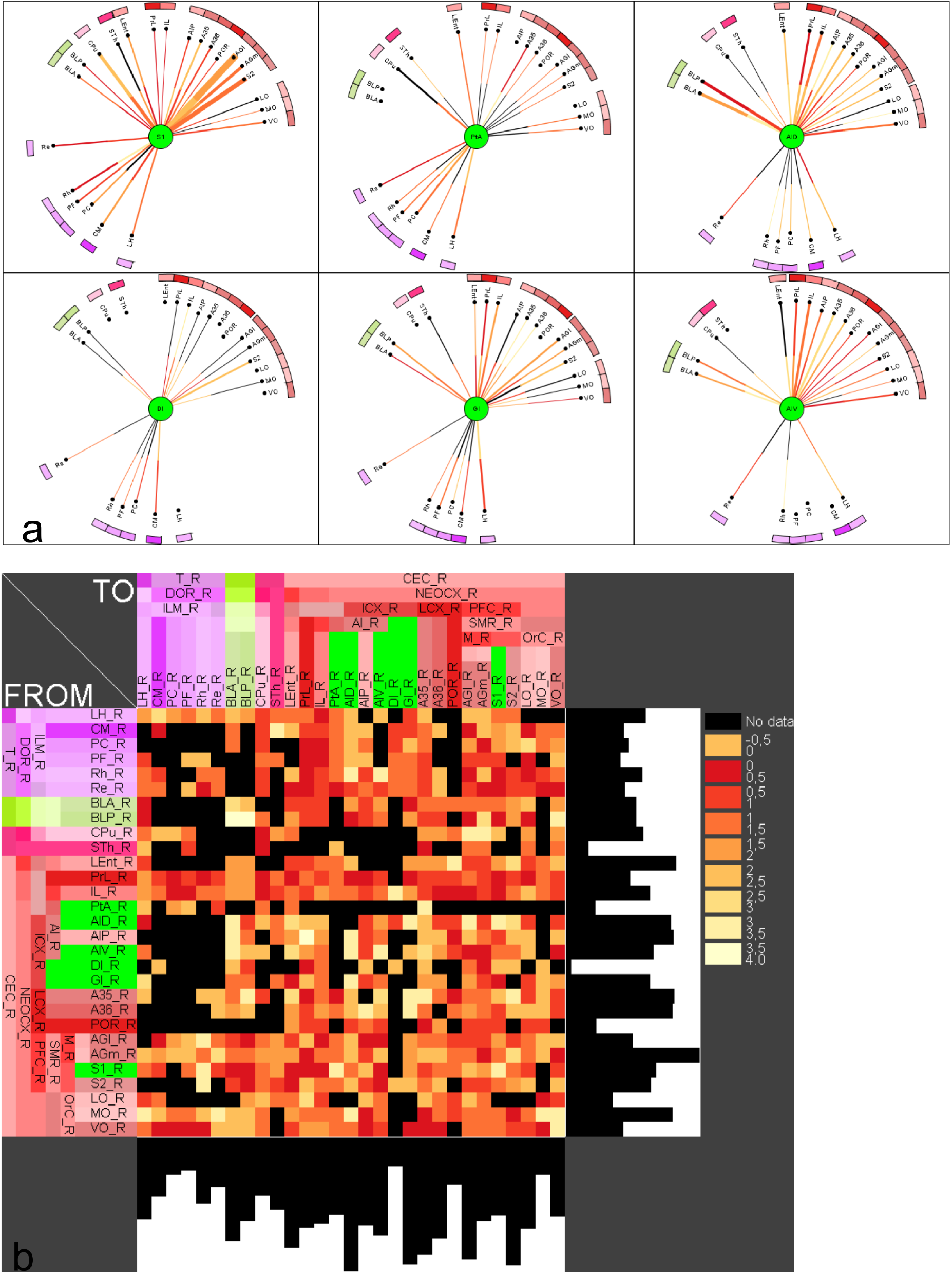
Filtering of most important regions which are connected with dMCAO lesioned regions. a) Remaining non-lesioned regions with direct reliable connections to the 6 lesioned regions following 3 parameter filtering. Distance from center region (lesioned region) is the average rank of local network parameters filtered for the 20% of lowest ranks (largest importance for the network), connections to all other lesioned regions (6) and 2 observations per connection. The lesioned regions were sorted with regard to the number of filtered and connected regions. b) Adjacency matrix of lesioned regions and non-lesioned regions of the dMCAO experiment.

#### ICH model

The numbers of links between the 22 ICH lesioned regions and non-lesioned regions are shown in the complete (Supporting information Figure 2) partially filtered (Supporting information Figure 2) or filtered MCRD (for top 20% important regions) (Figure 10), respectively. The lesioned regions Ce, BL and CPu are subcortical nuclei with a large number of connections to different subcortical and cortical regions, critically affecting information processing. The number of connections between lesioned regions and functionally defined regions as well as control regions without a functional definition are shown in Supporting information Table 5. Lesioned regions and with the most abundant connections with the functional regions are CL (11), B (10), PC (19), MDL (10), VM (10), BLA (9) and CPud (9). There were more connections found between the lesioned regions and motor regions (6-15 connections) than with learning regions (6-7 connections) (Supporting information Table 4), supporting that ICH lesion may lead to significant motor impairment. The CA1 region has 6 non-reciprocal and 1 reciprocal connections with the lesioned regions.

**Figure 10.**
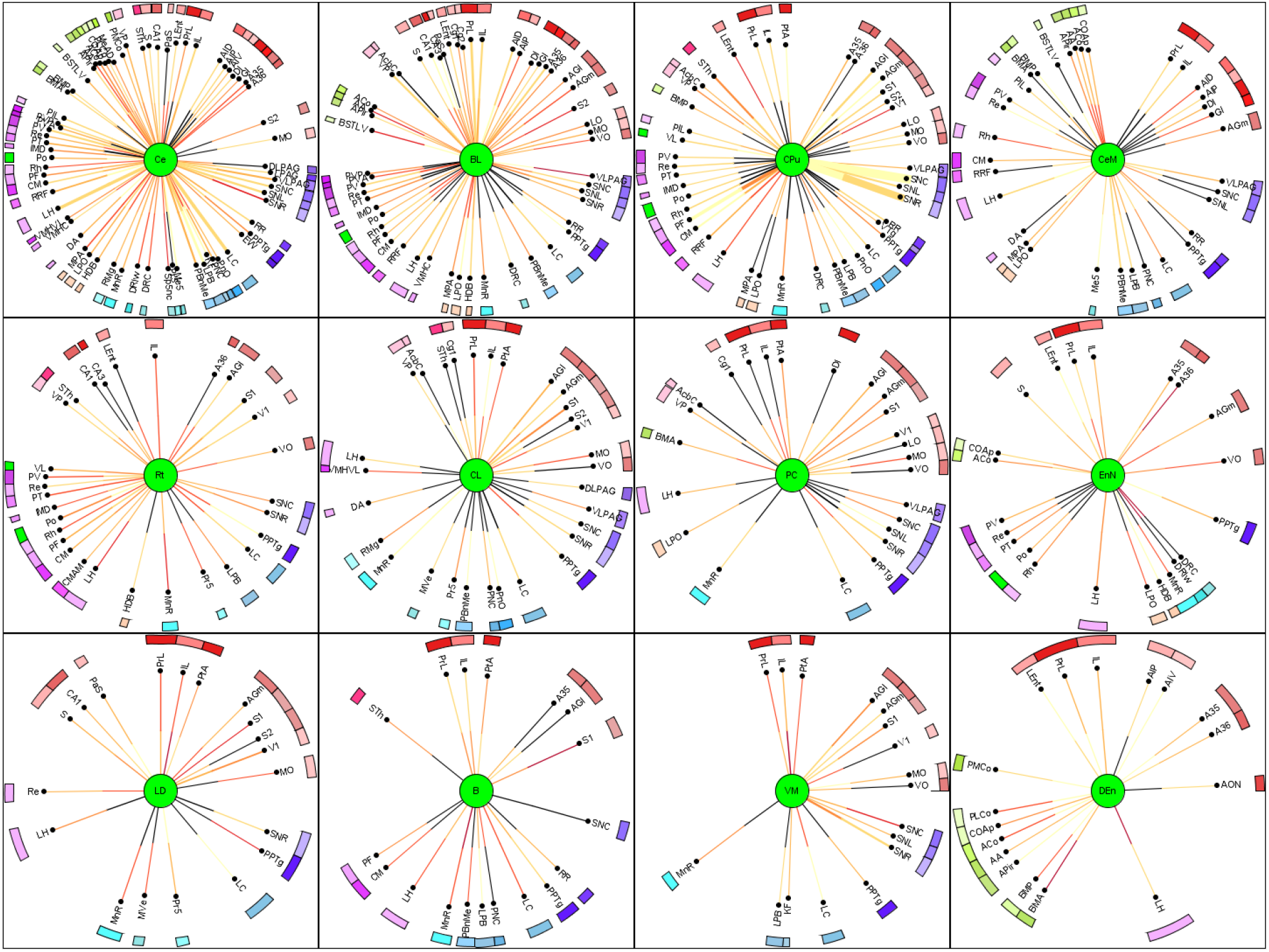
MCRD of 12 from a total of 22 ICH lesioned regions and control connectome regions. The lesioned regions were sorted with regard to the number of filtered and connected regions. For a more detailed description see Figure 8. The lesioned regions are in the centers of the MCRDs (green filled circles). The same filter condition as in Figure 8b was applied: The 20% of regions with highest importance or lowest mean ranks and regions which are connected to at least 6 lesioned regions and 2 observations per connection.

#### sMCAO model

The complete and filtered MCRD representations of the connections between 57 sMCAO lesioned regions and non-lesioned regions are shown in the Supporting information Figure 4 - Figure 6 and in Figure 11, respectively. The diencephalic lateral hypothalamic nucleus appears to be the most important region in terms of network architecture followed by the amygdalar regions BST and Ce and the thalamic zona incerta. The lesioned regions with most connections are connected to several identical contralateral non-lesioned regions, suggesting that the function of latter regions is strongly affected by sMCAO (Supporting information Table 6).

**Figure 11.**
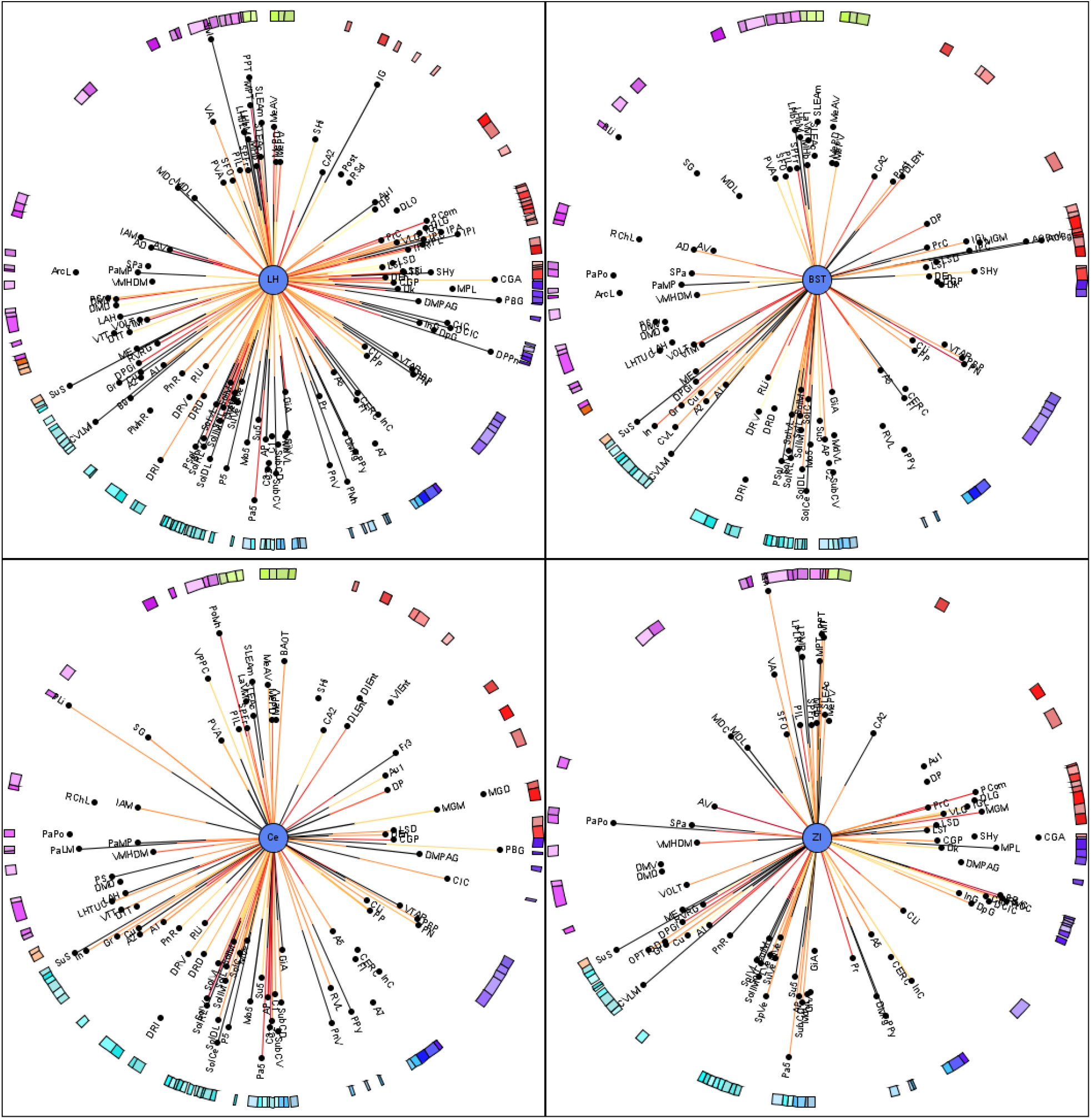
A filtered MCRD visualization of sMCAO lesioned regions which share most connections and their relations to control connectome regions. Distance from center region (lesioned region) is the average rank of local network parameters filtered for the 80% of lowest ranks (largest importance for the network). The lesioned regions were sorted with regard to the number of filtered and connected regions. Regions without lines indicate connections to or from the lesioned regions in the center by one of their subregions.

Five lesioned regions, namely the ventrolateral thalamic nucleus (VL), caudate putamen (CPu), cingulate cortical area 1 (Cg1), cingulate cortical area 2 (Cg2), perirhinal cortex (A35) also belong to a functional group. Therefore, these functionally defined regions were removed from the functional groups of motor and learning processing. The sum of connections of the set of lesioned regions and motor regions is the greatest for the substantia nigra pars compacta (42), followed by subthalamic nucleus (31) and substantia nigra reticular part (31). The CA1 region has 48 connections followed by the lateral entorhinal cortex (45). The maxima of connections (input and output connections) of lesioned regions are found for the infralimbic cortex (56), the prelimbic cortex (55), CA1 (48), reuniens nucleus (46) and rhomboid thalamic nucleus (43) among important primary cortical regions (primary visual cortex, orbital cortices) interconnected through more than 20 links with lesioned regions. This pattern of connections between learning regions and primary cortex areas suggests that specific learning functions of the visual system (especially secondary visual cortex) and regions controlling value-based decision making (such as striatum and ventromedial prefrontal cortex) are particularly affected functionally in sMCAO stroke as shown in the reduced adjacency matrices Figure (6).

The number of connections from lesioned regions to motor or learning regions is 248 (33%) or 507 (67%) for sMCAO, compared to 75 (61%) or 48 (39%) in ICH, reflecting the dominant nature of functional impairment afflicted by each type of injury. Consistent with this metric, we found that the connectivity pattern between lesioned regions and the regions processing learning behavior (40) have more frequently larger *CMI_All_* (connectivity matching index) values (0.3058 *±* 0.0864) than those with the motor behavior regions (21) (0.2776 *±* 0.0885). However, the sMCAO model still has a greater impact on overall connectivity loss as the connections between the lesioned and non-lesioned regions is 35%, compared to 13% for the ICH model and to 1.13% for the dMCAO model.

#### Relationship between dMCAO lesioned regions and functionally defined regions

The similarity of connections between lesioned regions and functionally defined regions (motor or learning function region) was performed by calculating the connectivity matching matrix for input and output connections (*CMI_All_*) as described in Methods and Materials. The FitzHugh-Nagumo (FHN) neuron model was also applied to the connectome data accordingly [211]. The FHN-coactivation matrix was used in the same way as the *CMI_All_* matrix for the comparison of FHN-coactivations and connectivity matchings of pairs of regions. For example, subparafascicular thalamic nucleus rostral part and PtA have a *CMI_All_* value of 0.222 and a FHN-coactivation value of 0.172 as shown in the first row of Supporting information Table 7, which further demonstrate the comparison of each pair of a non-lesioned motor or learning region (1 row) and a lesioned region with regard to 5 parameters (graph distance D for output Distance *D_out_* and input distance *D_in_*, Euclidian distance *D_spat_*, *CMI_All_*, *FHN*) shown in 5 columns.

To find out which learning regions share a high similarity in connectivity with lesioned regions, data from Supporting information Table 7 was first sorted for the functional markers (2: motor, 3: learning), followed by secondary sorting with respect to the spatial distance between the functional learning group and the motor group (*D_spat_*) of the PtA R region, to determine the connection similarity between lesioned and functional regions (Supporting information Table 8).

The same applies to the similarity of the activation dynamics if sorted by FHN rank (Supporting information Table 9). As a result we found that, the postrhinal, perirhinal and cingulate cortex (CG1) displayed the greatest similarities in connectivity to PtA R. After sorting the *CMI_All_* values of the lesioned PtA region it turns out that perirhinal cortex (Peri), CG1 and LEnt have the largest *CMI_All_* values (0.588, 0.482, 0.462). Interestingly, the LEnt is connected with all lesioned regions, and the most similar input and output connections when compared with each of the 6 lesioned regions.

Not surprisingly, LEnt is the major source for the trisynaptic circuit through the hippocampal learning system. CG1 has a much larger coactivation value of 0.32 in the FHN matrix in comparison with Peri and LEnt (maximal coactivation is 0.533 for the region pair PtA-Subiculum). By sorting the coactivation matrix values of the coupled FitzHugh-Nagumo neuron simulation of Supporting information Table 7, a hippocampal pattern of coactivations was found among the lesioned regions. S1, PtA, AID and AIV are strongly co-activated with CA1, S, Par and Pre. DI and GI are strongly co-activated with CA1, S, Par and Pre. This is a further indication of considerable involvement through lesioned regions and their connections with regions functionally embedded in spatial learning processes. Following sorting the *CMI_All_* values of the two functional sets (motor behavior, learning behavior) and dMCAO lesioned regions, highly ranked pairs were found for lesioned regions and both functional groups with rank number smaller than 50 (out of maximum rank of 242) (Supporting information Table 9). One top ranked pair between the perirhinal cortex and the lesioned PtA indicate that PtA has the largest connectional similarity (sharing connections with same source and target regions) with the non-lesioned perirhinal cortex region. A similar trend was also found by sorting the ranks of the FHN coactivation matrix accordingly (Supporting information Table 10), pairs of lesioned regions and either functional region came up among the top 50 ranked groups. Thus, based on structural connectome analysis using *CMI_All_* and functional FHN model, dMCAO preferential affects brain regions crucial for motor and learning functions.

#### Relationship between ICH lesioned regions and functionally defined regions

*CMI_All_* and the *FHN* coactivation matrices were used to identify the functional regions with extensive connections with the lesioned regions (Supporting information Table 1 and Table 11). We found that the lesioned anterior basolateral nucleus shares similar connections with motor region medial agranular prefrontal cortex per *CMI_All_* rank, and with the laterodorsal thalamic nucleus generalized topology matrix (GTOM) which estimates the normalized counts of the number of m-step neighbors that a pair of nodes share [206]. Additionally, the laterodorsal thalamic nucleus showed a large coactivation of the FitzHugh-Nagumo neuron simulation [211] with the lesioned central amygdaloid nucleus medial division. The connectivity similarity (connectivity matching) of lesioned regions with functional regions is also high as evidenced by the high ranks of *CMI_All_* and *FHN* values, between 1 and 61.

#### Relationship between sMCAO lesioned regions and functionally defined regions

The lesioned ectorhinal cortex was identified as the most highly connected lesioned region per *CMI_All_* ranking with 6 different learning regions, followed by anteromedial thalamic nucleus (rank 2), the piriform cortex (3) and the cortical amygdaloid nucleus (rank 5). Whereas the lateral globus pallidus (1), the cerebellar nuclei (2), M1 (6) and the zona incerta (9) had rich connections with motor regions.

Likewise agranular insular cortex (1), the basal nucleus of Meynert (1), the posterior basomedial nucleus (2), the nucleus of the vertical limb of the diagonal band (2), the primary somatosensory cortex (2), the dysgranular insular cortex (2) and the secondary somatosensory cortex (2) were the top lesioned regions highly connected to learning regions based on *FHN* -simulation coactivation values.

Apart from *CMI_All_* and *FHN* coactivation, we found that the mean graph theoretical distance of lesioned regions to motor or learning functional regions. The mean functional distance of lesioned regions to motor or learning functional regions to be 7.8/8.2 (Input/Output) or 13.1/12.9 (Input/Output). This suggests a stronger impact of sMCAO lesion on learning than motor function.

As a proof of principle approach to confirm the learning impairment caused by sMCAO, we performed a homogeneous spiking population simulation of the sMCAO control and the lesioned connectome. For each region, 500 leaky integrate-and-fire (LIF) neurons with 20% of inhibitory and 80% excitatory neurons were modeled using the NEST simulator [79,129] in *neuroVIISAS* [157] (total of 229116 LIF neurons). The synaptic delay was set to 1 ms, the locus coeruleus was stimulated by 10kHz and simulation time was 300 ms. We found that the mean coefficient of variation of spike intervals (*CE_isi_*) increases in CA1, CA3 and DG after sMCAO, from 0.027, 0.071 and 0.098 in the control connectome to 0.183, 0.282 and 0.287 in the sMCAO lesioned connectome, respectively. In addition to the hippocampal core areas, regions that are integrated into the functional circuits of memory and learning like the mammillary body and the subiculum also increased their *CE_isi_* from 0.004 and 0,026 to 0.047 and 0.132, respectively. This dynamic changes of the sMCAO lesioned network is consistent with the hypoactivation as marked by reduced Fos immunoreactivity in lesioned animals comparison to sham operated animals during functional activation (Figure 5).

In conclusion, the simulation revealed an increase of interspike variability of learning and motor behavior regions. Similar effects have been described in lesion studies and neurophysiological measurements of interspike intervals [41, 58, 59, 208].

#### Similarity of functionally defined regions with lesioned regions in each model

By determining the similarity of connections, coactivation or cross-correlation patterns between the lesioned regions and functional groups or non-lesioned regions, we assessed the associations between lesioned regions and functions. The similarity between lesioned and functional regions was computed for the structural feature *CMI_All_* values of the connectomes and for the dynamic feature of mean coactivations between lesioned and the functional groups. The CMI of input and output connections of a pair of regions were first compared against *CMI_All_* values of all pairs of regions, and the significance determined by the Student’s T-test. Since each group is composed of several regions, the mean *CMI_All_* values of a lesioned region were calculated with all regions of a functional group and the results were show in Supporting information Table 12.

The FHN dynamics has been performed by applying 500 iterations with a step size of 1.0 using weighted connections, generating cross-correlation matrix used for the multiple subset statistics in *neuroVIISAS*. Consistent with the *CMI_All_* similarity values, FHN model model also revealed a greater similarity between learning regions and lesioned regions of dMCAO and sMCAO than between the motor regions and the lesioned counterparts. To the contrary, the ICH lesioned regions showed a greater similarity to motor regions for both *CMI_All_* values and the FHN model. The similarity analysis affirmed a stronger impact of dMCAO and sMCAO lesion on learning function, contrary to the stronger impact of ICH on motor function.

#### Comparison dMCAO and ICH

In contrast to the apparent discrepancy in lesion size, the extent of functional impairment between the dMCAO and ICH models are comparable. We thus compared the relative loss of neuronal connections due to gray matter destruction between the dMCAO and ICH groups. This was done with respect to the relative loss of connections of the functionally defined regions in the control connectome after removal of the lesioned regions. The relative loss of neuronal connections of motor regions averaged 5.63% and cognitive regions 5.6% for dMCAO, compared to 33.9% and 10.97% for ICH. With respect to the loss of reciprocal connections, 4.18%/4.3% for dMCAO relative to 35.35%/11.19% for ICH for the motor/learning systems. Based on these analyses, the ICH model suffers from a bigger relative loss of connectivity between lesioned and functional regions compared to the dMCAO model, likely to account for the severe functional impairment relative to its lesion size. This clearly indicates that ICH has a greater impact on motor than learning function compared to the dMCAO stroke, which is supported by the behavioral data.

#### Dynamical modeling of CA1 and DG comparing control and lesioned connectomes predicts hippocampal functional impairment

Stroke by dMCAO led to impairment of hippocampus-dependent function and brain oscillations [70], we first determined whether dMCAO-induced structural network lesion has dynamic effects on spike propagation in remote brain regions like the CA1 or DG in a directed weighted bilateral connectome of a simple network consisting of a total of 40 regions as shown in the matrix (Figure 12) following the removal of 6 lesioned regions in the ipsilateral hemishphere using 3 dynamic models; namely the coupled FHN model (membrane potentials), the Wilson-Cowan neural mass model (inhibitory and excitatory interacting populations) and the Mimura-Murray reaction-diffusion model. The overall amplitude level of the propagation waves of CA1 region is consistently smaller than that of the DG region in both the control and lesion connectome detected in the FHN model, while a slight increase of membrane potential was detected from 30 in control- to 36 in lesioned connectome, resulting in an increased periodicity in spike propagation waves in both CA1 and DG compared to control (Figure 12). In addition to the change in the frequency of CA1 and DG waves, a slight phase shift of the CA1 region after about 20 iterations in the control connectome was noticeable, in contrast to the nearly synchronous waves in two regions in the lesioned connectome.

**Figure 12.**
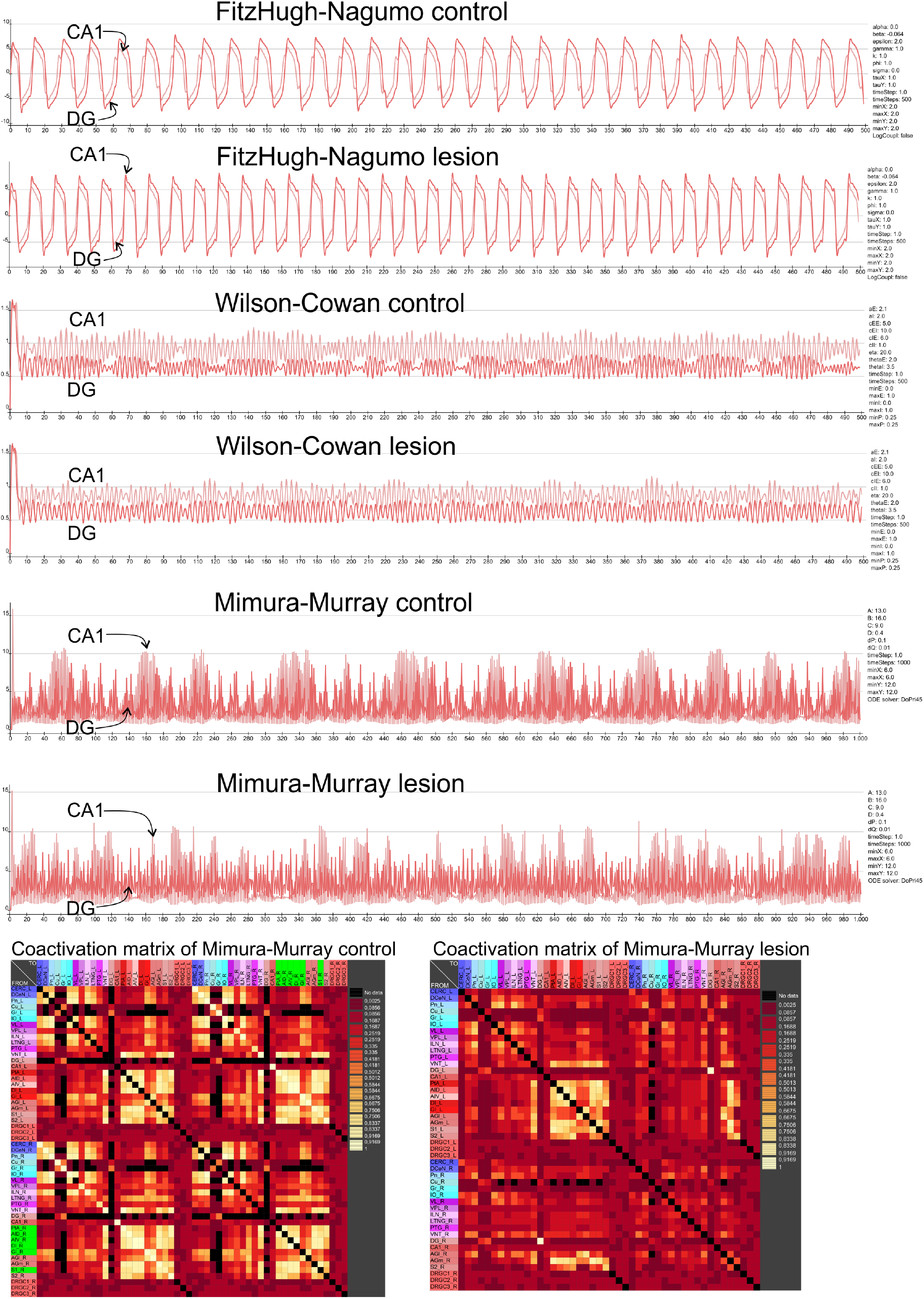
Dynamics of FHN, Wilson-Cowan and Mimura-Murray models in control and lesioned connectomes. The dynamics of the hippocampal CA1 and DG regions which are not directly concerned in the dMCAO models are shown. The coherence of diffused concentrations are shown for the Mimura-Murray reaction-diffusion model. Parameters of the models are shown on the right of the diagrams. They were hold constant for the control, resp., lesion models.

The Wilson-Cowan neural mass model reflects connectome activity as spindle shaped oscillations with variable amplitudes. The lesioned connectome oscillations showed a larger variability but with smaller overall amplitudes in the CA1 region compared to control connectome. In the DG, compared to the alternating large and small amplitude spindle clusters in the control connectome, the small amplitude-spindles seemed to diminish in the lesioned connectome.

In the Mimura-Murray reaction-diffusion model the control connectome CA1 showed clusters of large amplitude spikes in greater periodicity compared to those in lesioned connectome. In contrast, the lesioned connectome showed less pronounced clustering with a blend of random large and small amplitude spikes in CA1 and DG regions.

In general, the CA1 clusters lasted longer and had larger amplitudes compared to those of the DG region. The general dynamical tendency of the spikes of the Mimura-Murray model in the DG region seems to resemble the distinct patterns of spindles in the Wilson-Cowan model, providing converging evidence that connectivity changes after dMCAO may have altered the spike propagation in the hippocampus.

Since the Mimura-Murray reactions-diffusion model showed the greatest extent of differences in dynamic signal propagation in CA1 and DG between control and lesioned connectome, this model was selected to compare the three lesion models with identical model parameters (Figure 13). The locus coeruleus with its very extensive cortical connections was chosen as the region from which the initial signal excitation originates.

**Figure 13.**
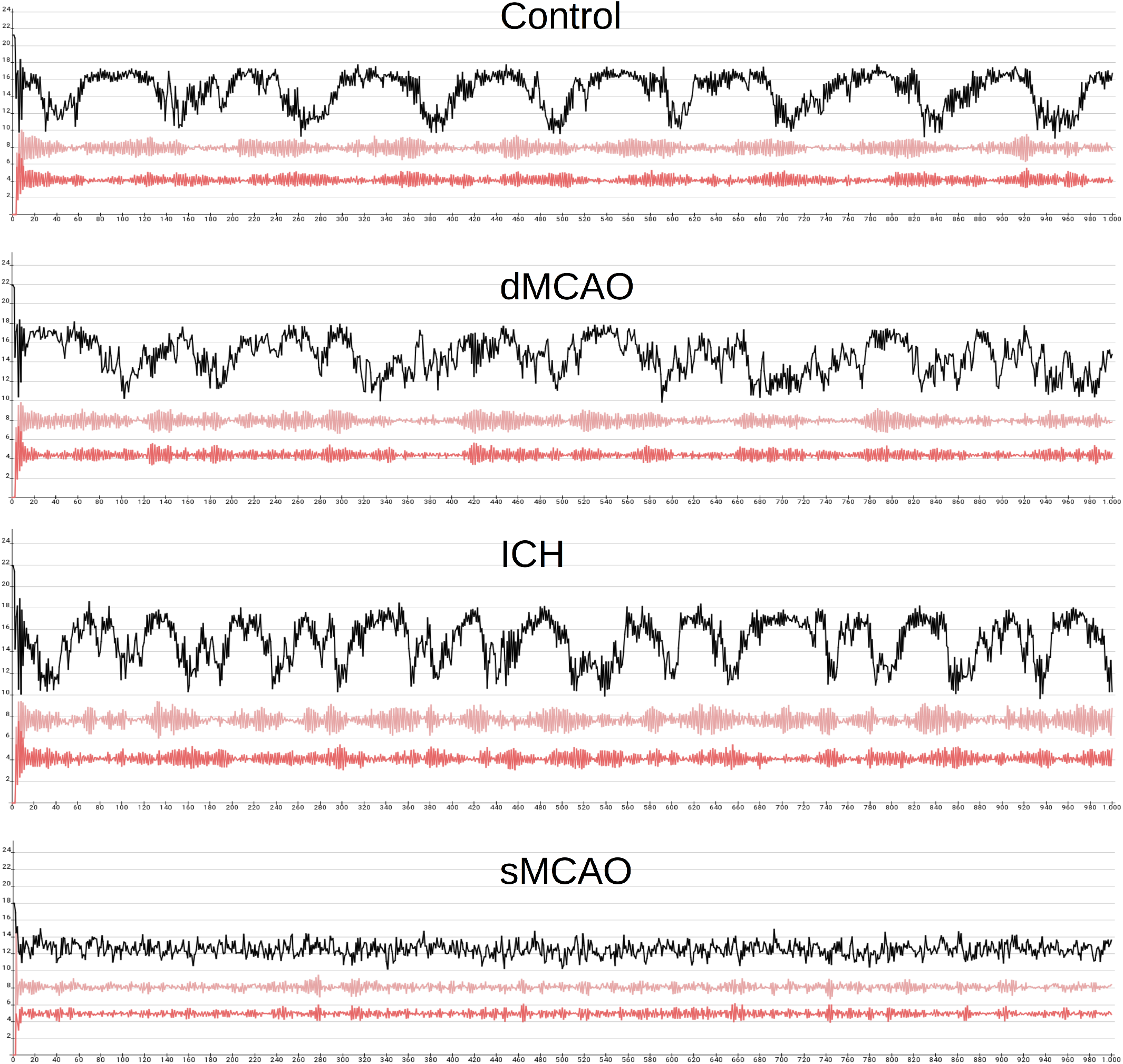
Mimura-Murray simulation of control and lesioned connectomes. The black curve depicts the Kuramoto index. Light red curve corresponds to CA1 and the dark red curve to DG.

For the simulation, the same parameters were used for the control, dMCAO, ICH, and sMCAO connectome, respectively: A: 13.0, B: 16.0, C: 9.0, D: 0.4, dP: 0.1, dQ: 0.01, minX: 6.0, maxX: 6.0, minY: 12.0, maxY: 12.0 timeStep: 1.0, timeSteps: 1000 and the DoPri45 ODE solver. The global dynamic behavior of the non-lesioned and lesioned connectome can be characterized by the average activation, average coactivation and Kuramoto index, among which the latter two measures also provide estimates of the extent of signal coherence and signal similarity, respectively. The Kuramoto index, also known as the Kuramoto order parameter, calculates the synchronicity of the CA1 and DG function curves [164, 214]. In the control connectome, the Kuramoto index curve displayed alternating phases of high and low synchronicity between DG and CA1, indicating a periodic pattern of coherence between the two regions. Both the CA1 and DG signals also exhibit alternating phases of large and low amplitude spindles, in which the high amplitude clusters have a duration of about 60-80 iterations and usually correspond to the high Kuramoto index values.

dMCAO lesion appeared to disrupt the synchronicity between two regions as reflected by the irregular pattern of Kuramoto index curve. Although the alternating phases of large and low amplitude spindles were largely preserved in the CA1 and DG signals, the maximal amplitude has increased in the CA1 spindles. In addition, there were both a decrease in the coherence measures average coactivation and Kuramoto index and a general decrease in overall connectome activity, consistent with the cognitive impairment detected in the dMCAO model. Compared to dMCAO, the ICH simulation showed a more synchronized connectome activity between CA1 and DG based on Kuramoto index curve, despite shorter but more frequent synchronous phases relative to those of control connectome. Large amplitude clusters of spindles in both CA1 and DG are also more frequent but shorter with increased maximal amplitude compared to control connectome. The maximal amplitude of CA1 spindles were also larger than that of the control. The Kuramoto index curve of sMCAO completely lacked alternating phases of high and low synchronicity as in the control or ICH connectome simulations. Also, the local maxima and minima of the Kuramoto index curve are significantly lower than in the ICH or control connectome case. The typical high-amplitude intercept and low-amplitude gap pattern is no longer evident for the CA1 and DG regions (Figure 13). The decrease in signal coherence both for the Kuramoto index and average coactivation as well as the mean connectome activity were somewhat more pronounced in the ICH model relative to dMCAO model, yet most severe in the sMCAO model. Overall, the results of the CA1 and DG in 3 stroke models using the Mimura-Murray dynamic simulation are consistent with the extent of functional impairment displayed for the Barnes maze learning and spatial exploration.

## Discussion

Stroke is known to change brain activity [87], structure [149] and induce reorganization [167], yet how stroke affects functional activation at the level of neuronal networks is not well understood [173]. We compared the degree of functional impairment and mapped the changes of connectivity patterns among three commonly used experimental stroke models, namely the sMCAO, dMCAO and ICH models with progressively smaller lesion sizes but with distinct lesion locations [5, 110, 142, 155, 156, 183]. The sMCAO model consistently exhibited the biggest lesion size and most severe functional impairment with the greatest number of lesioned regions distributed in both the cortical and subcortical regions, resulting in the greatest change in structural and functional connectivity. Interestingly, despite a 3.5-fold smaller lesion size in the ICH model relative to dMCAO model, the former displayed a comparable if not greater degree of functional impairment compared to the latter, albeit with higher number of lesioned regions exclusively in the subcortical locations. This is consistent with the fact that subcortical regions are generally ranked with greater importance than cortical regions according to the analysis based on local network parameters. Further, all three models produced impaired function involving regions remote from ischemic injury, highlighting the contributory role of connectivity loss in functional impairment. Connectome analysis also revealed that brain regions corresponding to learning and motor behavior exhibit structural connection changes and altered dynamics during functional simulation following the removal of regions and their connections in the investigated connectomes. The lesioned regions seem to have a greater structural connection and dynamic relationship with the functional groups compared to intact regions. Most importantly, via the connectome informatics, we have identified key brain regions affected by the lesions of each injury model, providing complementary information in predicting how function might be affected in each type of stroke.

Both global and local network parameters [148] were used to analyze structural connectivity changes. Following differential connectome analysis with 6 chosen global network parameters, the number of reciprocal connections was found to be increasingly reduced in the order of dMCAO, ICH and sMCAO models, in which a massive decrease of 59.8% of reciprocal connections was detected in the lesioned sMCAO connectome. In contrast, the small-worldness [14] was progressively increased among the 3 stroke models, with the biggest increase for the sMCAO by 34.7%. Although there are only 1.03 more connections loss in the ICH model than in the dMCAO model, the mean path length of the former has increased by 5.3 times, hence the accessibility of the regions to each other become much reduced. Similarly, lesioned regions in sMCAO quadrupled when compared to those in ICH, but the reduction of the cluster coefficient was about 27 times. Thus, there is no linear relationship between the number of lesioned regions and the global parameters determined here, which could be attributed to damage of specific connections with particularly important topological network properties. To identify these structurally important regions, average rank of each region was determined in the 3 non-lesioned and lesioned connectomes using an array of 50 different local network parameters including centrality measures, Shapley index and the Katz index [159], means of coactivations of regions derived from dynamic models. We found that in general subcortical regions ranked higher (with low rank number) in importance than cortical regions. In the case of non-lesioned dMCAO connectome, subcortical nuclei including diencephalic nuclei, angular thalamic nucleus, the submedius thalamic nucleus and the ventral posterior thalamic nucleus were deemed most important with regard to network architecture and formed intense connections with the cortical regions. In the sMCAO control connectome the regions M2, M1, IL and posterior hypothalamic nucleus ranked the highest, in parallel with median raphe nucleus, IL, posterior hypothalamic nucleus, prelimbic cortex and the medial septal nucleus in the sMCAO lesioned connectome. The shared top ranked regions of IL and posterior hypothalamic nucleus between the control and lesioned connectome reflects the existence of stability for important regions of brain networks.

To further identify regions most affected by each model of stroke, MDCR diagrams were first filtered with at least 4 connections between the non-lesioned regions and two different lesioned regions to reflect a strong multiple lesion effect, followed by ranking for importance in network architecture. We found that the lesioned regions in the dMCAO model had stronger and reciprocal connections with other cortical regions compared to subcortical or diencephalic nuclei, projecting a stronger dysfunction at the cortical level. In contrast, in the ICH model more regions with strong connectivity to lesioned regions are located within subcortical and diencephalic regions rather than within the cortex. Unlike in the dMCAO model, the lesioned regions that are strongly interconnected in the ICH model are distributed much more diffusively in brainstem, mesencephalic, diencephalic and subcortical as well as cortical regions, yet the densest connections of lesioned areas in the ICH model and non-lesioned areas lie in the lateral hypothalamus, medial agranular prefrontal cortex, prelimbic cortex and infralimbic cortex. Similar to the ICH model, lesioned regions of sMCAO with the densest connections such as the LH, BST, Ce and ZI also belong to subcortical nuclei and diencephalon.

To fine tune how function of regions or neuronal ensembles are affected by stroke, an approach similar to a more precise micro-mapping of functional topography as proposed recently by [89] were applied to the regions of interest in this study. By interactive filtering and visualization in MCRD diagrams there revealed numerous connections of lesioned regions with the two sets of functionally defined regions for motor or spatial learning behavior. By analyzing ranks of similarities of *CMI_All_*, GTOM and FHN coactivation matrices, one can predict how function is affected in each model [12, 29, 35, 50, 80, 104].

It appears that the ICH lesion has a more distributed lesional effect to many different diencephalic and subcortical relay regions with key functions of signal transmission to higher level cortical regions. For example, most connections were detected for the ICH lesioned amygdalar region CeM with control regions [161]. Another lesion region VPL with many connections to the control regions lies in is an important relay nucleus in the thalamus for peripheral sensory signal transmission. It suggests that these polyfunctional areas with their subareas may induce multiple fine granular behavioral changes due to the known regulatory microcircuits as well as massive subregional, regional and supraregional intrinsic connectivity, which were not captured by the type of behavioral tests performed here. Interestingly, based on the connection similarity comparison approach with *CMI_All_* values, ICH lesions demonstrated a greater similarity with the motor than with learning regions.

In contrast, the mean *CMI_All_* values of dMCAO lesioned regions are larger for the set of learning regions than for the set of motor regions. A comparable result was obtained by analyzing the coactivation pattern of a FHN simulation [37, 57, 84, 119, 121, 135]. Thus, it can be predicted that a dMCAO lesion is very likely to lead to learning disorders per lesion-to-functionally defined region relationship. From the perspective of connectomics, the dMCAO lesion has stronger remote effects on the learning system rather than on the motor system. In support of the effect of dMCAO on remote region involved in learning function, we have detected reduced hippocampal activation following exploration of a novel environment in CA1, CA3 and DG, indicating changes in spatial memory processing [155, 156].

In the sMCAO model, the lesions are so extensive that even regions assigned to specific functional motor and learning groups of the control connectome were destroyed. Thus, pronounced impairment in both motor and learning function can be predicted. A large number of connections between lesioned regions and motor regions was found in the sMCAO model for the subcortical nuclei SNC, STh and SNR, suggesting that these core regions of the basal ganglia seem to be particularly affected by the sMCAO stroke. Yet there are almost twice as many connections between sMCAO lesioned regions and the learning regions are compared to those between lesioned regions and the motor regions, with particularly dense connections to CA1 and LEnt and the thalamic nuclei. Thus, from the perspective of connection structure, the sMCAO lesion seems to also affect the learning regions more than the motor regions.

Apart from changes in the structural network parameters, we detected changes in the signal dynamics in the CA1 and DG regions when the nonlinear Mimura-Murray reaction diffusion model [163] was applied to the control and lesioned connectomes from a well-defined source region [108] or modeled neuron population [13]. Unlike the control connectome that showed a relatively stable periodicity of alternating high and low-amplitude oscillations, incrementally reduced periodicity was found in the lesioned connectome of ICH and dMCAO showed, or a complete lack of periodicity in sMCAO, correlating with the extent of damage among the 3 injury models. However, although ICH had the least reduction in synchronicity between CA1 and DG, the number of synchronous phases for both functional curves of CA1 and DG increased compared with the control connectome. The relationship between the oscillatory changes in the dynamic network model applied here and real-life neurophysiology remains unclear to us at the moment, further investigation is warranted to better understand the causality between network change and oscillatory changes in different models. For this purpose, future investigation of time-dependent relationships in the synchronization behavior of all regions of a connectome using Granger causality analysis [15, 25, 67, 171, 207] may provide a deeper understanding.

One caveat of our study is that we have only performed the connectome analysis assuming a complete destruction of the identified lesioned regions. However, there could be various extent of survived neurons particularly in larger anatomical regions. Another limitation is that the *neuroVIISAS* connectome is restricted to information in the grey matter. Future fine tuning is warranted taking into consideration of the extent of neuronal loss within each region and white matter connectivity. A third limitation is that we derived the dynamic simulation of spike propagation by assuming a universal percentages of excitatory and inhibitory neurons for all brain regions in the connectome. A further refinement of the percentages of excitatory versus inhibitory neurons in each brain region in the connectome may improve the precision of the dynamic modeling of spike propagation in the hippocampus regions. In conclusion, the three models of stroke investigated led to varying lesion sizes that do not correlate with the extent of functional impairment assessed by behavioral tests. Using connectomics, our study provides insight into the effects of lesion location and connectivity loss on connectome architecture with respect to motor and learning systems. Our proof of principle study based on structural and functional connectomics approach offers a viable platform in predicting stroke induced functional impairment and identifying remote brain regions along the connection pathways crucial for motor and learning functions.

## Acknowledgments

Acknowledgements: This work was supported by NIH grant R01NS102886 (JL), R21NS120193 (JL), Research Career Scientist award IK6BX004600 (JL).

## Authores Contributions

OS contributes to the writing and interpretation connectome analyses, overall concept deveopment. PE contributes to the connectome analysis and modeling.

YW contributes to the stroke models, behavioral and histological assessment.

AK contributes to the identification of lesions in the connectome based on histological characterization. GR contributes to the identification of lesions in the connectome based on histological characterization. JL contributes to the overall concept, interpretation of data and manuscript writing and editing.

## Supporting Information

### 1 List of Abbreviation

4: Trochlear nucleus
6: Abducens nucleus
10: Dorsal motor nucleus of vagus
12Sprin: Principal hypoglossal nucleus
3PC: Oculomotor nucleus parvicellular part
5Sol: Trigeminal solitary transition zone
7DI: Facial nucleus dorsal intermediate subnucleus
7DL: Facial nucleus dorsolateral subnucleus
7DM: Facial nucleus dorsomedial subnucleus
7L: Facial nucleus lateral subnucleus
7VI: Facial nucleus ventral intermediate subnucleus
7VM: Facial nucleus ventromedial subnucleus
A1: A1 noradrenergic cells
A11: A11 dopamine cells
A13: A13 dopamine cells
A2: A2 noradrenergic cells
A35: Perirhinal cortex
A36: Ectorhinal cortex
A5: A5 noradrenaline cells
A7: A7 noradrenaline cells
AA: Anterior amygdaloid area
AcbC: Accumbens nucleus core
AcbShl: Lateral accumbens shell
ACo: Anterior cortical amygdaloid nucleus
AD: Anterodorsal thalamic nucleus
AGl: Lateral agranular prefrontal cortex
AGm: Medial agranular prefrontal cortex
AHAA: Anterior hypothalamic area anterior part
AHAC: Anterior hypothalamic area central part
AHAP: Anterior hypothalamic area posterior part
AHiAL: Amygdalohippocampal area anterolateral part
AHiPL: Amygdalohippocampal area posterolateral part
AHiPM: Amygdalohippocampal area posteromedial part
AID: Agranular insular cortex dorsal part
AIP: Agranular insular cortex posterior part
AIV: Agranular insular cortex ventral part
AM: Anteromedial thalamic nucleus
AmbC: Ambiguus nucleus compact part
AmbL: Ambiguus nucleus loose part
Ang: Angular thalamic nucleus
AOBepl: External plexiform layer of the accessory olfactory bulb
AOBgl: Granule cell layer of the accessory olfactory bulb
AOBml: Mitral cell layer of the accessory olfactory bulb
AON: Anterior olfactory nucleus
AP: Area postrema
APir: Amygdalopiriform transition area
APTD: Anterior pretectal nucleus dorsal part
APTV: Anterior pretectal nucleus ventral part
ArcD: Arcuate nucleus dorsal part
ArcL: Arcuate nucleus lateral part
ArcLP: Arcuate hypothalamic nucleus lateroposterior part
ArcMP: Arcuate hypothalamic nucleus medial posterior part
AStr: Amygdalostriatal transition area
ATg: Anterior tegmental nucleus
Au1: Primary auditory cortex
AuD: Secondary auditory cortex dorsal area
AuV: Secondary auditory cortex ventral area
AV: Anteroventral thalamic nucleus
AVPe: Anteroventral periventricular nucleus [Anterior hypothalamic area]
B: Basal nucleus Meynert
B9: B9 serotonin cells
BAC: Bed nucleus of the anterior commissure
BAOT: Bed nucleus of the accessory olfactory tract
Bar: Barringtons nucleus
BIC: Nucleus of the brachium of the inferior colliculus
BLA: Anterior basolateral nucleus
BLP: Posterior basolateral nucleus
BLV: Ventral basolateral nucleus
BMA: Anterior basomedial nucleus
BMP: Posterior basomedial nucleus
Bo: Boetzinger complex
BSTd: Bed nucleus of the stria terminalis dorsal nucleus
BSTIA: Bed nucleus of the stria terminalis intraamygdaloid division
BSTLD: Bed nucleus of the stria terminalis lateral division dorsal part
BSTLI: Bed nucleus of the stria terminalis lateral division intermedi-ate part
BSTLJ: Bed nucleus of the stria terminalis lateral division juxtacapsu-lar part
BSTLP: Bed nucleus of the stria terminalis lateral division posterior part
BSTLV: Bed nucleus of the stria terminalis lateral division ventral part
BSTMA: Bed nucleus of the stria terminalis medial division anterior part
BSTMAl: Bed nucleus of the stria terminalis anterior medial part lateral subpart
BSTMAm: Bed nucleus of the stria terminalis anterior medial part medial subpart
BSTMP: Bed nucleus of the stria terminalis medial division posterior part
BSTMPI: Bed nucleus of the stria terminalis medial division postero-intermediate part
BSTMPL: Bed nucleus of the stria terminalis medial division posterol-ateral part
BSTMPM: Bed nucleus of the stria terminalis medial division postero-medial part
BSTMV: Bed nucleus of the stria terminalis medial division ventral part
BSTSl: Supracapsular bed nucleus of the stria terminalis lateral part
BSTSm: Supracapsular bed nucleus of the stria terminalis medial part
C1: C1 adrenaline cells
C2: C2 adrenaline cells
C3: C3 adrenaline cells
CA1: Field CA1 of hippocampus
CA2: Field CA2 of hippocampus
CA3: Field CA3 of hippocampus
CeC: Capsular part
CeL: Central amygdaloid nucleus lateral division
CeM: Central amygdaloid nucleus medial division
CEnt: Caudomedial entorhinal cortex
CERC: Cerebellar cortex
Cg1: Cingulate cortex area 1
Cg2: Cingulate cortex area 2
CGA: Central gray alpha part
CGP: Central gray pons part
CI: Caudal interstitial nucleus of the medial longitudinal fascicu-lus
CIC: Central nucleus of the inferior colliculus
CL: Centrolateral thalamic nucleus
CLi: Caudal linear nucleus of the raphe
CM: Central medial thalamic nucleus
CMAM: Mammillary body
CnFD: Cuneiforme nucleus dorsal part
CnFV: Cuneiforme nucleus ventral part
COAp: Posterior amygdaloid nucleus
Com: Commissural nucleus of the inferior colliculus
CPu: Caudate putamen
CPud: Dorsal striatum
CPuim: Intermediate caudate putamen
CPul: Lateral striatum
Cu: Cuneate nucleus
CVL: Caudoventral reticular nucleus
CVLMl: Caudal ventrolateral medulla lateral part
CxA1: Cortex amygdala transition zone layer 1
DA: Dorsal hypothalamic area
DCDp: Dorsal cochlear nucleus deep core
DCeN: Cerebellar nuclei
DCFu: Dorsal cochlear nucleus fusiform layer
DCIC: Dorsal cortex of the inferior colliculus
DCl: Dorsal part of claustrum
DCNsL: Dorsal cochlear nucleus superficial layer
DI: Dysgranular insular cortex
DIEnt: Dorsal intermediate entorhinal cortex
Dk: Nucleus of Darkschewitsch
DLEnt: Dorsolateral entorhinal cortex
DLG: Dorsal geniculate nucleus
DLL: Dorsal nucleus of the lateral lemniscus
DLO: Dorsolateral orbital cortex
DLPAG: Dorsolateral periaqueductal gray
DMC: Dorsomedial hypothalamic nucleus compact part
DMD: Dorsomedial hypothalamic nucleus dorsal part
DMPAG: Dorsomedial periaqueductal gray
DMTg: Dorsomedial tegmental area
DMV: Dorsomedial hypothalamic nucleus ventral part
DP: Dorsal peduncular cortex
DpG: Deep gray layer of the superior colliculus
DPGi: Dorsal paragigantocellular nucleus
DPO: Dorsal periolivary region
DPPn: Dorsal peduncular pontine nucleus
DpWh: Deep white layer of the superior colliculus
DRC: Dorsal raphe nucleus caudal part
DRD: Dorsal raphe nucleus dorsal part
DRI: Dorsal raphe nucleus interfascicular part
DRlw: Dorsal raphe nucleus lateral wing
DRV: Dorsal raphe nucleus ventral part
DTgC: Dorsal tegmental nucleus central part
DTgP: Dorsal tegmental nucleus pericentral part
DTM: Dorsal tuberomammillary nucleus
DTr: Dorsal transition zone
DTT: Dorsal tenia tecta
ECICL1: External cortex of the inferior colliculus layer 1
ECICL2: External cortex of the inferior colliculus layer 2
ECICL3: External cortex of the inferior colliculus layer 3
ECu: External cuneate nucleus
EpP: Epipeduncular nucleus
ESO: Episupraoptic nucleus
Eth: Ethmoid thalamic nucleus
EVe: Nucleus of origin of efferents of the vestibular nerve
EW: Edinger Westphal nucleus
F: Nucleus of the fields of Forel
FC: Fasciola cinereum
Fl: Flocculus
Fr3: Frontal cortex area 3
Fu: Bed nucleus of the stria terminalis fusiform part
FVe: F cell group of the vestibular complex
GeL: Substantia gelatinosa of the trigeminal sensory nuclear com-plex
Gem: Gemini hypothalamic nucleus
GI: Granular insular cortex
GiA: Gigantocellular reticular nucleus alpha part
GiV: Gigantocellular reticular nucleus ventral part
GlA: Glomerular layer accessory olfactory bulb
Gr: Gracile nucleus principal part
GrDG: Granular layer of the dentate gyrus
HDB: Nucleus of the horizontal limb of the diagonal band
I: Intercalated nuclei of the amygdala
I8: Interstitial nucleus of the vestibulocochlear nerve
IAM: Interoanteromedial thalamic nucleus
IcaM: Intercalated amygdaloid nucleus main part
ICjM: Islands of Calleja major island
ID: Interstitial nucleus of the decussation of the superior cerebellar peduncle
IF: Interfascicular nucleus
IG: Indusium griseum
IGL: Intergeniculate leaf
II: Intermediate interstitial nucleus of the medial longitudinal fasciculus
IL: Infralimbic cortex
ILL: Intermediate nucleus of the lateral lemniscus
IMA: Intramedullary thalamic area
IMD: Intermediodorsal thalamic nucleus
IMG: Amygdaloid intramedullary gray
iml: Internal medullary lamina
In: Intercalated nucleus of the medulla
InC: Interstitial nucleus of Cajal
InG: Intermediate gray layer of the superior colliculus
InWh: Intermediate white layer of the superior colliculus
IOA: Inferior olive subnucleus A of medial nucleus
IOB: Inferior olive subnucleus B of medial nucleus
IOBe: Inferior olive beta subnucleus
IOC: Inferior olive subnucleus C of medial nucleus
IOD: Inferior olive dorsal nucleus
IODM: Inferior olive dorsomedial cell group
IOK: Inferior olive cap of Kooy of the medial nucleus
IOVL: Inferior olive ventrolateral protrusion
IPA: Interpeduncular nucleus apical subnucleus
IPACL: Interstitial nucleus of the posterior limb of the anterior com-missure lateral part
IPACM: Interstitial nucleus of the posterior limb of the anterior com-missure medial part
IPC: Interpeduncular nucleus caudal subnucleus
IPDL: Interpeduncular nucleus dorsolateral subnucleus
IPDM: Interpeduncular nucleus dorsomedial subnucleus
IPI: Interpeduncular nucleus intermediate subnucleus
IPL: Interpeduncular nucleus lateral subnucleus
IPR: Interpeduncular nucleus rostral subnucleus
IRtA: Intermediate reticular nucleus alpha part
IS: Inferior salivatory nucleus
isRT: Isthmic reticular formation
JPV: Juxtaparaventricular part [Lateral hypothalamic area [Preoptic anterior region]]
JxO: Juxtaolivary nucleus
KF: Koelliker Fuse nucleus
LaDL: Dorsolateral part of the lateral nucleus
LAH: Lateroanterior hypothalamic nucleus
LaVL: Ventrolateral part of the lateral nucleus
LaVM: Ventromedial part of the lateral nucleus
LC: Locus coeruleus
Ld: Lambdoid septal zone
LDDM: Laterodorsal thalamic nucleus dorsomedial part
LDTgV: Laterodorsal tegmental nucleus ventral part
LDVL: Laterodorsal thalamic nucleus ventrolateral part
LEnt: Lateral entorhinal cortex
LGP: Lateral globus pallidus
LH: Lateral hypothalamic area
LHbL: Lateral habenular nucleus lateral part
LHbM: Lateral habenular nucleus medial part
LHTUC: Lateral hypothalamic area [Region of tuber cinereum]
Li: Linear nucleus of the medulla
LNol: Lacunosum molecular layer of the hippocampus
LO: Lateral orbital cortex
LOTL1: Nucleus of the lateral olfactory tract layer 1
LOTL2: Nucleus of the lateral olfactory tract layer 2
LOTL3: Nucleus of the lateral olfactory tract layer 3
LPAG: Lateral periaqueductal gray
LPB: Lateral parabrachial nucleus
LPBGiA: Lateral paragigantocellular nucleus alpha part
LPBGiE: Lateral paragigantocellular nucleus external part
LPLC: Lateral posterior thalamic nucleus laterocaudal part
LPLR: Lateral posterior thalamic nucleus laterorostral part
LPMC: Lateral posterior thalamic nucleus mediocaudal part
LPMR: Lateral posterior thalamic nucleus mediorostral part
LPO: Lateral preoptic area
LRtPC: Lateral reticular nucleus parvicellular part
LRtS5: Lateral reticular nucleus subtrigeminal part
LSD: Lateral septal nucleus dorsal part
LSI: Lateral septal nucleus intermediate part
LSO: Lateral superior olive
LSS: Lateral stripe of the striatum
LSV: Lateral septal nucleus ventral part
LTeN: Lateral terminal nucleus of the accessory optic tract
LTer: Lamina terminalis
LVe: Lateral vestibular nucleus
LVPO: Lateroventral periolivary nucleus
MA3: Medial accessory oculomotor nucleus
MCLH: Magnocellular nucleus of the lateral hypothalamus
MCPC: Magnocellular nucleus of the posterior commissure
MCPO: Magnocellular preoptic nucleus
MDC: Mediodorsal thalamic nucleus central part
MDL: Mediodorsal thalamic nucleus lateral part
MDM: Mediodorsal thalamic nucleus medial part
MdVL: Medullary reticular nucleus ventrolateral part
ME: Median eminence
Me5: Mesencephalic trigeminal nucleus
MeAD: Medial amygdaloid nucleus anterodorsal part
MeAV: Medial amygdaloid nucleus anteroventral part
MEI: Medial eminence internal layer
MEntR: Medial entorhinal cortex rostral part
MePD: Medial amygdaloid nucleus posterodorsal part
MePV: Medial amygdaloid nucleus posteroventral part
MGD: Medial geniculate nucleus dorsal part
MGM: Medial geniculate nucleus medial part
MGP: Medial globus pallidus
MGV: Medial geniculate nucleus ventral part
MHb: Medial habenular nucleus
MiTg: Microcellular tegmental nucleus
MnPO: Median preoptic nucleus
MnR: Median raphe nucleus
MO: Medial orbital cortex
Mo5: Motor trigeminal nucleus
Mol: Molecular layer of the dentate gyrus
MPA: Medial preoptic area
MPL: Medial paralemniscal nucleus
MPO: Medial preoptic nucleus
MPOC: Medial preoptic nucleus central part
MPOL: Medial preoptic nucleus lateral part
MPOM: Medial preoptic nucleus medial part
MPT: Medial pretectal nucleus
MS: Medial septal nucleus
MSO: Medial superior olive
MT: Medial terminal nucleus of the accessory optic tract
MTuN: Medial tuberal nucleus
MVe: Medial vestibular nucleus
MVPO: Medioventral periolivary nucleus
Mx: Matrix region of the medulla
MZMG: Marginal zone of the medial geniculate
NC: Nucleus circularis
NCAT: Nucleus of the central acoustic tract
Neck: Nucleus of the spinal accessory nerve
Nv: Navicular nucleus of the basal forebrain
O: Nucleus O
OcxL: Olfactory cortex layers
Op: Optic nerve layer of the superior colliculus
OPC: Oval paracentral thalamic nucleus
OPT: Olivary pretectal nucleus
OT: Nucleus of the optic tract
OV: Olfactory ventricle
p1PAG: P1 periaqueductal gray
p1RT: Reticular thalamic nucleus prosomere 1
P5: Peritrigeminal zone
P7: Perifacial zone
Pa4: Paratrochlear nucleus
Pa5: Paratrigeminal nucleus
Pa6: Paraabducens nucleus
PaAMp: Paraventricular nucleus of the hypothalamus magnocellular division posterior magnocellular part
PaAP: Paraventricular hypothalamic nucleus anterior parvicellular part [Periventricular zone anterior region]
PaDC: Paraventricular hypothalamic nucleus dorsal cap [Periventricu-lar zone anterior region]
PaLM: Paraventricular hypothalamic nucleus lateral magnocellular part [Periventricular zone anterior region]
PaMP: Paraventricular hypothalamic nucleus medial parvicellular part [Periventricular zone anterior region]
PaPo: Paraventricular hypothalamic nucleus posterior part [Periventricular zone anterior region]
PaR: Pararubral nucleus
PaS: Parasubiculum
PaV: Paraventricular hypothalamic nucleus ventral part [Periventric-ular zone anterior region]
PBG: Parabigeminal nucleus
PBnMe: Parabrachial nucleus medial
PBP: Parabrachial pigmented nucleus
PC: Paracentral thalamic nucleus
PCGS: Parachochlear glial substance
PCnFa: Precuneiform area
PCom: Nucleus of the posterior commissure
PCRtA: Parvicellular reticular nucleus alpha part
PDP: Posterodorsal preoptic nucleus
PDR: Posterodorsal raphe nuclei
PDTg: Posterodorsal tegmental nucleus
PeRHO: Area periventricularis hypothalamica communis [Regio hypothalamus oralis]
PF: Parafascicular thalamic nucleus
PH: Posterior hypothalamic nucleus
PHD: Posterior hypothalamic area dorsal part
PIL: Posterior intralaminar thalamic nucleus
Pir1a: Piriform cortex layer 1a
Pir1b: Piriform cortex layer 1b
PirL2: Piriform cortex layer 2
PirL3: Piriform cortex layer 3
PiSt: Pineal stalk
PLCo: Posterolateral cortical nucleus
PLi: Posterior limitans thalamic nucleus
PMCo: Posteromedial cortical nucleus
PMn: Paramedian reticular nucleus
PMnR: Paramedian raphe nucleus
PN: Paranigral nucleus
Pn: Pontine nuclei
PNC: Pontine reticular nucleus caudal part
PnO: Pontine reticular nucleus oral part
PnR: Pontine raphe nucleus
PnV: Pontine reticular nucleus ventral part
Po: Posterior thalamic nuclear group
POAd7fn: Periolivary area between superior olivary nucleus and descend-ing root of the facial nerve
PoMn: Posteromedian thalamic nucleus
POR: Postrhinal cortex
Post: Postsubiculum
PP: Peripeduncular nucleus
PPT: Posterior pretectal nucleus
PPTg: Pedunculopontine tegmental nucleus
PPy: Parapyramidal nucleus
Pr: Prepositus nucleus
PR: Prerubral field
Pr5: Principal sensory trigeminal nucleus
PrBo: Pre Boetzinger complex
PrC: Precommissural nucleus
PrL: Prelimbic cortex
PrS: Presubiculum
PS: Parastriatal nucleus
PSol: Parasolitary nucleus
PT: Paratenial thalamic nucleus
PtA: Parietal association cortex
PV: Paraventricular thalamic nucleus
PVA: Paraventricular thalamic nucleus anterior part
PVP: Paraventricular thalamic nucleus posterior part
RAmb: Retroambiguus nucleus
Rbd: Rhabdoid nucleus
RChL: Retrochiasmatic area lateral part
Re: Reuniens thalamic nucleus
REth: Retroethmoid nucleus
Rh: Rhomboid thalamic nucleus
RI: Rostral interstitial nucleus of medial longitudinal fasciculus
RIP: Raphe interpositus nucleus
RL: Retrolemniscal nucleus
RLi: Rostral linear nucleus of the raphe
RMC: Red nucleus magnocellular part
RMg: Raphe magnus nucleus
Ro: Nucleus ofoller
ROb: Raphe obscurus nucleus
ROC: Red nucleus parvicellular part
RPa: Raphe pallidus nucleus
RPF: Retroparafascicular nucleus
RR: Retrorubral nucleus
RRE: Retroreuniens area
RRF: A8 dopamine cells retrorubral group
RSd: Retrosplenial dorsal
RSGaL: Retrosplenial granular a cortex layers
RSGbL: Retrosplenial granular b cortex layers
RSGC: Retrosplenial granular cortex caudal part
RSGcC: Retrosplenial granular cortex c region caudal part
RtTgL: Reticulotegmental nucleus of the pons lateral part
RtTgP: Reticulotegmental nucleus of the pons pericentral part
RVL: Rostroventrolateral reticular nucleus
RVRG: Rostral ventral respiratory group
S: Subiculum
S1: Primary somatosensory cortex
S2: Secondary somatosensory cortex
S5: Sensory root of the trigeminal nerve
Sag: Sagulum nucleus
SChDL: Suprachiasmatic nucleus dorsolateral part
SChVM: Suprachiasmatic nucleus ventromedial part
SCO: Subcommissural organ
SCzo: Superior colliculus zonal layer
SFi: Septofimbrial nucleus
SFO: Subfornical organ
SG: Suprageniculate thalamic nucleus
SGe: Supragenual nucleus
SHi: Septohippocampal nucleus
SHy: Septohypothalamic nucleus
SIB: Substantia innominata basal part
SLEAc: Central division of sublenticular extended amygdala
SLEAm: Medial division of the sublenticular extended amygdala
SM: Nucleus of the stria medullaris
SNC: Substantia nigra compact part
SNL: Substantia nigra lateral part
SNR: Substantia nigra reticular part
SolC: Nucleus of the solitary tract commissural part
SolCe: Nucleus of the solitary tract central part
SolDL: Nucleus of the solitary tract dorsolateral part
SolDM: Nucleus of the solitary tract dorsomedial part
SolG: Nucleus of the solitary tract gelatinous part
Soli: Nucleus of the solitary tract interstitial part
SolIM: Nucleus of the solitary tract intermediate part
SolL: Nucleus of the solitary tract lateral part
SolM: Nucleus of the solitary tract medial part
SolRL: Nucleus of the solitary tract rostrolateral part
SolV: Nucleus of the solitary tract ventral part
SolVL: Nucleus of the solitary tract ventrolateral part
SOR: Supraoptic nucleus retrochiasmatic part
Sp5nc: Spinal trigeminal nucleus
SPa: Subparaventricular zone of the hypothalamus
SPFPC: Submedius thalamic nucleus parvicellular part
SPFr: Subparafascicular thalamic nucleus rostral part
Sph: Sphenoid nucleus
SPO: Superior paraolivary nucleus
SPTg: Subpeduncular tegmental nucleus
SpVe: Spinal vestibular nucleus
StA: Strial part of the preoptic area
STh: Subthalamic nucleus
StHy: Striohypothalamic nucleus
Su3: Supraoculomotor periaqueductal gray
Su3C: Supraoculomotor cap
Su5: Supratrigeminal nucleus
SubB: Subbrachial nucleus
SubCA: Subcoeruleus nucleus alpha part
SubCD: Subcoeruleus nucleus dorsal part
SubCV: Subcoeruleus nucleus ventral part
SubD: Submedius thalamic nucleus dorsal part
SubI: Subincertal nucleus
SubP: Subpostrema area
SubV: Submedius thalamic nucleus ventral part
SuG: Superficial gray layer of the superior colliculus
SuS: Superior salivatory nucleus
SuVe: Superior vestibular nucleus
TC: Tuber cinereum area
Te: Terete hypothalamic nucleus
TeA: Temporal association cortex 1
TS: Triangular septal nucleus
TuOd: Olfactory tubercle densocellular layer
TuOLa1: Olfactory tubercle plexiform layer
TuOpo: Olfactory tubercle polymorph layer
Tz: Nucleus of the trapezoid body
V1: Primary visual cortex
V1B: Primary visual cortex binocular area
V1M: Primary visual cortex monocular area
V2L: Secondary visual cortex lateral area
V2ML: Secondary visual cortex mediolateral area
V2MM: Secondary visual cortex mediomedial area
VA: Ventro anterior thalamic nucleus
VCA: Ventral cochlear nucleus anterior part
VCAGr: Ventral cochlear nucleus granule cell layer
VCl: Ventral part of claustrum
VCP: Ventral cochlear nucleus posterior part
VDB: Nucleus of the vertical limb of the diagonal band
VeCb: Vestibulocerebellar nucleus
VEn: Ventral endopiriform nucleus
VG: Ventral geniculate nucleus
VIEnt: Ventral intermediate entorhinal cortex
VL: Ventrolateral thalamic nucleus
VLG: Ventral lateral geniculate nucleus
VLH: Ventrolateral hypothalamic nucleus
VLL: Ventral nucleus of the lateral lemniscus
VLPAG: Ventrolateral periaqueductal gray
VLPO: Ventrolateral preoptic nucleus
VM: Ventromedial thalamic nucleus
VMHC: Ventromedial hypothalamic nucleus central part
VMHDM: Ventromedial hypothalamic nucleus dorsomedial part
VMHVL: Ventromedial hypothalamic nucleus ventrolateral part
VMPO: Ventromedial preoptic nucleus
VO: Ventral orbital cortex
VOLT: Vascular organ of the lamina terminalis
VP: Ventral pallidum
VPL: Ventral posterolateral thalamic nucleus
VPM: Ventral posteromedial thalamic nucleus
VPPC: Ventral posterior thalamic nucleus parvicellular part
VRe: Ventral reuniens thalamic nucleus
VTAR: Ventral tegmental area rostral part
VTg: Ventral tegmental nucleus
VTM: Ventral tuberomammillary nucleus
VTT: Ventral tenia tecta
X: Nucleus X
Xi: Xiphoid thalamic nucleus
Y: Nucleus Y
Z: Nucleus Z
ZIC: Zona incerta caudal part
ZID: Zona incerta dorsal part
ZIR: Zona incerta rostral part
ZIV: Zona incerta ventral part
ZL: Zona limitans

### S2 Tables

**Table 1.**
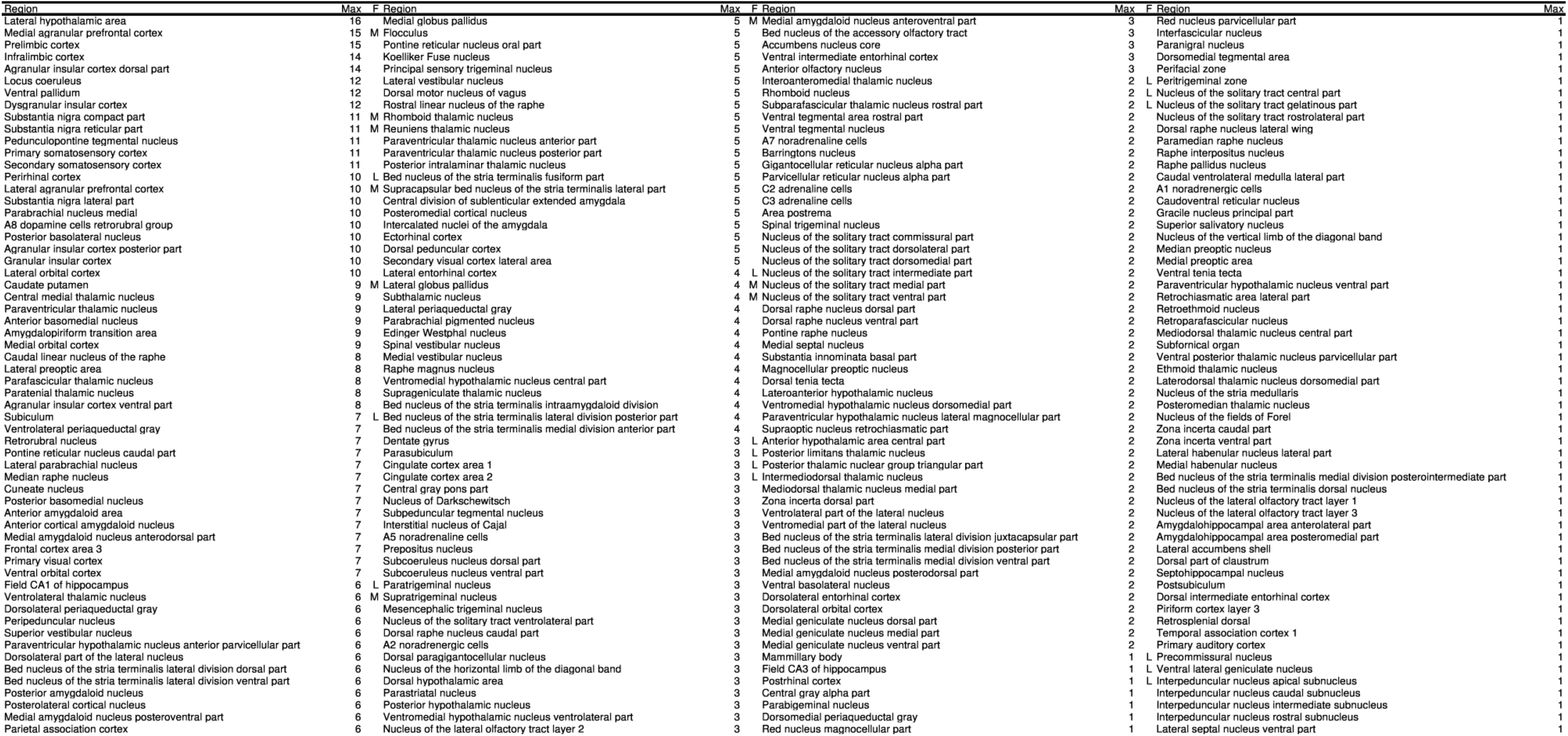
Sorted sum of connections of non-lesioned and lesioned ICH regions. The column “Region” contains non-lesioned regions and the column “Max” the number of connections of non-lesioned regions with lesioned regions. The regions with functional markers (F) are motor (M) or learning regions (L). E.g., the medial agranular prefrontal cortex is a motor region (M) and has 15 connections with ICH lesioned regions. In the last column of non-lesioned regions which have only 1 connection with ICH lesioned region no functional assignments are available. Anterodorsal thalamic nucleus rostral part (L), anteroventral thalamic nucleus ventral part (L), field CA2 of hippocampus (L), presubiculum (L), cerebellar cortex (M), cerebellar nuclei (M) and pontine nuclei (M) are non-lesioned regions with functional assignments, however, without connections from or to ICH lesioned regions.

**Table 2.** Connections of dMCAO lesioned regions to motor and learning regions. The relation of functional groups (Marker: motor behavior (2), learning behavior (3)) to 6 lesioned regions in the dMCAO model. Regions were sorted by their sum of connections and reciprocal connections. E.g. the lateral enthorinal cortex has 6 connections to all lesioned regions and all of these 6 connections are reciprocal.

| Region | Sum | PtA | AID | AIV | DI | GI | S1 | Reciprocal | Marker |
| --- | --- | --- | --- | --- | --- | --- | --- | --- | --- |
| Lateral entorhinal cortex | 6 | 1 | 1 | 1 | 1 | 1 | 1 | 6 | 3 |
| Perirhinal cortex | 6 | 1 | 1 | 1 | 1 | 1 | 1 | 6 | 3 |
| Postrhinal cortex | 6 | 1 | 1 | 1 | 1 | 1 | 1 | 5 | 3 |
| Reuniens thalamic nucleus | 6 | 1 | 1 | 1 | 1 | 1 | 1 | 4 | 3 |
| Rhomboid thalamic nucleus | 6 | 1 | 1 | 1 | 1 | 1 | 1 | 3 | 3 |
| Field CA1 of hippocampus | 5 | 1 | 1 | 1 | 1 | 1 | 0 | 0 | 3 |
| Posterior thalamic nuclear group | 4 | 1 | 0 | 0 | 1 | 1 | 1 | 3 | 3 |
| Anteroventral thalamic nucleus | 4 | 1 | 1 | 0 | 1 | 0 | 1 | 1 | 3 |
| Mammillary body | 4 | 1 | 1 | 0 | 0 | 1 | 1 | 0 | 3 |
| Cingulate cortex area 1 | 3 | 1 | 1 | 0 | 0 | 0 | 1 | 2 | 3 |
| Cingulate cortex area 2 | 3 | 1 | 1 | 0 | 0 | 0 | 1 | 2 | 3 |
| Presubiculum | 3 | 1 | 0 | 1 | 0 | 1 | 0 | 1 | 3 |
| Parasubiculum | 3 | 1 | 0 | 1 | 0 | 1 | 0 | 1 | 3 |
| Subiculum | 3 | 1 | 1 | 1 | 0 | 0 | 0 | 0 | 3 |
| Anterodorsal thalamic nucleus | 1 | 0 | 1 | 0 | 0 | 0 | 0 | 0 | 3 |
| Interoanteromedial thalamic nucleus | 1 | 0 | 0 | 0 | 1 | 0 | 0 | 0 | 3 |
| Field CA3 of hippocampus | 1 | 1 | 0 | 0 | 0 | 0 | 0 | 0 | 3 |
| Subparafascicular thalamic nucleus rostral part | 0 | 0 | 0 | 0 | 0 | 0 | 0 | 0 | 3 |
| Dentate gyrus | 0 | 0 | 0 | 0 | 0 | 0 | 0 | 0 | 3 |
| Field CA2 of hippocampus | 0 | 0 | 0 | 0 | 0 | 0 | 0 | 0 | 3 |
| Lateral agranular prefrontal cortex | 6 | 1 | 1 | 1 | 1 | 1 | 1 | 6 | 2 |
| Medial agranular prefrontal cortex | 6 | 1 | 1 | 1 | 1 | 1 | 1 | 5 | 2 |
| Subthalamic nucleus | 6 | 1 | 1 | 1 | 1 | 1 | 1 | 3 | 2 |
| Caudate putamen | 6 | 1 | 1 | 1 | 1 | 1 | 1 | 2 | 2 |
| Substantia nigra compact part | 4 | 1 | 0 | 0 | 1 | 1 | 1 | 3 | 2 |
| Pontine nuclei | 4 | 1 | 0 | 1 | 0 | 1 | 1 | 1 | 2 |
| Substantia nigra reticular part | 4 | 1 | 1 | 0 | 0 | 1 | 1 | 0 | 2 |
| Ventrolateral thalamic nucleus | 3 | 1 | 0 | 1 | 0 | 0 | 1 | 2 | 2 |
| Medial globus pallidus | 2 | 1 | 0 | 0 | 0 | 0 | 1 | 1 | 2 |
| Cerebellar cortex | 2 | 1 | 0 | 0 | 0 | 0 | 1 | 0 | 2 |
| Cerebellar nuclei | 1 | 0 | 0 | 0 | 0 | 0 | 1 | 0 | 2 |
| Lateral globus pallidus | 0 | 0 | 0 | 0 | 0 | 0 | 0 | 0 | 2 |

**Table 3.**
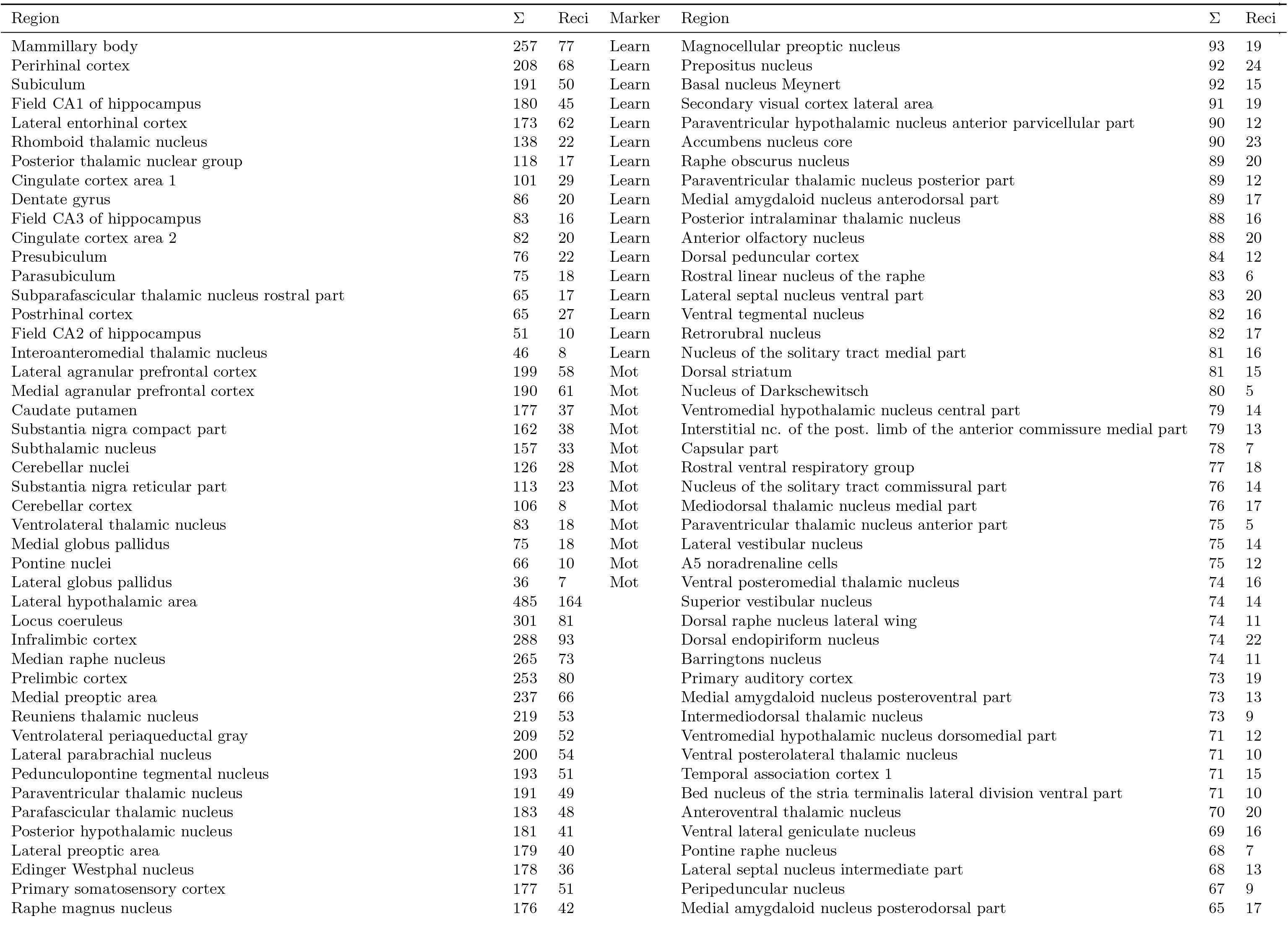
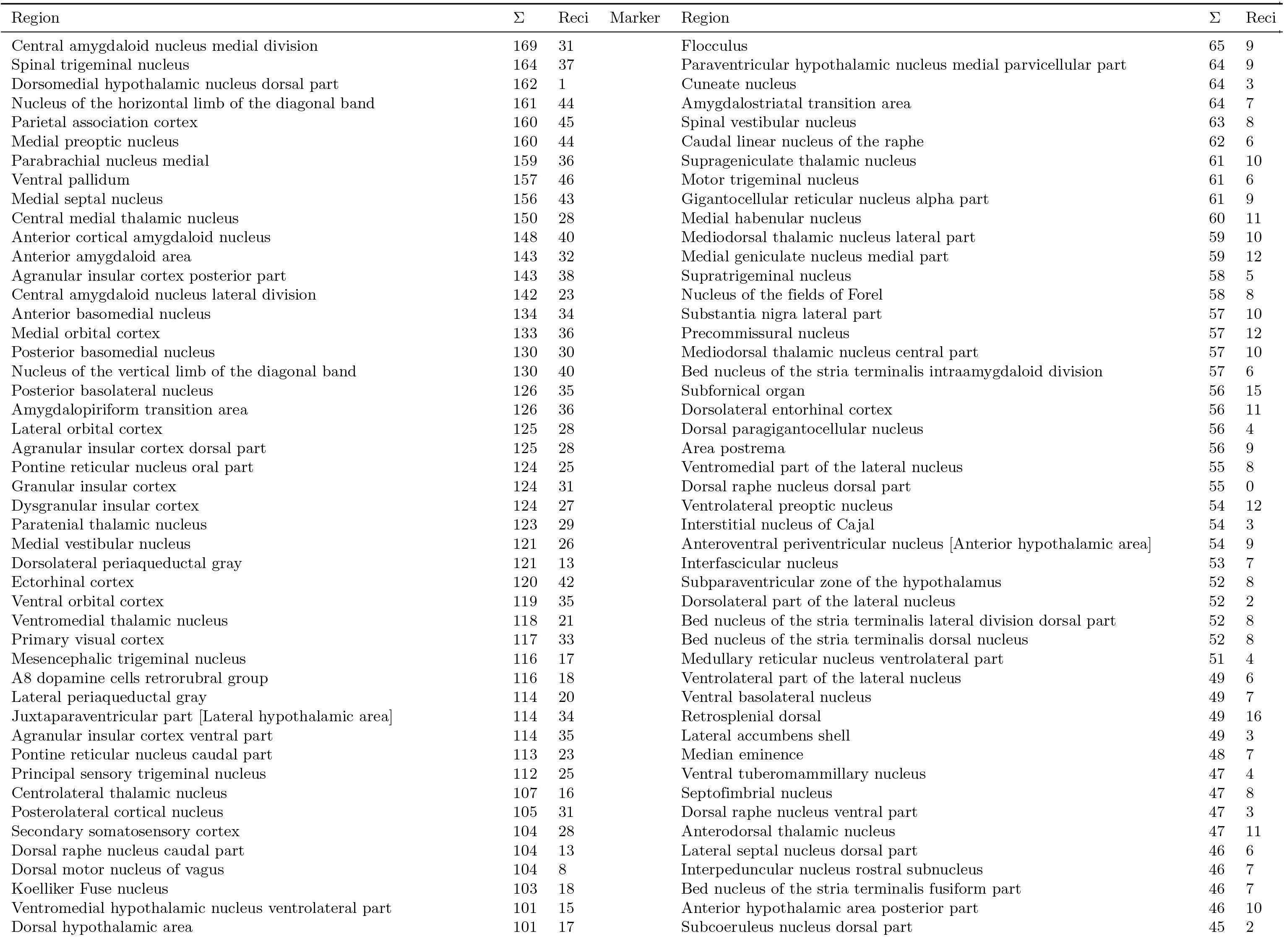
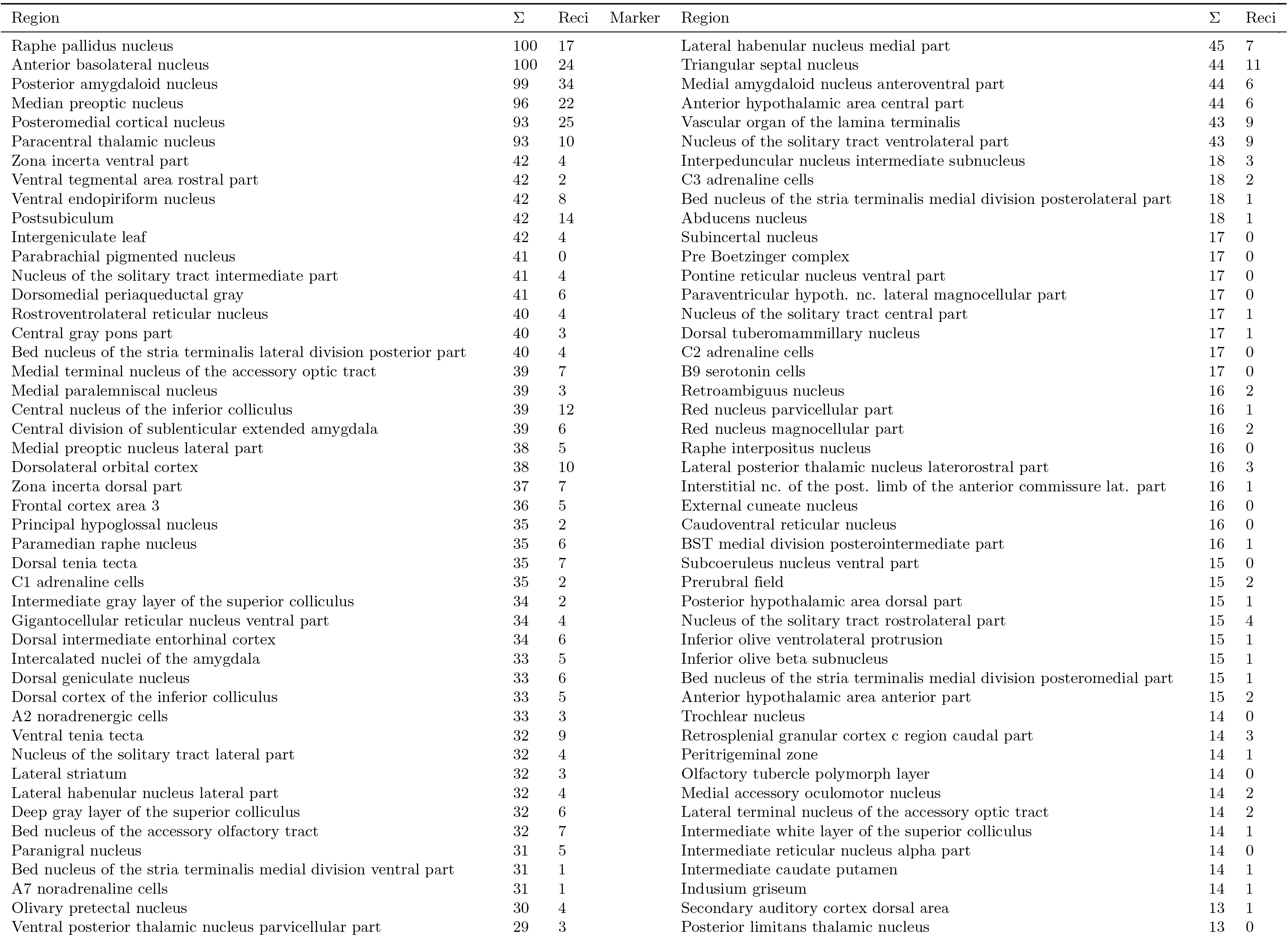
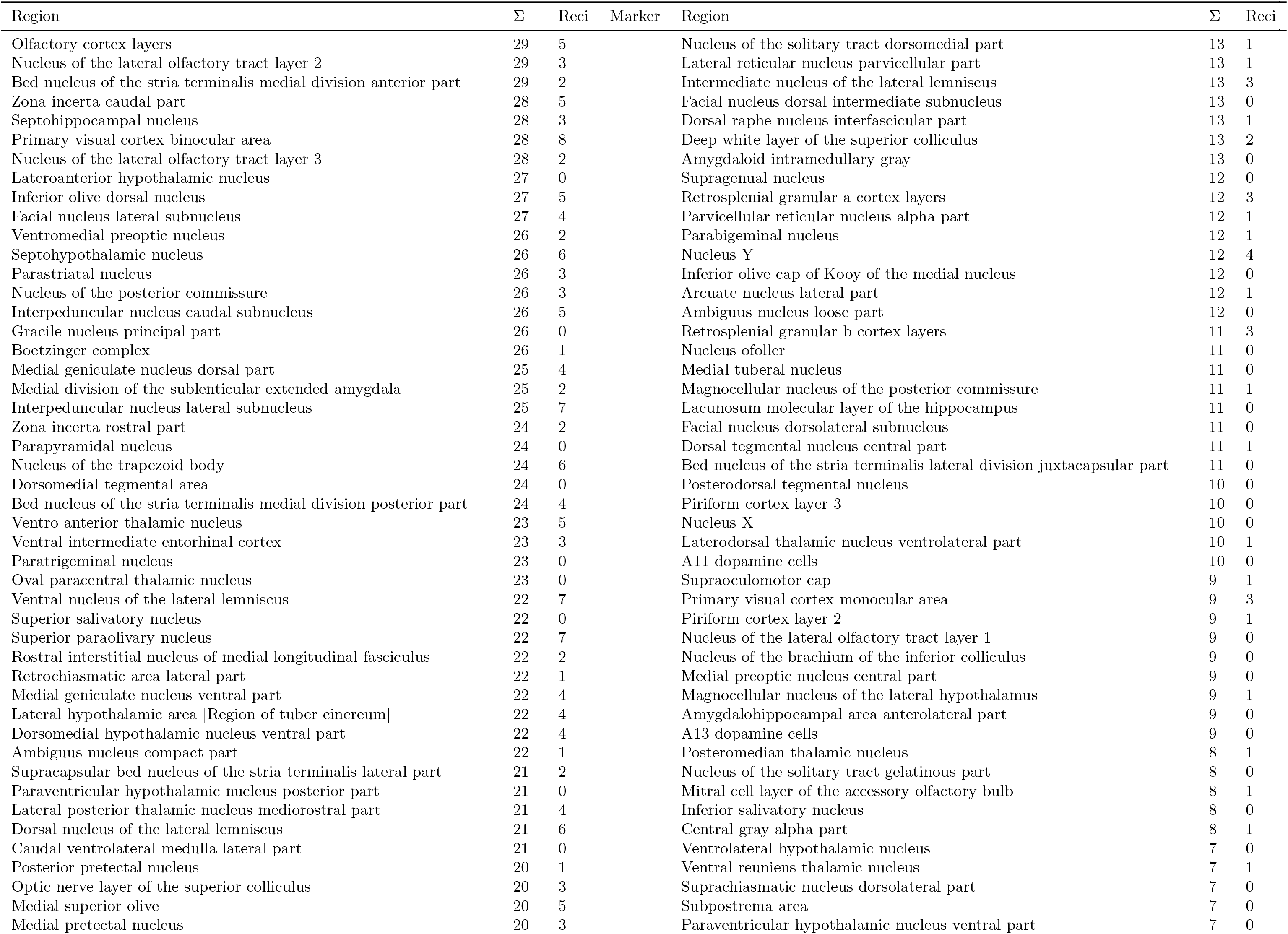
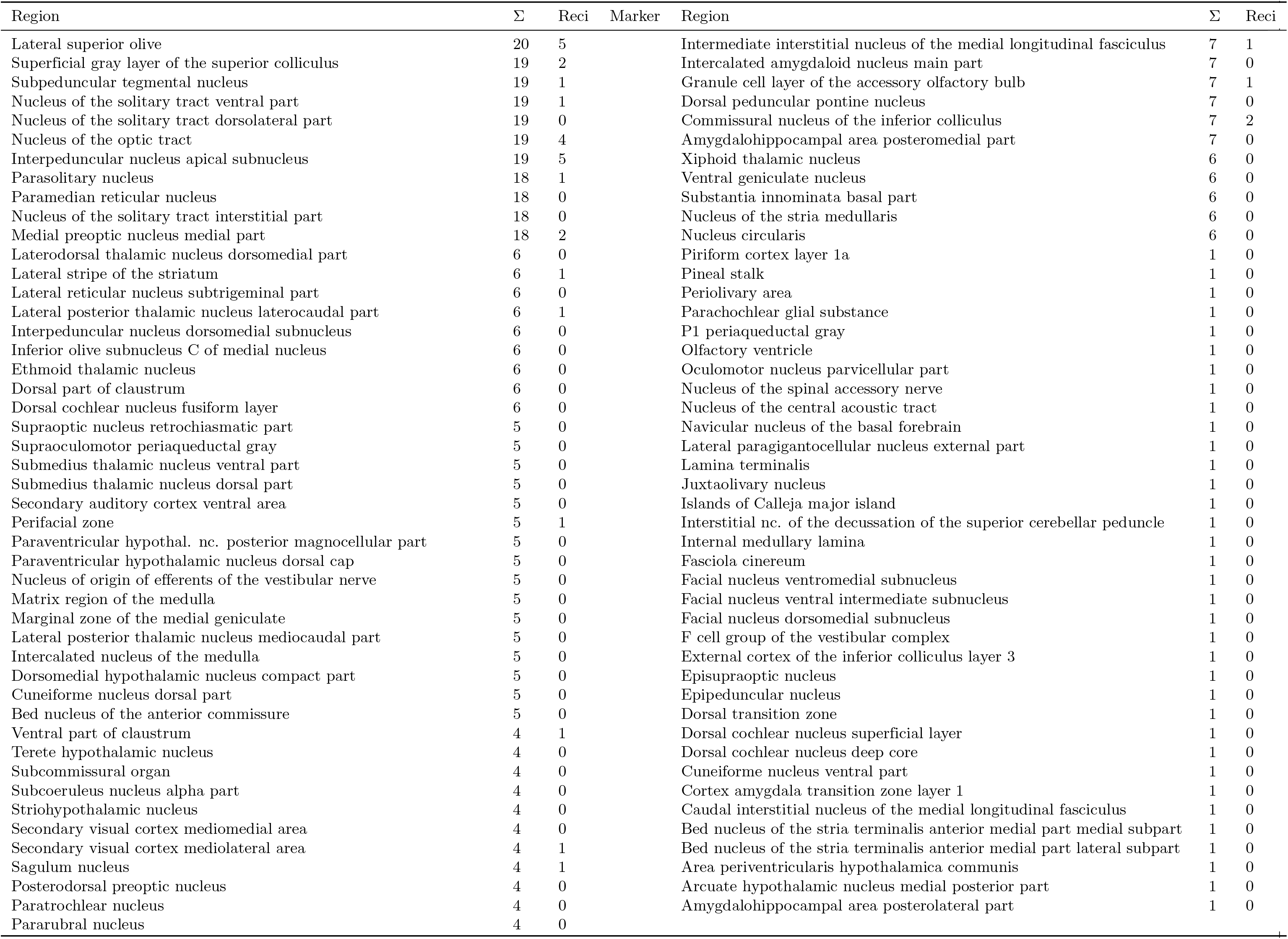
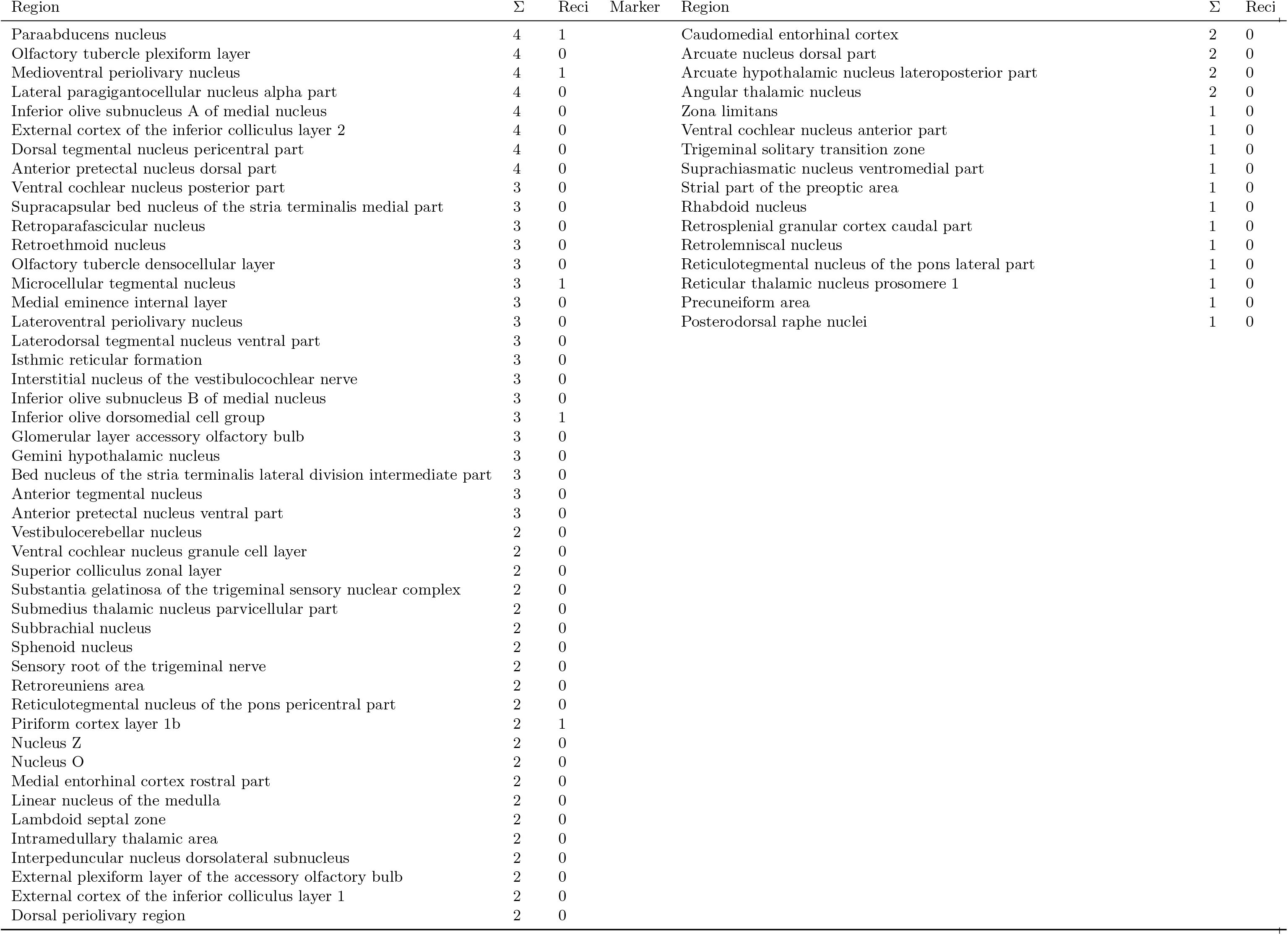
Overview of all connections of dMCAO lesioned regions. Connections of dMCAO lesioned regions, functionally defined regions and control regions without functional definitions.

**Table 4.**
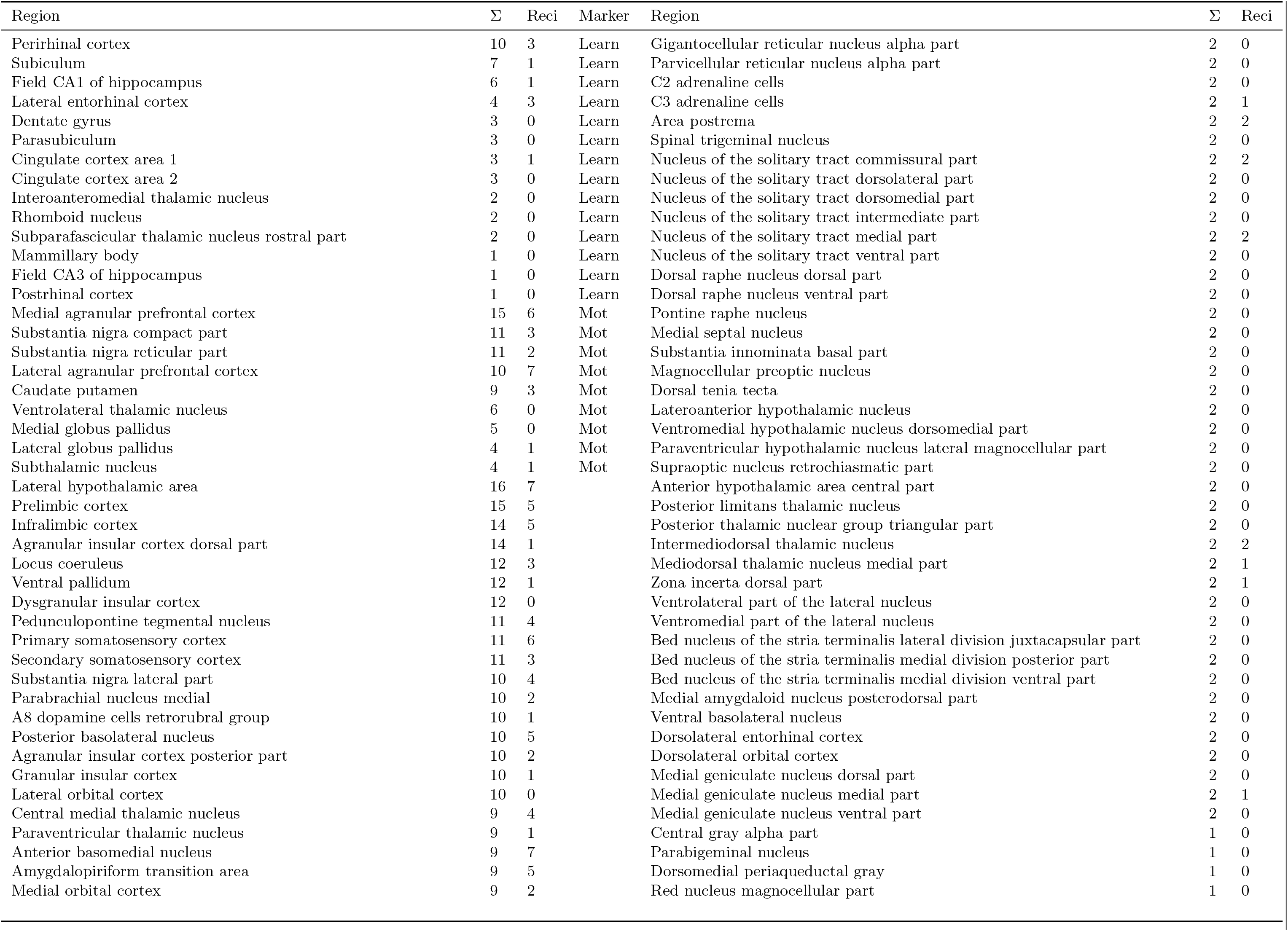
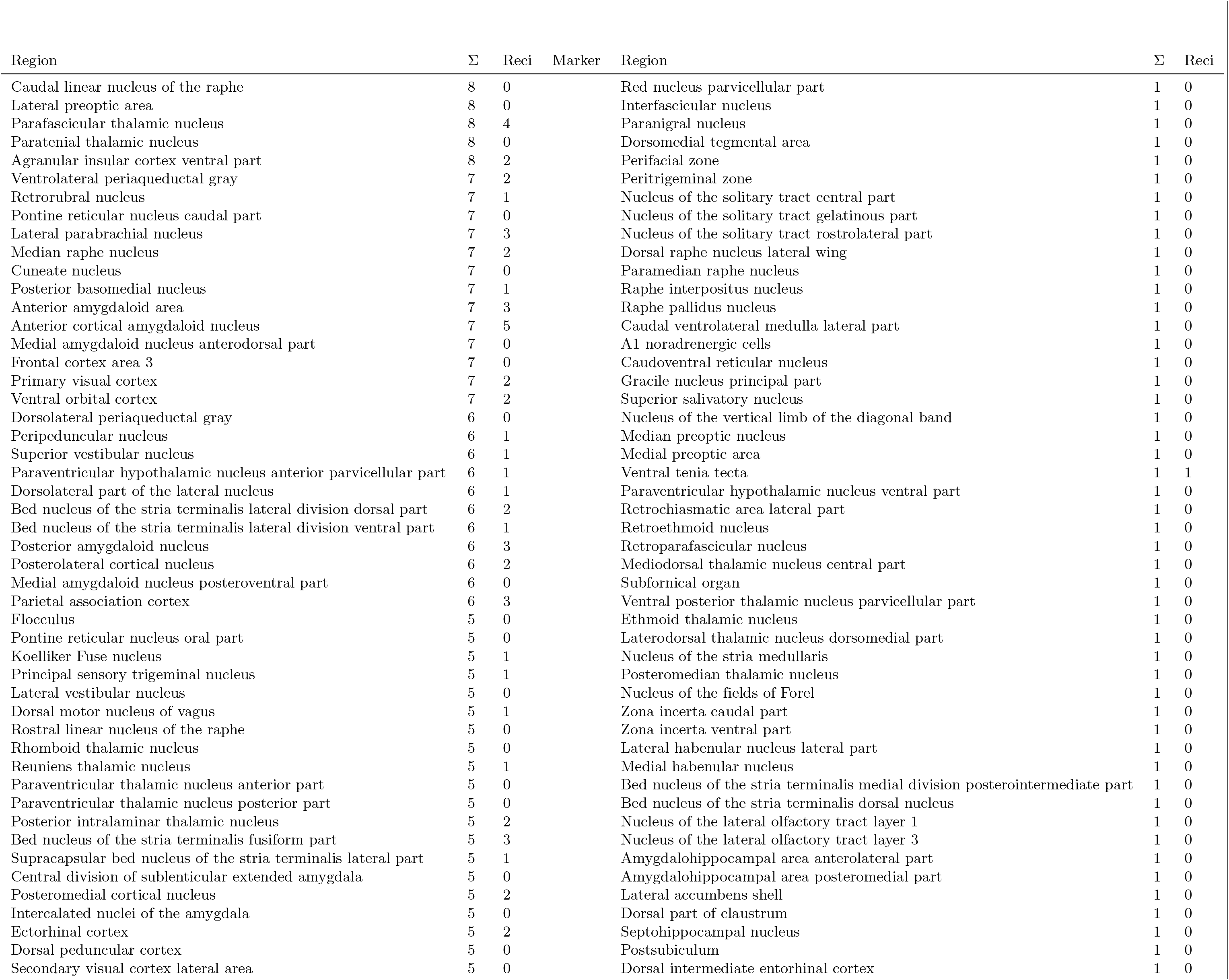
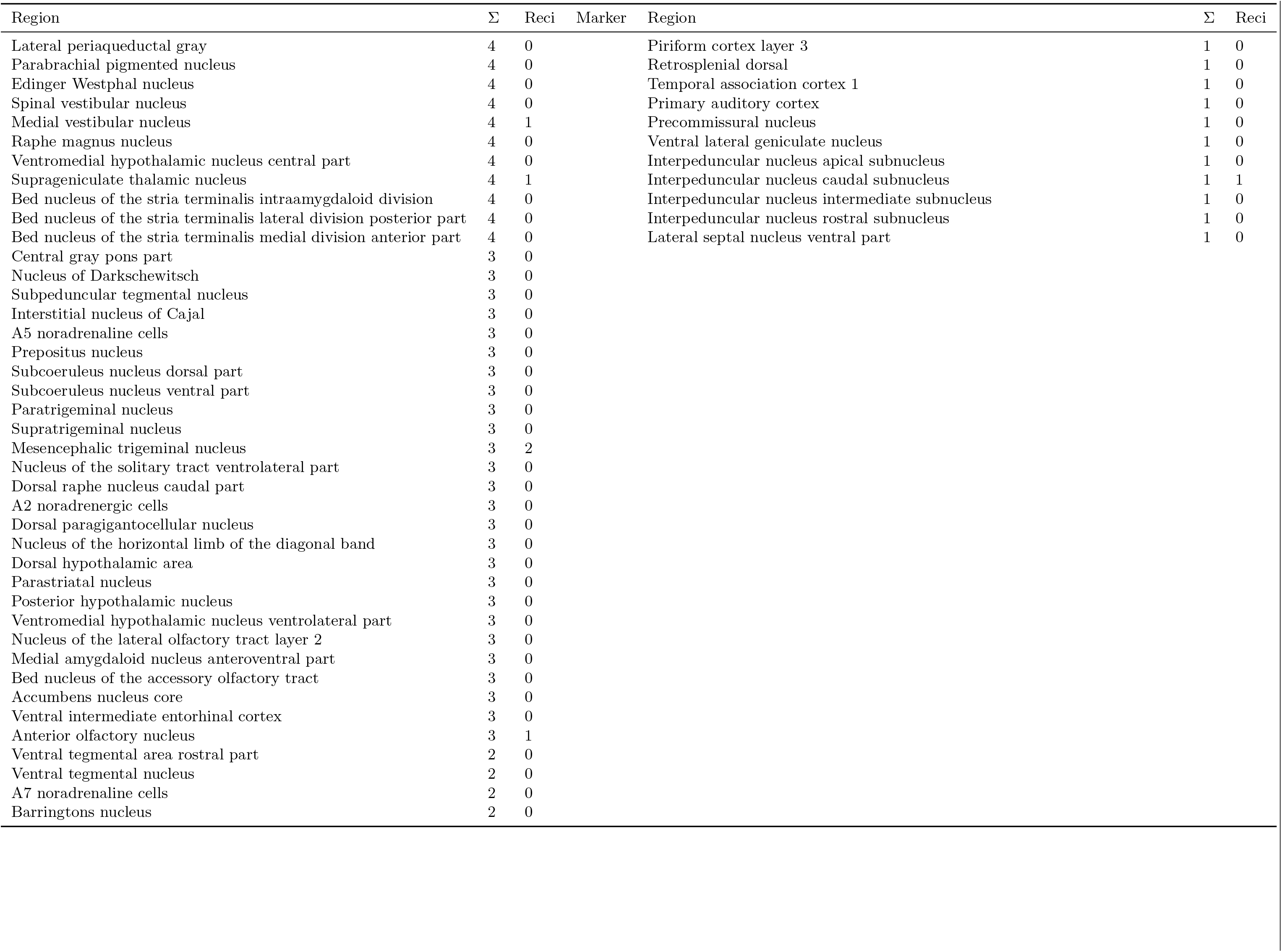
Overview of all connections of ICH lesioned regions. Connections of ICH lesioned regions, functionally defined regions and control regions without functional definitions.

**Table 5.**
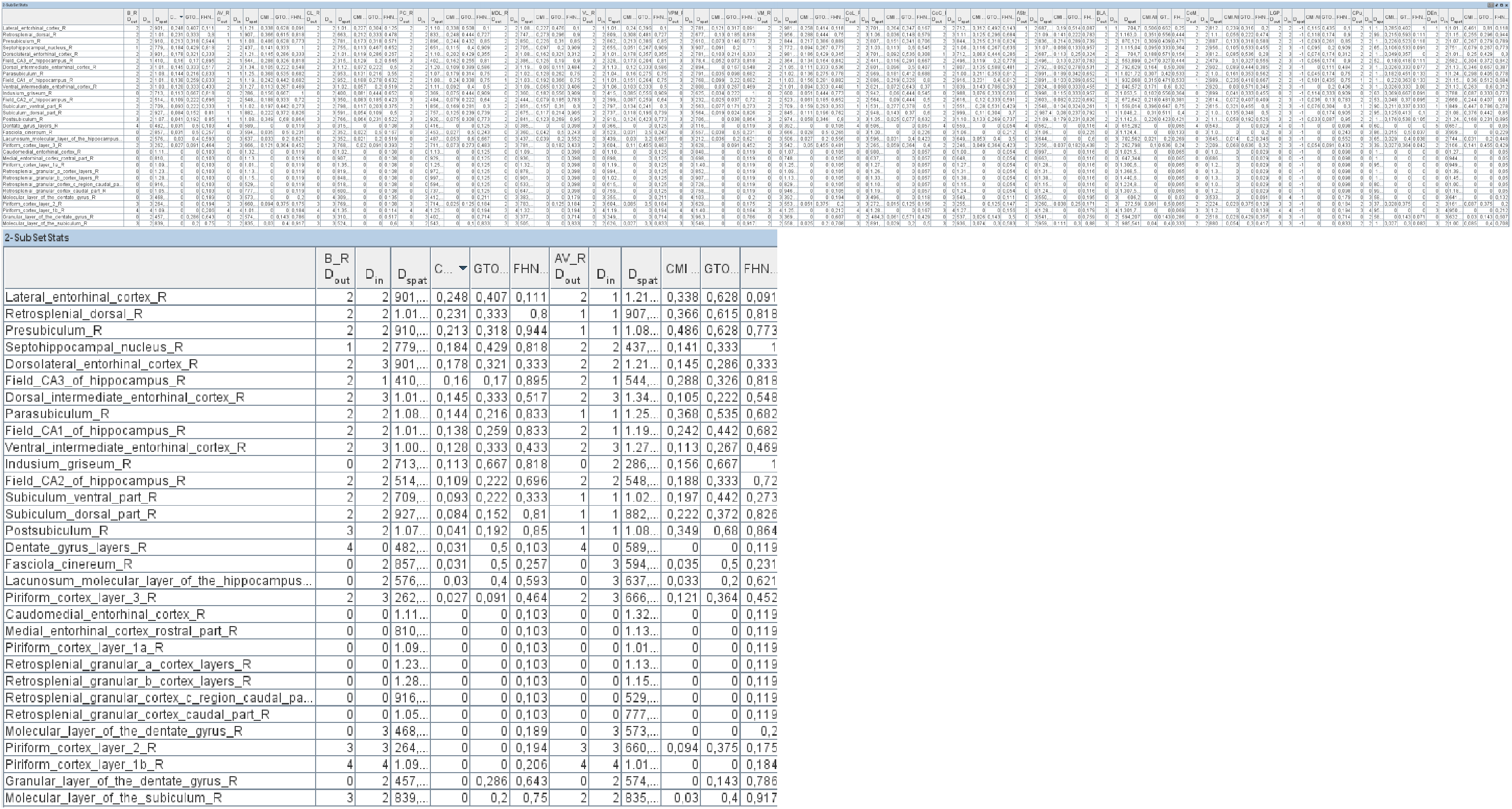
Identification of ICH lesioned regions strongly connected with functionally defined regions. This table (subset stats table) allows comparison of several parameters and matrices like the connectivity matching matrix (*CMI_All_*), GTOM and FHN of the lesioned regions (2nd, 8th, 16th . . . column) of the ICH connectome with non lesioned regions (first colum, all rows). The lower table displays a clipping of the lesioned regions basal nucleus Meynert (B R) and anteroventral thalamic nucleus (AV R). The non lesioned regions were sorted by the *CMI_All_* parameter: The lateral enthorinal cortex has the largest *CMI_All_* value (from all non lesioned regions) with the basal nucleus Meynert.

**Table 6.**
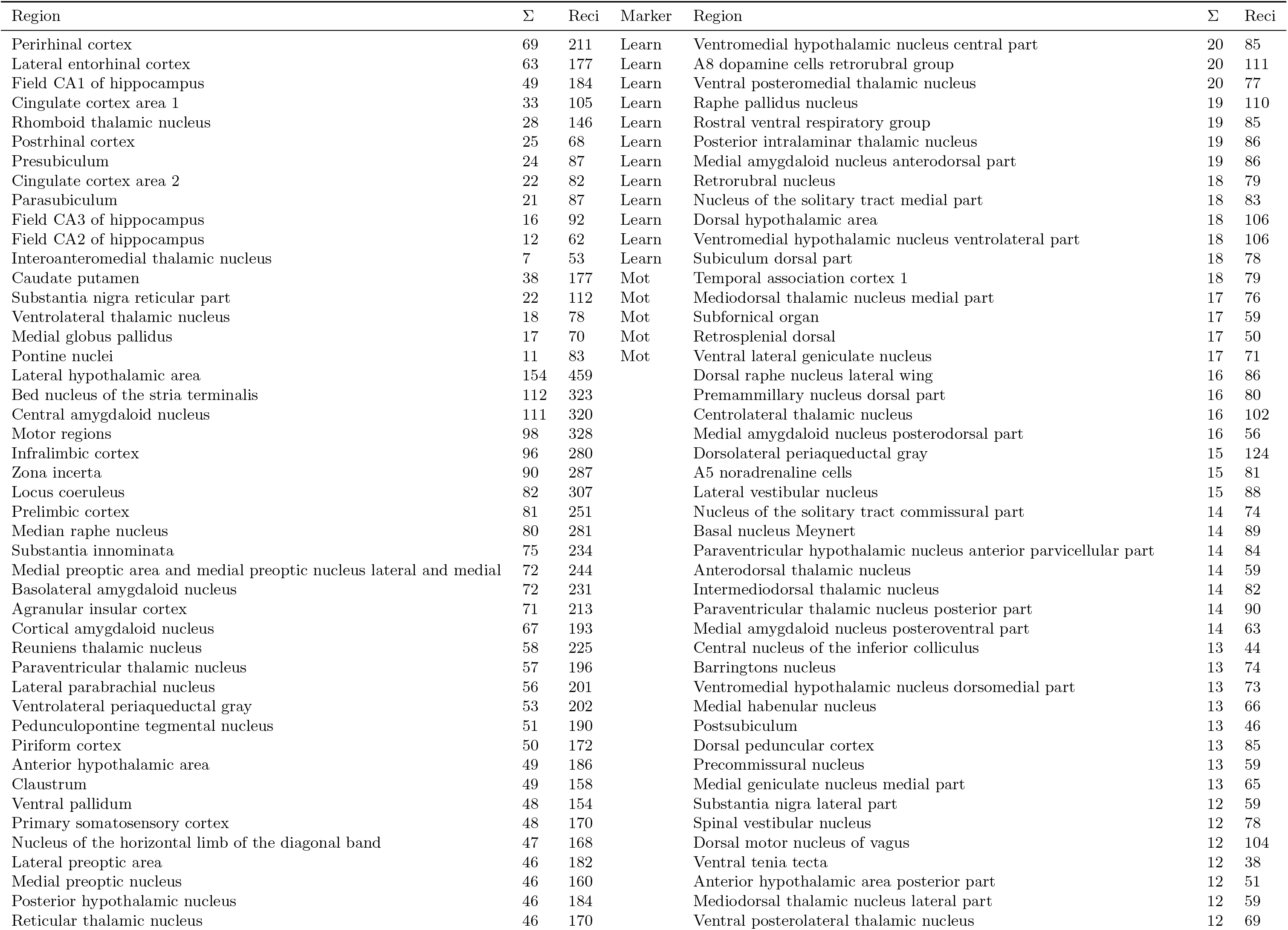
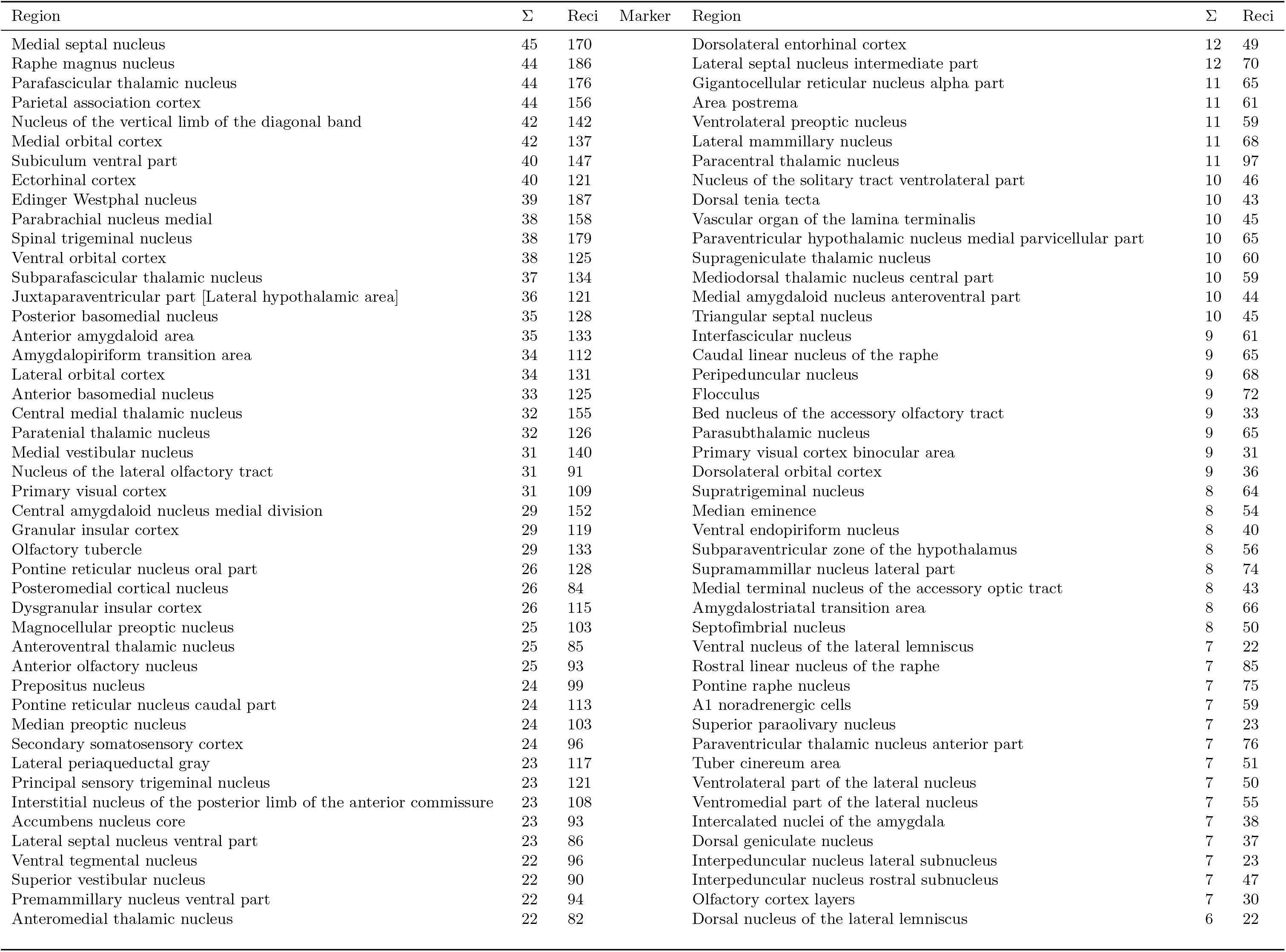
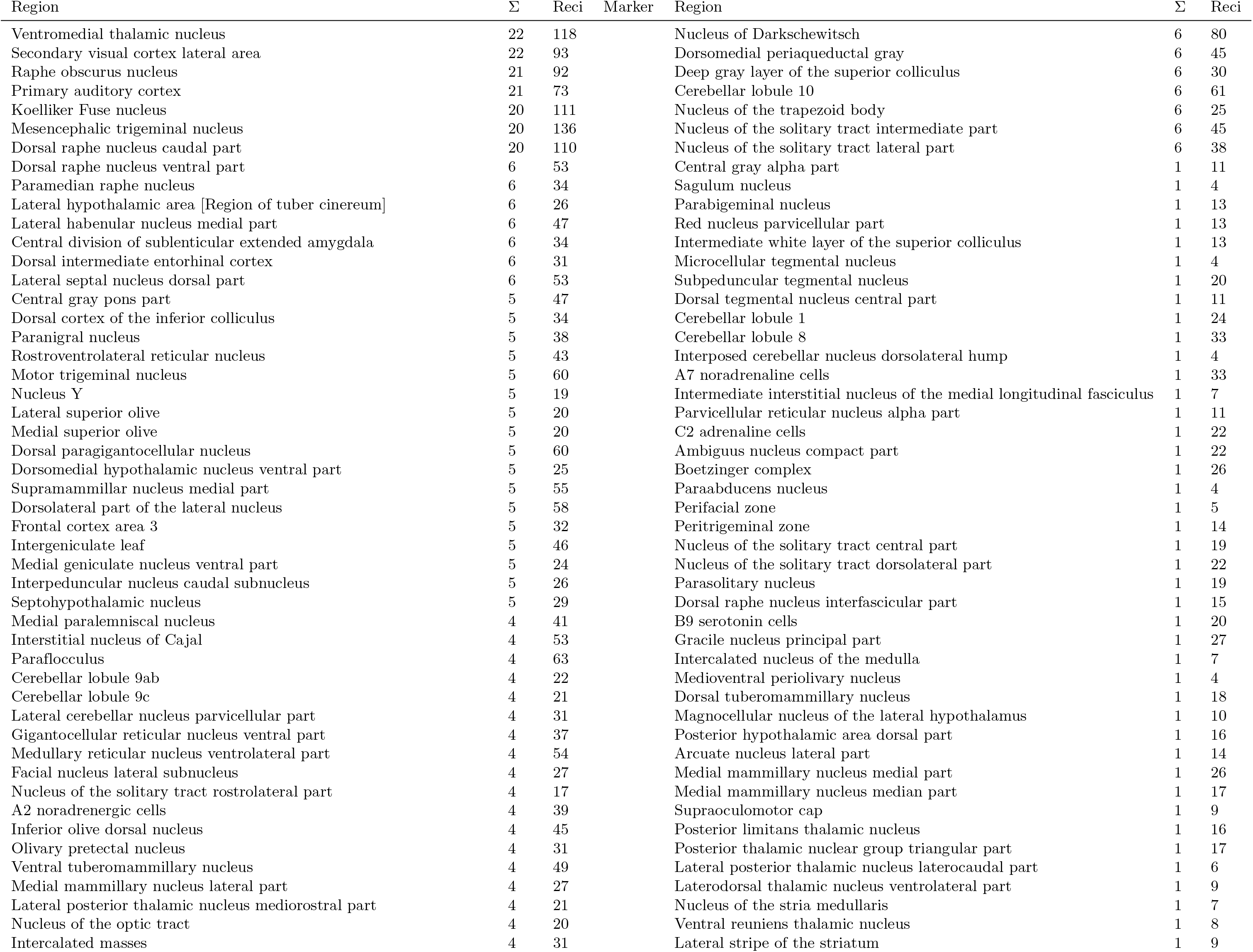
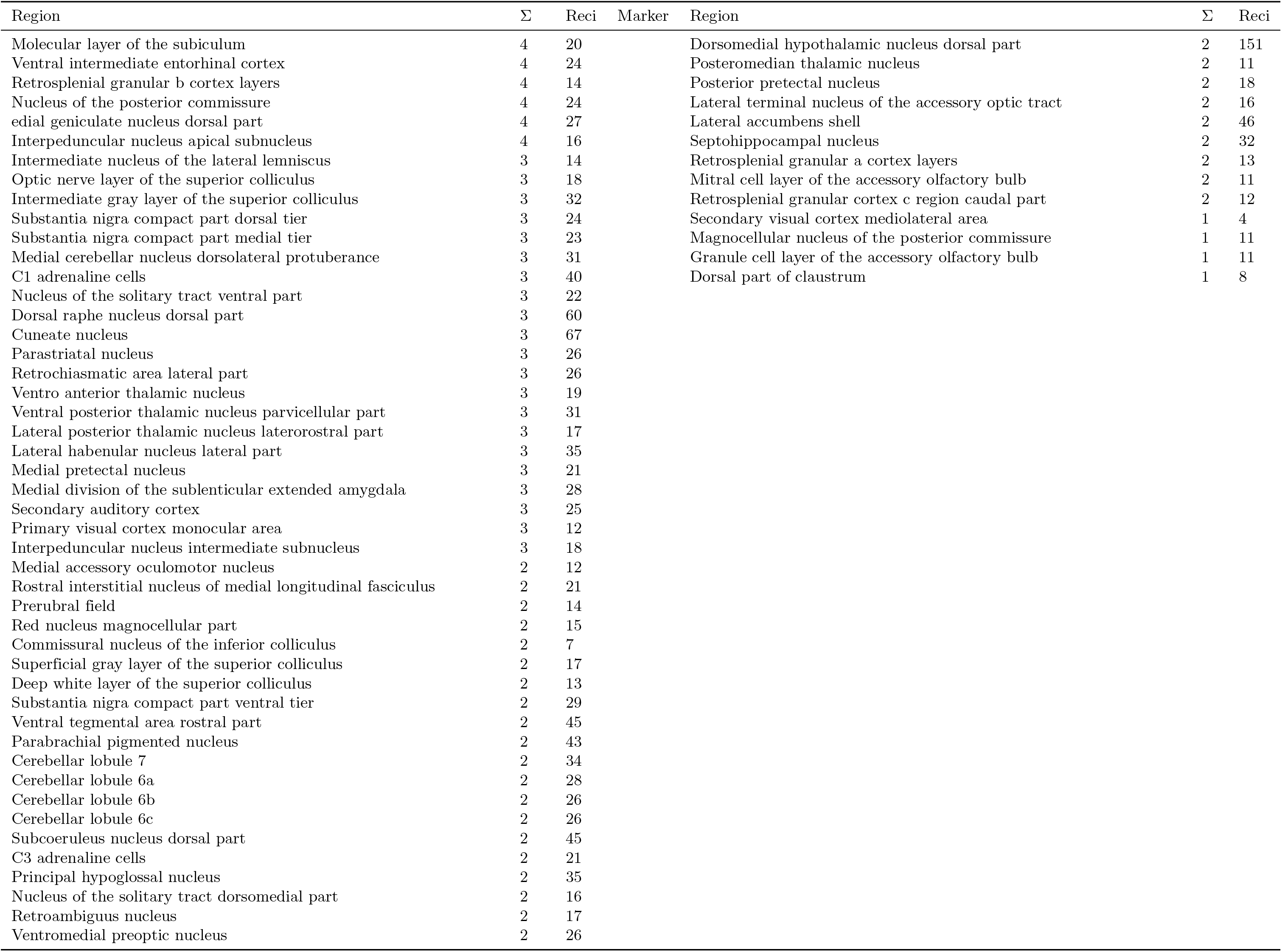
Overview of all connections of sMCAO lesioned regions. Connections of sMCAO lesioned regions, functionally defined regions and control regions without functional definitions.

**Table 7.**
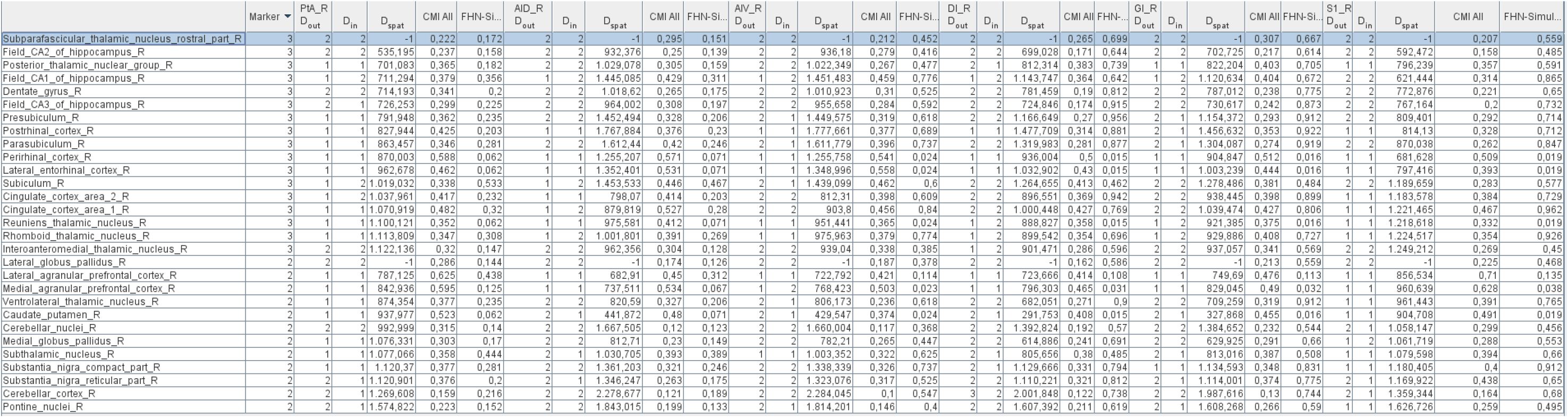
The similarity of functionally defined regions listed in the first column and the 6 lesioned regions of the dMCAO model. The marker column contains the code “3” for regions involved in learning and “2” for regions involved in motor behavior. The lesioned regions are shown in the columns PtA R, AID R, AIV R, DI R, GI R, S1 R. Dout and Din are indicating the minimal graph theoretical distance to link a functional region and a lesioned regions (columns 3, 8, 13, 18. . .). Dspat is the spatial distance between a pair of regions. CMIAll are the pairwise values for the similarity of connections of a functional region and a lesioned region. FHN are pairwise coactivations of a functional region and a lesioned region. The regions were sorted by their functional groups (marker column).

**Table 8.** Ranks of lesioned regions of the ICH experiment and the motor (Marker: 2) as well learning behavior (Marker: 3) groups. Sorting was performed for the coactivation matrix of a FHN simulation.

| Control connectome region | Marker | $CMI_{All}$ rank | Lesioned region | FHN rank | Lesioned region | Average | rank |
| --- | --- | --- | --- | --- | --- | --- | --- |
| Perirhinal cortex | 3 | 3 | Basolateral amygdaloid nucleus | 1 | Reticular thalamic nucleus | 27 |  |
| Lateral entorhinal cortex | 3 | 4 | Basolateral amygdaloid nucleus | 1 | Reticular thalamic nucleus | 9 |  |
| Mammillary body | 3 | 10 | Central amygdaloid nucleus | 1 | Reticular thalamic nucleus | 28 |  |
| Rhomboid thalamic nucleus | 3 | 5 | Paracentral thalamic nucleus | 2 | Ventromedial thalamic nucleus | 30 |  |
| Subiculum | 3 | 5 | Basolateral amygdaloid nucleus | 2 | Endopiriform system | 41 |  |
| Field CA1 of hippocampus | 3 | 7 | Basolateral amygdaloid nucleus | 2 | Central amygdaloid nucleus medial division | 61 |  |
| Dentate gyrus | 3 | 11 | Anteroventral thalamic nucleus | 2 | Medial globus pallidus | 102 |  |
| Cingulate cortex area 2 | 3 | 12 | Basal nucleus Meynert | 3 | Basal nucleus Meynert | 63 |  |
| Presubiculum | 3 | 7 | Anteroventral thalamic nucleus | 5 | Basal nucleus Meynert | 84 |  |
| Subparafascicular thalamic nucleus rostral part | 3 | 46 | Basolateral amygdaloid nucleus | 9 | Mediodorsal thalamic nucleus lateral part | 134 |  |
| Parasubiculum | 3 | 11 | Anteroventral thalamic nucleus | 11 | Interstitial nucleus of the posterior limb of the anterior commissure | 87 |  |
| Cingulate cortex area 1 | 3 | 11 | Caudate putamen | 11 | Interstitial nucleus of the posterior limb of the anterior commissure | 24 |  |
| Field CA3 of hippocampus | 3 | 15 | Endopiriform system | 11 | Paracentral thalamic nucleus | 104 |  |
| Field CA2 of hippocampus | 3 | 7 | Endopiriform system | 13 | Anteroventral thalamic nucleus | 147 |  |
| Postrhinal cortex | 3 | 16 | Anteroventral thalamic nucleus | 13 | Basal nucleus Meynert | 98 |  |
| Medial agranular prefrontal cortex | 2 | 10 | Caudate putamen | 1 | Reticular thalamic nucleus | 1 |  |
| Cerebellar cortex | 2 | 3 | Ventrobasal complex | 2 | Paracentral thalamic nucleus | 180 |  |
| Subthalamic nucleus | 2 | 2 | Caudate putamen | 3 | Endopiriform system | 55 |  |
| Lateral agranular prefrontal cortex | 2 | 4 | Caudate putamen | 3 | Caudate putamen | 4 |  |
| Ventrolateral thalamic nucleus | 2 | 1 | Ventromedial thalamic nucleus | 4 | Basal nucleus Meynert | 89 |  |
| Substantia nigra compact part | 2 | 17 | Basolateral amygdaloid nucleus | 7 | Interstitial nucleus of the posterior limb of the anterior commissure | 35 |  |
| Substantia nigra reticular part | 2 | 5 | Medial globus pallidus | 10 | Mediodorsal thalamic nucleus lateral part | 51 |  |
| Cerebellar nuclei | 2 | 21 | Ventrobasal complex | 31 | Ventro anterior thalamic nucleus | 120 |  |
| Pontine nuclei | 2 | 2 | Centrolateral thalamic nucleus | 32 | Anteroventral thalamic nucleus | 108 |  |
| Lateral hypothalamic area | 0 | 1 | Central amygdaloid nucleus | 1 | Reticular thalamic nucleus | 45 |  |
| Bed nucleus of the stria terminalis lateral division ventral part | 0 | 1 | Bed nucleus of the stria terminalis medial division | 3 | Mediodorsal thalamic nucleus lateral part | 100 |  |
| Bed nucleus of the stria terminalis intraamygdaloid division | 0 | 1 | Bed nucleus of the stria terminalis medial division | 5 | Mediodorsal thalamic nucleus lateral part | 143 |  |
| Dorsal striatum | 0 | 1 | Caudate putamen | 7 | Laterodorsal thalamic nucleus | 141 |  |
| Amygdaloid intramedullary gray | 0 | 1 | Amygdalostriatal transition area | 89 | Ventro anterior thalamic nucleus | 393 |  |

**Table 9.** Ranks of lesioned regions of the dMCAO experiment and the motor (Marker: 2) as well learning behavior (Marker: 3) groups. The articles which provides evidence for at least one of these functions are quoted in the column References. Sorting was performed for *CMI_All_* ranks. Marker 0 indicates regions of the control connectome. Those with large ranks were shown, only. The average rank column displays the average rank values of the local parameter computation.

| Control connectome region | Marker | References | $CM_{All}$ rank | Lesioned region | FHN rank | Lesioned region | Average rank |
| --- | --- | --- | --- | --- | --- | --- | --- |
| Perirhinal cortex | 3 | [96, 130, 143, 184] | 5 | Parietal association cortex | 13 | Agranular insular cortex dorsal part | 30 |
| Cingulate cortex area 1 | 3 | [72, 125, 188] | 6 | Agranular insular cortex dorsal part | 2 | Primary somatosensory cortex | 29 |
| Lateral entorhinal cortex | 3 | [101] | 11 | Agranular insular cortex ventral part | 13 | Agranular insular cortex dorsal part | 13 |
| Cingulate cortex area 2 | 3 | [72, 125, 188] | 11 | Parietal association cortex | 9 | Dysgranular insular cortex | 55 |
| Posterior thalamic nuclear group | 3 | [102] | 12 | Granular insular cortex | 90 | Dysgranular insular cortex | 97 |
| Parasubiculum | 3 | [97] | 15 | Agranular insular cortex dorsal part | 9 | Granular insular cortex | 88 |
| Anteroventral thalamic nucleus | 3 | [28, 32, 124] | 17 | Parietal association cortex | 213 | Dysgranular insular cortex | 119 |
| Interoanteromedial thalamic nucleus | 3 | [56, 82, 83, 170] | 17 | Granular insular cortex | 228 | Dysgranular insular cortex | 178 |
| Subiculum | 3 | [127, 136] | 18 | Agranular insular cortex ventral part | 23 | Agranular insular cortex ventral part | 46 |
| Dentate gyrus | 3 | [26, 146, 203] | 19 | Parietal association cortex | 47 | Dysgranular insular cortex | 89 |
| Subparafascicular thalamic nucleus rostral part | 3 | [147, 189] | 20 | Granular insular cortex | 207 | Dysgranular insular cortex | 132 |
| Postrhinal cortex | 3 | [4, 47, 202] | 20 | Parietal association cortex | 5 | Granular insular cortex | 100 |
| Field CA1 of hippocampus | 3 | [27, 103, 105] | 23 | Agranular insular cortex ventral part | 6 | Primary somatosensory cortex | 63 |
| Presubiculum | 3 | [78, 190] | 25 | Parietal association cortex | 7 | Dysgranular insular cortex | 81 |
| Anterodorsal thalamic nucleus | 3 | [42, 115] | 30 | Parietal association cortex | 265 | Dysgranular insular cortex | 153 |
| Field CA2 of hippocampus | 3 | [45, 75] | 33 | Agranular insular cortex ventral part | 215 | Dysgranular insular cortex | 161 |
| Rhomboid thalamic nucleus | 3 | [11, 64, 113] | 36 | Granular insular cortex | 4 | Primary somatosensory cortex | 40 |
| Reuniens thalamic nucleus | 3 | [11, 64, 113] | 40 | Agranular insular cortex dorsal part | 13 | Agranular insular cortex dorsal part | 10 |
| Field CA3 of hippocampus | 3 | [101, 152, 187] | 41 | Agranular insular cortex dorsal part | 14 | Dysgranular insular cortex | 109 |
| Mammillary body | 3 | [62, 131, 209, 210] | 63 | Parietal association cortex | 13 | Agranular insular cortex dorsal part | 17 |
| Lateral agranular prefrontal cortex | 2 | [93, 98] | 2 | Primary somatosensory cortex | 4 | Parietal association cortex | 6 |
| Medial agranular prefrontal cortex | 2 | [150, 151, 176] | 2 | Primary somatosensory cortex | 13 | Parietal association cortex | 5 |
| Caudate putamen | 2 | [36, 66] | 5 | Parietal association cortex | 13 | Agranular insular cortex dorsal part | 44 |
| Substantia nigra reticular part | 2 | [65, 86] | 9 | Primary somatosensory cortex | 49 | Dysgranular insular cortex | 53 |
| Lateral globus pallidus | 2 | [10, 141] | 11 | Parietal association cortex | 248 | Dysgranular insular cortex | 184 |
| Ventrolateral thalamic nucleus | 2 | [51] | 15 | Primary somatosensory cortex | 14 | Granular insular cortex | 80 |
| Cerebellar nuclei | 2 | [24, 38, 204] | 25 | Parietal association cortex | 280 | Dysgranular insular cortex | 118 |
| Substantia nigra compact part | 2 | [61, 63] | 27 | Primary somatosensory cortex | 5 | Primary somatosensory cortex | 31 |
| Medial globus pallidus | 2 | [46, 52, 61] | 27 | Parietal association cortex | 207 | Dysgranular insular cortex | 85 |
| Pontine nuclei | 2 | [30, 31, 76, 94, 137] | 29 | Granular insular cortex | 256 | Dysgranular insular cortex | 108 |
| Subthalamic nucleus | 2 | [16, 39, 141] | 33 | Primary somatosensory cortex | 24 | Primary somatosensory cortex | 54 |
| Cerebellar cortex | 2 | [34, 122, 186, 204] | 91 | Primary somatosensory cortex | 37 | Granular insular cortex | 182 |
| Agranular insular cortex posterior part | 0 |  | 1 | Agranular insular cortex ventral part | 7 | Agranular insular cortex ventral part | 47 |
| Secondary somatosensory cortex | 0 |  | 1 | Primary somatosensory cortex | 2 | Granular insular cortex | 58 |
| Primary visual cortex | 0 |  | 1 | Parietal association cortex | 7 | Granular insular cortex | 67 |
| Anterior basolateral nucleus | 0 |  | 2 | Agranular insular cortex dorsal part | 8 | Primary somatosensory cortex | 57 |
| Frontal cortex area 3 | 0 |  | 3 | Agranular insular cortex ventral part | 225 | Dysgranular insular cortex | 197 |
| Secondary visual cortex lateral area | 0 |  | 3 | Parietal association cortex | 244 | Dysgranular insular cortex | 133 |
| Lateral orbital cortex | 0 |  | 3 | Agranular insular cortex dorsal part | 14 | Parietal association cortex | 34 |
| Ventral orbital cortex | 0 |  | 3 | Agranular insular cortex dorsal part | 7 | Parietal association cortex | 21 |
| Dorsal striatum | 0 |  | 4 | Parietal association cortex | 9 | Granular insular cortex | 143 |
| Temporal association cortex 1 | 0 |  | 4 | Parietal association cortex | 233 | Dysgranular insular cortex | 154 |
| Dorsal peduncular cortex | 0 |  | 4 | Agranular insular cortex ventral part | 254 | Dysgranular insular cortex | 159 |
| Dorsolateral orbital cortex | 0 |  | 4 | Agranular insular cortex dorsal part | 234 | Dysgranular insular cortex | 172 |
| Medial orbital cortex | 0 |  | 4 | Agranular insular cortex dorsal part | 16 | Agranular insular cortex ventral part | 22 |

**Table 10.** Ranks of lesioned regions of the dMCAO experiment and the motor (Marker: 2) as well learning behavior (Marker: 3) groups. Sorting was performed for the FHN rank computed with the coactivation matrix of a FHN simulation. The average rank column displays the average rank values of the local parameter computation.

| Control connectome region | Marker | $CM I_{All}$ rank | Lesioned region | FHN rank | Lesioned region | Average rank |
| --- | --- | --- | --- | --- | --- | --- |
| Cingulate cortex area 1 | 3 | 6 | Agranular insular cortex dorsal part | 2 | Primary somatosensory cortex | 29 |
| Rhomboid thalamic nucleus | 3 | 36 | Granular insular cortex | 4 | Primary somatosensory cortex | 40 |
| Postrhinal cortex | 3 | 20 | Parietal association cortex | 5 | Granular insular cortex | 100 |
| Field CA1 of hippocampus | 3 | 23 | Agranular insular cortex ventral part | 6 | Primary somatosensory cortex | 63 |
| Presubiculum | 3 | 25 | Parietal association cortex | 7 | Dysgranular insular cortex | 81 |
| Cingulate cortex area 2 | 3 | 11 | Parietal association cortex | 9 | Dysgranular insular cortex | 55 |
| Parasubiculum | 3 | 15 | Agranular insular cortex dorsal part | 9 | Granular insular cortex | 88 |
| Perirhinal cortex | 3 | 5 | Parietal association cortex | 13 | Agranular insular cortex dorsal part | 30 |
| Lateral entorhinal cortex | 3 | 11 | Agranular insular cortex ventral part | 13 | Agranular insular cortex dorsal part | 13 |
| Reuniens thalamic nucleus | 3 | 40 | Agranular insular cortex dorsal part | 13 | Agranular insular cortex dorsal part | 10 |
| Mammillary body | 3 | 63 | Parietal association cortex | 13 | Agranular insular cortex dorsal part | 17 |
| Field CA3 of hippocampus | 3 | 41 | Agranular insular cortex dorsal part | 14 | Dysgranular insular cortex | 109 |
| Subiculum | 3 | 18 | Agranular insular cortex ventral part | 23 | Agranular insular cortex ventral part | 46 |
| Dentate gyrus | 3 | 19 | Parietal association cortex | 47 | Dysgranular insular cortex | 89 |
| Posterior thalamic nuclear group | 3 | 12 | Granular insular cortex | 90 | Dysgranular insular cortex | 97 |
| Subparafascicular thalamic nucleus rostral part | 3 | 20 | Granular insular cortex | 207 | Dysgranular insular cortex | 132 |
| Anteroventral thalamic nucleus | 3 | 17 | Parietal association cortex | 213 | Dysgranular insular cortex | 119 |
| Field CA2 of hippocampus | 3 | 33 | Agranular insular cortex ventral part | 215 | Dysgranular insular cortex | 161 |
| Interanteromedial thalamic nucleus | 3 | 17 | Granular insular cortex | 228 | Dysgranular insular cortex | 178 |
| Anterodorsal thalamic nucleus | 3 | 30 | Parietal association cortex | 265 | Dysgranular insular cortex | 153 |
| Lateral agranular prefrontal cortex | 2 | 2 | Primary somatosensory cortex | 4 | Parietal association cortex | 6 |
| Substantia nigra compact part | 2 | 27 | Primary somatosensory cortex | 5 | Primary somatosensory cortex | 31 |
| Medial agranular prefrontal cortex | 2 | 2 | Primary somatosensory cortex | 13 | Parietal association cortex | 5 |
| Caudate putamen | 2 | 5 | Parietal association cortex | 13 | Agranular insular cortex dorsal part | 44 |
| Ventrolateral thalamic nucleus | 2 | 15 | Primary somatosensory cortex | 14 | Granular insular cortex | 80 |
| Subthalamic nucleus | 2 | 33 | Primary somatosensory cortex | 24 | Primary somatosensory cortex | 54 |
| Cerebellar cortex | 2 | 91 | Primary somatosensory cortex | 37 | Granular insular cortex | 182 |
| Substantia nigra reticular part | 2 | 9 | Primary somatosensory cortex | 49 | Dysgranular insular cortex | 53 |
| Medial globus pallidus | 2 | 27 | Parietal association cortex | 207 | Dysgranular insular cortex | 85 |
| Lateral globus pallidus | 2 | 11 | Parietal association cortex | 248 | Dysgranular insular cortex | 184 |
| Pontine nuclei | 2 | 29 | Granular insular cortex | 256 | Dysgranular insular cortex | 108 |
| Cerebellar nuclei | 2 | 25 | Parietal association cortex | 280 | Dysgranular insular cortex | 118 |
| Secondary somatosensory cortex | 0 | 1 | Primary somatosensory cortex | 2 | Granular insular cortex | 58 |
| Agranular insular cortex posterior part | 0 | 1 | Agranular insular cortex ventral part | 7 | Agranular insular cortex ventral part | 47 |
| Primary visual cortex | 0 | 1 | Parietal association cortex | 7 | Granular insular cortex | 67 |
| Ventral orbital cortex | 0 | 3 | Agranular insular cortex dorsal part | 7 | Parietal association cortex | 21 |
| Anterior basolateral nucleus | 0 | 2 | Agranular insular cortex dorsal part | 8 | Primary somatosensory cortex | 57 |
| Dorsal striatum | 0 | 4 | Parietal association cortex | 9 | Granular insular cortex | 143 |
| Lateral orbital cortex | 0 | 3 | Agranular insular cortex dorsal part | 14 | Parietal association cortex | 34 |
| Medial orbital cortex | 0 | 4 | Agranular insular cortex dorsal part | 16 | Agranular insular cortex ventral part | 22 |
| Frontal cortex area 3 | 0 | 3 | Agranular insular cortex ventral part | 225 | Dysgranular insular cortex | 197 |
| Temporal association cortex 1 | 0 | 4 | Parietal association cortex | 233 | Dysgranular insular cortex | 154 |
| Dorsolateral orbital cortex | 0 | 4 | Agranular insular cortex dorsal part | 234 | Dysgranular insular cortex | 172 |
| Secondary visual cortex lateral area | 0 | 3 | Parietal association cortex | 244 | Dysgranular insular cortex | 133 |
| Dorsal peduncular cortex | 0 | 4 | Agranular insular cortex ventral part | 254 | Dysgranular insular cortex | 159 |

**Table 11.**
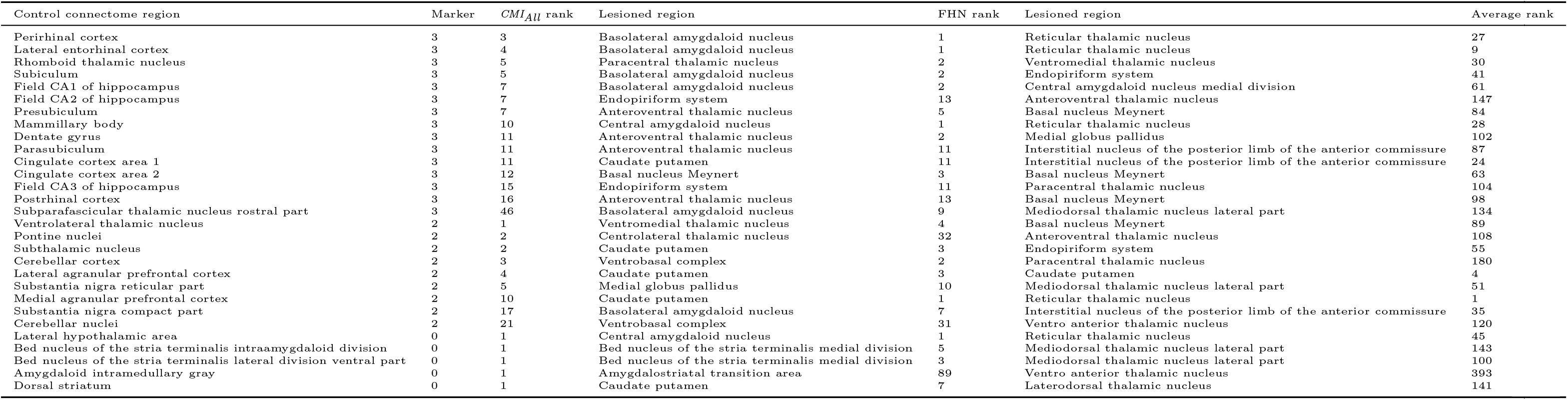
Ranks of lesioned regions of the ICH experiment and the motor (Marker: 2) as well learning behavior (Marker: 3) groups. Sorting was performed for the *CMI_All_* rank.

**Table 12.** Similarities of lesioned regions with regard to motor regions or learning regions or non-lesioned regions. Larger similarity values indicate a stronger similarity. The maximum values for each experimental group and method were highlighted.

| Method | Comparison | dMCAO | ICH | sMCAO |
| --- | --- | --- | --- | --- |
| *CMI<sub>All</sub>* | Lesioned vs motor | 0.328 | **0.271** | 0.251 |
|  | Lesioned vs learning | **0.352** | 0.256 | **0.302** |
|  | Lesioned vs non-lesioned | 0.119 | 0.103 | 0.104 |
| FHN-model | Lesioned vs motor | 0.359 | **0.399** | 0.425 |
|  | Lesioned vs learning | **0.432** | 0.384 | **0.471** |
|  | Lesioned vs non-lesioned | 0.278 | 0.343 | 0.355 |

### S2 Figures

**Figure 1.**
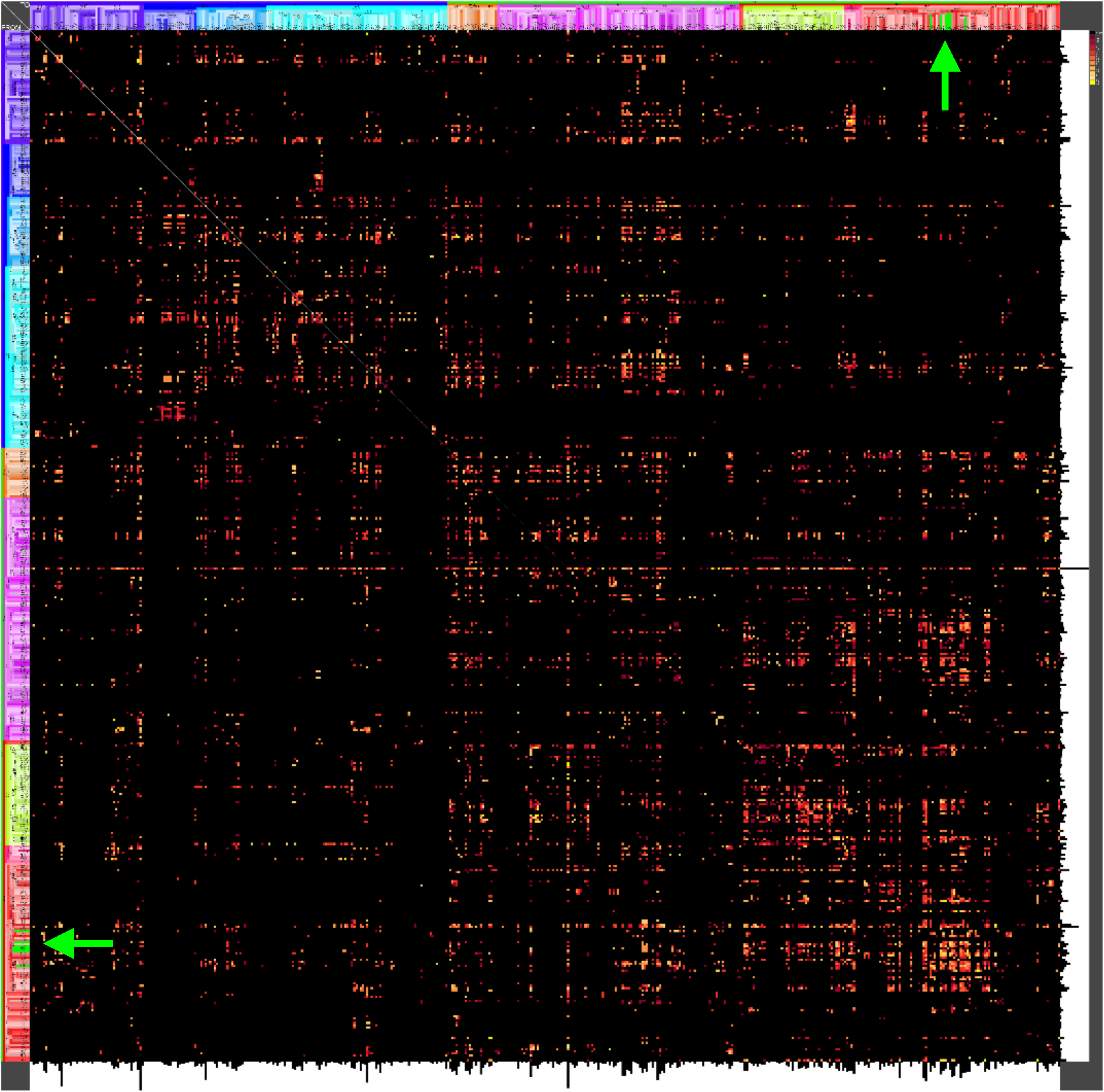
The directed and weighted ipsilateral adjacency matrix with 6 lesioned region of the dMCAO model (green). Cortical non-lesioned (red half-tones) and lesioned regions (arrow, green) are located in the upper right, resp., lower left part of the matrix.

**Figure 2.**
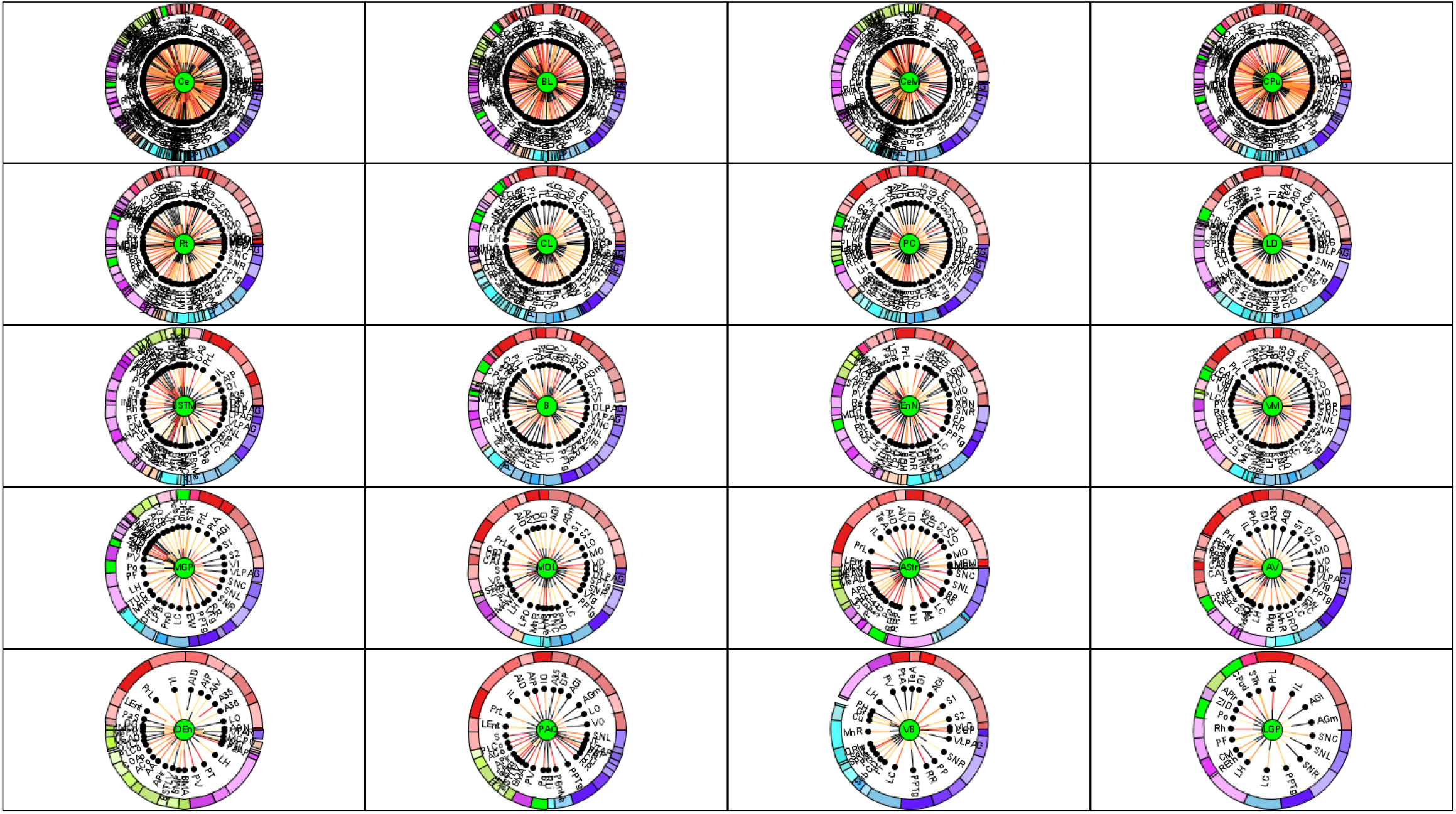
A complete MCRD visualization of ICH lesioned regions and their relations to control connectome regions. Distance from center region (lesioned region) is the average rank of local network parameters filtered for the 25% of lowest ranks (largest importance for the network). The lesioned regions were sorted with regard to the number of filtered and connected regions. All other lesioned regions have to large average ranks or too few connections and they are not displayed.

**Figure 3.**
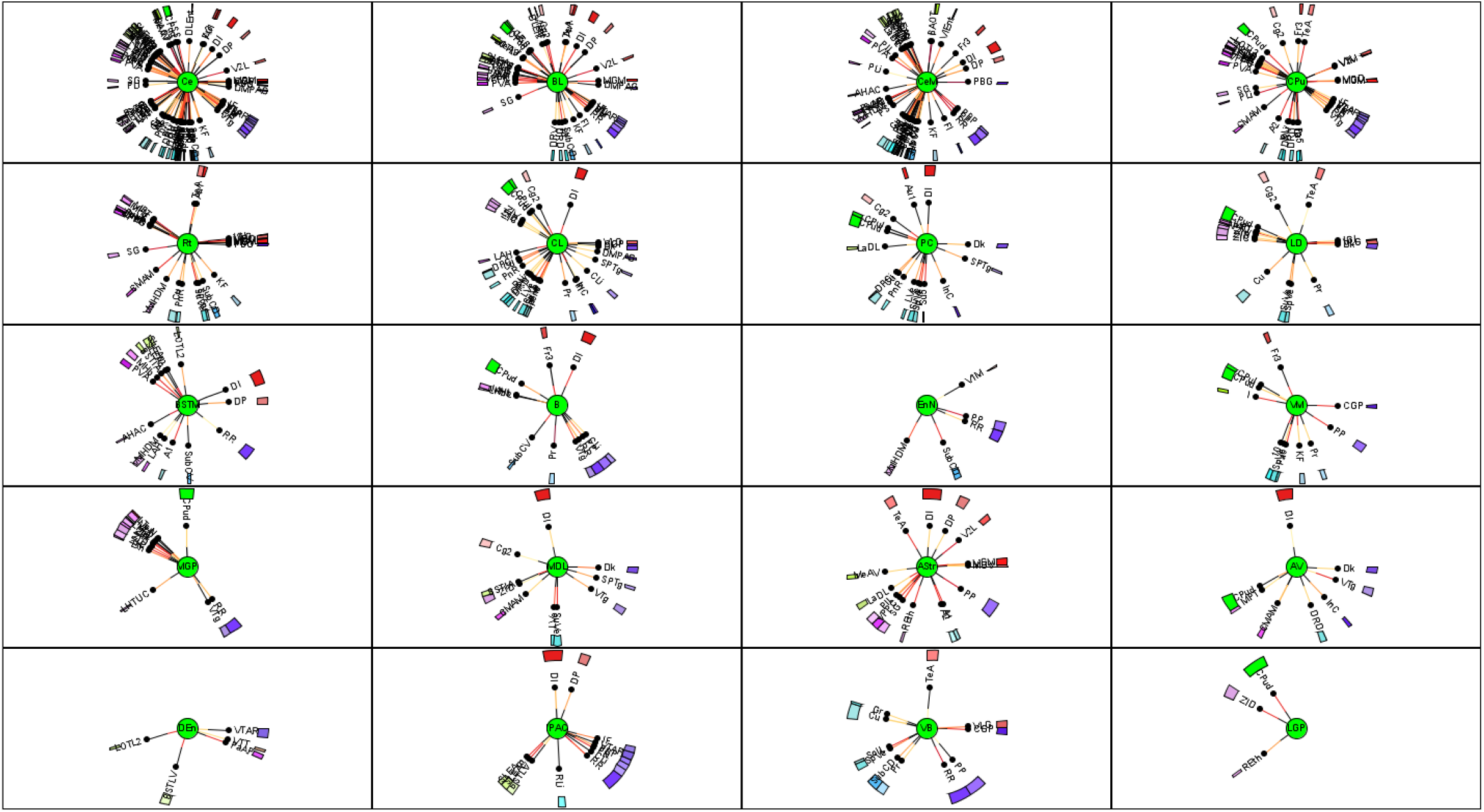
A filtered MCRD visualization of ICH lesioned regions and their relations to control connectome regions. Distance from center region (lesioned region) is the average rank of local network parameters filtered for the 80% of lowest ranks (largest importance for the network). The lesioned regions were sorted with regard to the number of filtered and connected regions.

**Figure 4.**
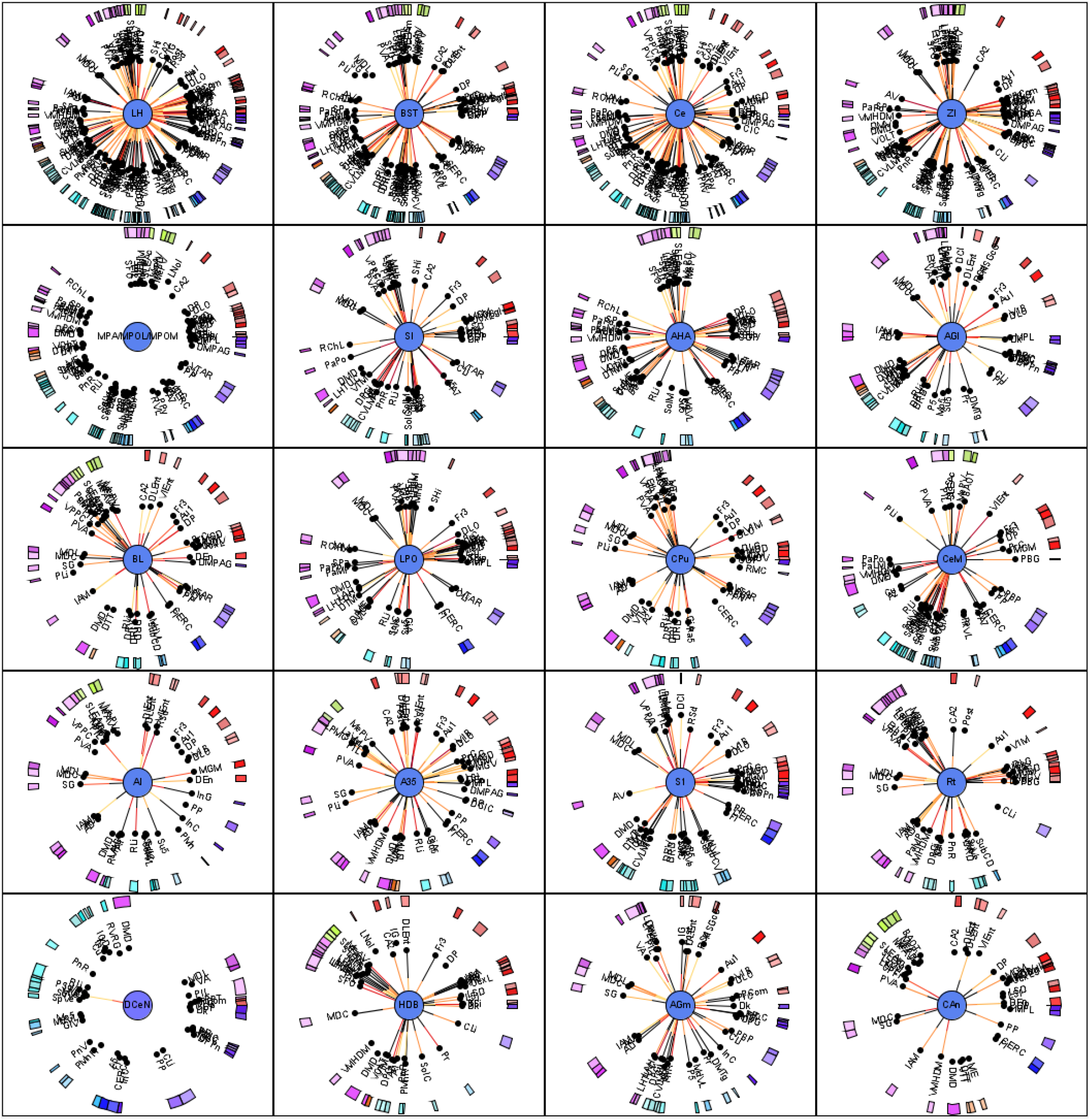
Part 1 of the sorted MCRD diagrams of the sMCAO model. Filtering conditions are the same as described in Figure 11.

**Figure 5.**
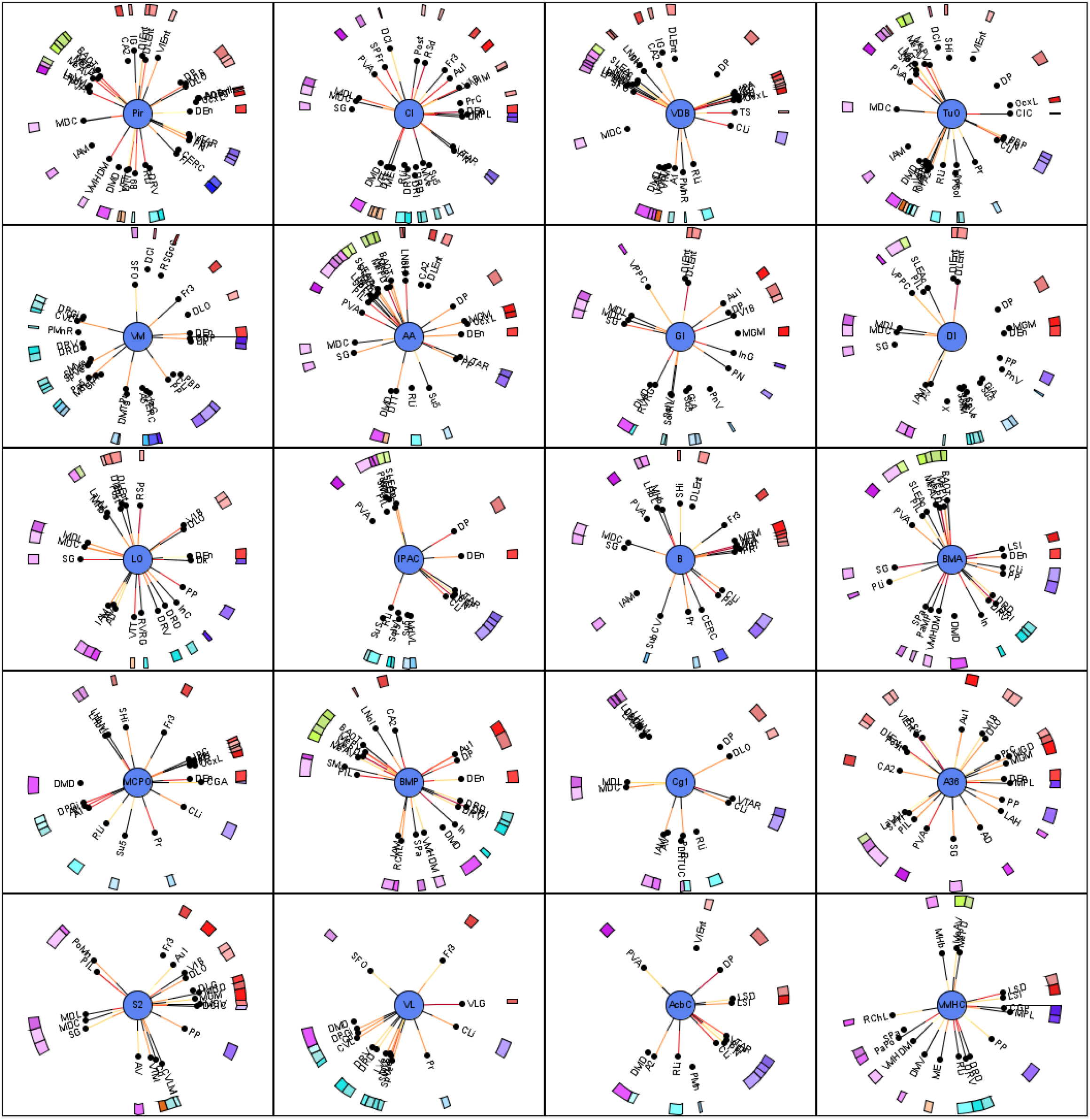
Part 2 of the sorted MCRD diagrams of the sMCAO model. Filtering conditions are the same as described in Figure 11.

**Figure 6.**
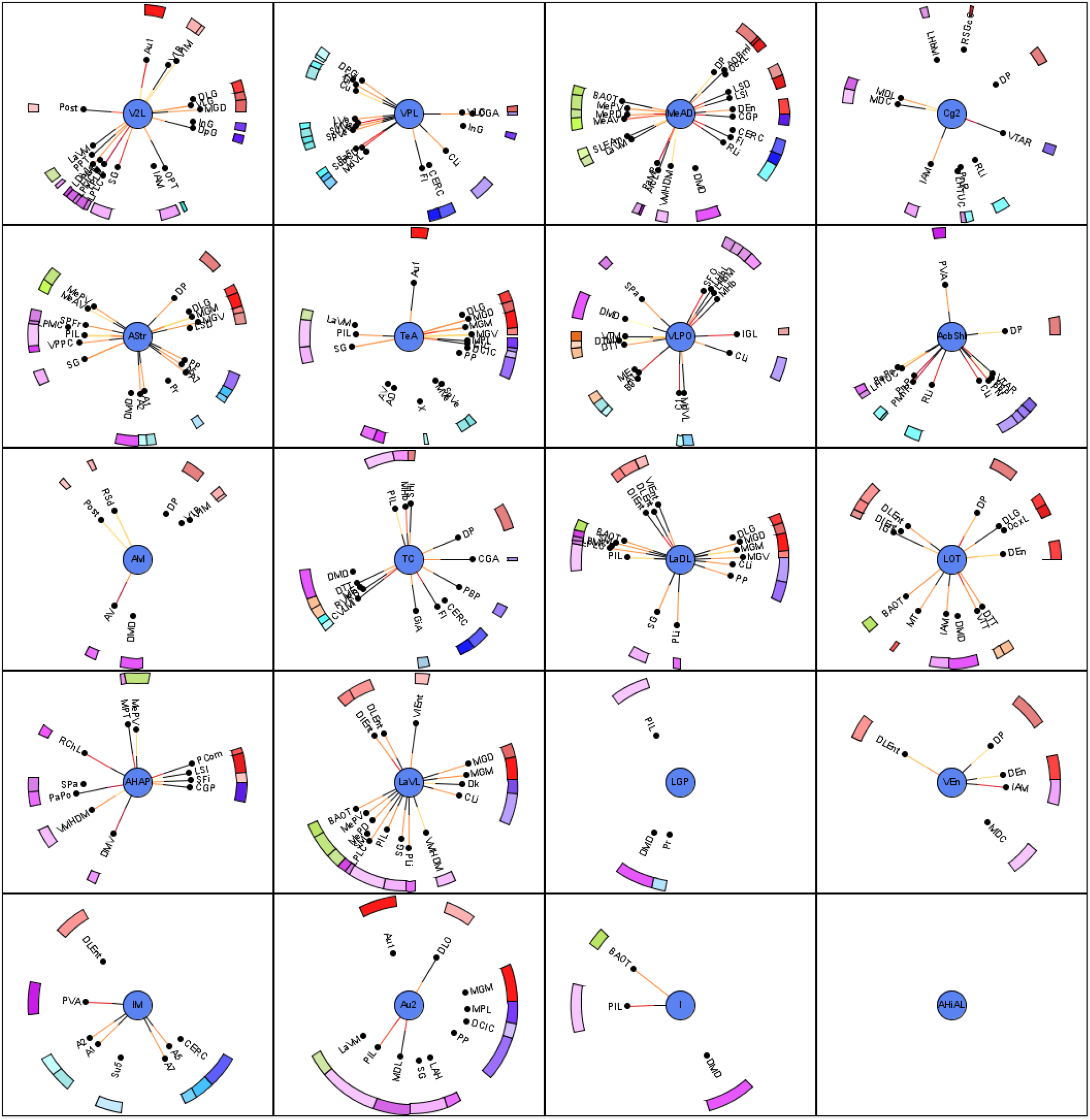
Part 3 of the sorted MCRD diagrams of the sMCAO model. Filtering conditions are the same as described in Figure 11.

